# Mechanisms used for cDNA synthesis and site-specific integration of RNA into DNA genomes by a reverse transcriptase-Cas1 fusion protein

**DOI:** 10.1101/2023.09.01.555893

**Authors:** Georg Mohr, Jun Yao, Seung Kuk Park, Laura M. Markham, Alan M. Lambowitz

**Affiliations:** Departments of Molecular Biosciences and Oncology University of Texas at Austin Austin TX, 78712

**Keywords:** genome engineering, prime editing, site-specific DNA integration, spacer acquisition

## Abstract

Reverse transcriptase-Cas1 (RT-Cas1) fusion proteins found in some CRISPR systems enable spacer acquisition from both RNA and DNA, but the mechanism of RNA spacer acquisition has remained unclear. Here, we found *Marinomonas mediterranea* RT-Cas1/Cas2 adds short 3’-DNA (dN) tails to RNA protospacers enabling their direct integration into CRISPR arrays as 3’-dN-RNA/cDNA duplexes or 3’-dN-RNAs at rates comparable to similarly configured DNAs. Reverse transcription of RNA protospacers occurs by multiple mechanisms, including recently described *de novo* initiation, protein priming with any dNTP, and use of short exogenous or synthesized DNA oligomer primers, enabling synthesis of cDNAs from diverse RNAs without fixed sequence requirements. The integration of 3’-dN-RNAs or single-stranded (ss) DNAs is favored over duplexes at higher protospacer concentrations, potentially relevant to spacer acquisition from abundant pathogen RNAs or ssDNA fragments generated by phage-defense nucleases. Our findings reveal novel mechanisms for site-specifically integrating RNA into DNA genomes with potential biotechnological applications.

## INTRODUCTION

Bacteria face incessant attacks by viruses and genomic parasites and have evolved defense systems to combat these threats. Among the most widely studied are CRISPR (clustered regularly interspaced short palindromic repeats)-Cas systems found in diverse bacteria and archaea.^1^ CRISPR-Cas systems typically include an RNA-guided nuclease (effector) complex, a CRISPR repeat locus that accepts snippets of nucleic acids (protospacers) derived from threats, enzymes (Cas6 or Cas4) that processes transcribed spacers into RNA guides for nuclease effector complexes, and a machinery (Cas1/Cas2) that site-specifically integrates new spacers from invading pathogens into CRISPR arrays.^2–8^ Six types of CRISPR systems (Type I to VI) each with multiple subclasses have been distinguished.^1^ Type III systems differ from the others in their ability to cleave both DNA and RNA in a transcriptionally coupled reaction.^9–11^ Additionally, some Type III CRISPR systems have an associated reverse transcriptase (RT), either co-expressed with Cas1/Cas2 or present as a reverse transcriptase-Cas1 (RT-Cas1) fusion protein, some of which also have an N-terminal Cas6 domain that processes guide RNAs for incorporation into effector complexes.^12–16^ Most CRISPR-associated RTs are closely related to the RTs encoded by mobile group II introns, prolific bacterial retrotransposons whose dissociated RTs have evolved in different bacteria to perform a variety of cellular functions.^17–22^ Four RT-Cas1 proteins have been shown to site-specifically integrate RNA as well as DNA into CRISPR arrays *in vivo*,^12,23–25^ but the mechanisms by which RNA protospacers are reverse transcribed and integrated into CRISPR arrays have remained unclear.

Although the six types of CRISPR systems vary in the mechanisms used for processing guide RNAs and destroying invading nucleic acids, all use the same Cas1/Cas2-mediated cleavage-ligation (transesterification) mechanism to site-specifically integrate double-strand (ds) DNA protospacers into CRISPR arrays.^26–30^ Structural studies have shown that Cas1 and Cas2 form a hexameric complex that binds a dsDNA protospacer across its length with overhanging or splayed single-stranded 3’ ends of opposite strands inserted in one or both active sites of Cas1 proteins on opposite sides of the complex.^25,31–37^ Cas1/Cas2 complexes from different organisms have structural variations that favor different length spacers and splay open different length single-stranded 3’ ends.^30,38^ A single-molecule study of spacer acquisition by an *Enterococcus faecalis* Type IIA CRISPR system indicated that the Cas1/Cas2 complex remains stably bound to the integrated spacer until it is dislodged by transcription-coupled DNA repair, which fills in and seals single-stranded gaps to fully integrate the newly acquired spacer into the host genome.^39^ This process enables integration of protospacers into CRISPR arrays without introducing deleterious double-strand breaks in bacterial chromosomal DNA.

Biochemical analyses of the mechanism by which RT-Cas1 proteins acquire spacers from RNA have been sparse. The 4 RT-Cas1 fusion proteins that acquire spacers from RNA are comprised of an RT domain corresponding to the fingers and palm of retroviral RTs, but fused directly to Cas1 rather than a thumb domain as in canonical RTs (Figure 1A). The RT domains of these proteins contain 7 conserved sequences blocks (RT1-7) found in all RTs plus an N-terminal extension (NTE) with an RT0 loop and two expanded regions (RT2a and RT3a) between the conserved RT sequence blocks (Figure 1A). These additional regions are absent in retroviral RTs, but structurally conserved and functionally important in group II intron and other bacterial RTs, as well as in LINE-1 and other eukaryotic non-LTR-retrotransposon RTs (collectively termed non-LTR-retroelement RTs).^40^ The *Marinomonas mediterranea* (Mm) RT-Cas1 protein, associated with a Type IIIA CRISPR system, was shown to function in complex with Cas2 to site-specifically integrate DNA and RNA protospacers into CRISPR arrays *in vitro* and to have an active Cas6 domain that functions in CRISPR RNA processing and whose interaction with the RT domain is required for RT activity.^12,14^ A cryo-EM structure showed that a closely related *Thiomicrospira* Type III Cas6-RT-Cas1/Cas2 forms a hexameric complex similar to Cas1/Cas2 proteins that acquire spacers from DNA, but with the Cas6 and RT domains interacting with each other to form flexibly attached lobes and with some structural differences in regions that function in protospacer binding.^14,25^ The *Fusicatenibacter saccharivorans* (Fs) and *Vibrio vulnificus* (Vv) RT-Cas1 proteins, both associated with Type IIID CRISPR systems, were shown to acquire RNA-derived spacers *in vivo,* but have not been investigated biochemically.^23,24^ Notably, Mm RT-Cas1/Cas2 could acquire both RNA and DNA protospacers in its native host, but only DNA protospacers in *E. coli*, suggesting that host-specific factors contribute to RNA protospacer acquisition *in vivo.*^12^

**Figure 1.**
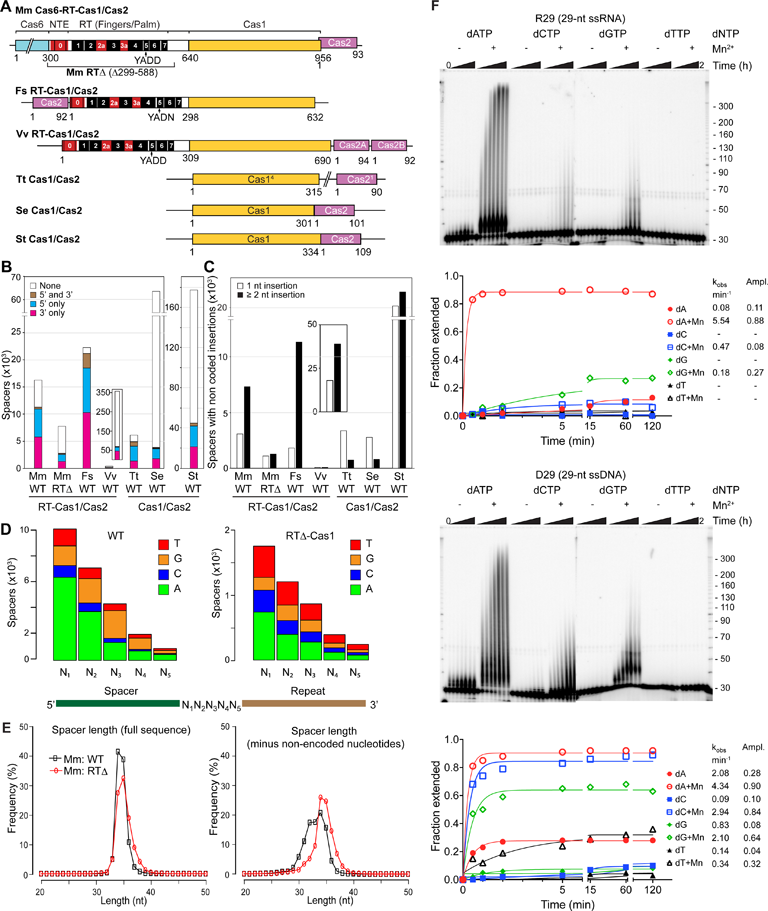
RT-Cas1/Cas2 terminal transferase activity adds non-coded nucleotides to the 3’ end RNA protospacers. (A) Schematics of RT-Cas1/Cas2 and Cas1/Cas2 proteins. Protein regions: RT1-7, conserved sequence blocks present in all RTs (black); NTE/RT0, RT2a and RT3a, structurally conserved regions of non-LTR-retroelement RTs that are absent in retroviral RTs (red); Abbreviations: Fs, *Fusicatenibacter saccharivorans*; Mm, *Marinomonas mediterranea*; Vv, *Vibrio vulnificus*; Tt, *Thermus thermophilus*; Se, *Staphylococcus epidermidis*; St, *Streptococcus thermophilus*. Stacked bar graphs comparing the numbers of 5’-leader proximal spacers with non-coded nucleotides at their 5’ end only (blue), 3’ end only (red), both ends (brown), or none (white) acquired *in vivo* by RT-Cas1/Cas2 or Cas1/Cas2 proteins associated with different Type III CRISPR systems. The analysis was done using the datasets listed in Methods for different unique spacer sequences that could be mapped to the host bacterium or phage genomes. The *V. vulnificus* and *S. thermophilus* data are plotted on different scales. (B) Stacked bar graphs comparing the numbers of spacers acquired by the RT-Cas1/Cas2 or Cas/Cas2 proteins in panel B with a single non-coded nucleotide (white bars) or 22 non-coded nucleotides (black bars) at either spacer-repeat junction. The analysis was done for the spacers analyzed in panel B, excluding small numbers of spacers that have non-coded nucleotides at both spacer-repeat junctions. (C) Stacked bar graphs comparing the frequencies of different non-coded nucleotides located at the 3’ end of spacers acquired by Mm WT RT-Cas1/Cas2 or RT1′-Cas1/Cas2. The analysis was done for the spacers analyzed in panel B that had non-coded nucleotides at one end only, putatively the 3’ end used for terminal transferase addition of non-coded nucleotides. (D) Length distribution of unique spacers acquired by the Mm WT RT-Cas1 (black) or RT1′-Cas1 (red). The analysis was done for the spacers that mapped to the *M. mediterranea* strain MMB1 genome. The plot on the left shows the length distribution of spacers defined as the sequence between two repeats, and the plot on the right shows the length distribution of the same spacers after removing non-coded nucleotides. (E) Terminal transferase assay done as described in Methods by incubating Mm RT-Cas1/Cas2 with a 5’-^32^P labeled 29-nt RNA (R29, top) or DNA (D29, bottom) oligonucleotides having the same nucleotide sequence with 1 mM of the indicated dNTP in reaction medium containing 10 mM MgCl_2_ with or without 1 mM MnCl_2_ at 37°C for times up to 2 h. The products were analyzed in a denaturing 6% polyacrylamide gel, which was dried and quantified with a phosphorimager. The hatch marks to the right of the gel indicate the positions of 5’-labeled size markers (10-nt ssDNA ladder; Invitrogen) run on the same gel. The plots below the gels show the fraction of labeled product as a function of time with the data fit to a single exponential equation to obtain values for k_obs_ and amplitude. R^2^ values for the curve fits are listed in the Supplemental data file. A minus sign (-) indicates no detectable extension of the oligonucleotide substrate.

Here, we focused on mechanisms used for reverse transcription and RNA protospacer integration by Mm RT-Cas1/Cas2. Zabrady *et al.* reported that Mm RT-Cas1/Cas2 could initiate reverse transcription *de novo* at C residues by using a Mn^2+^-dependent primase activity with a strong preference for initiating at CC sequences.^41^ Here, we found that Mm RT-Cas1/Cas2 could also initiate reverse transcription by protein priming with any dNTP and by using short exogenous and likely synthesized DNA oligonucleotide primers with no fixed sequence requirements, as expected for an enzyme whose biological function is to acquire spacers from diverse RNAs. Zabrady *et al*. also reported that Mm RT-Cas1/Cas2 could add short 3’-DNA (3’-dN) extensions to RNA protospacers but that the efficient integration of RNA protospacers into a CRISPR array required synthesis of a complementary cDNA that is a preferred substrate for the Cas1/Cas2 DNA integrase.^41^ Here, we found that Mm RT-Cas1/Cas2 could directly integrate 3’-dN-RNA/cDNA duplexes and single-stranded 3’-dN-RNAs into a CRISPR array in the absence of cDNA synthesis at rates comparable to similarly configured DNA protospacers and with bioinformatic analysis showing that spacers acquired from 3’-dN-tailed RNAs comprise a high proportion of those acquired by Mm and other RT-Cas1/Cas2 proteins *in vivo*.

## RESULTS

### Spacers acquired by Mm RT-Cas1/Cas1 *in vivo* have higher numbers of non-coded nucleotides at spacer-repeat junctions

We wondered if spacers acquired from RNA might have distinctive features that could provide clues about their acquisition mechanism. To identify such features, we compared the sequences of newly acquired spacers (*i.e.*, those closest to the 5’-leader sequence) for three RT-Cas1 proteins (Mm, Fs, and Vv) in host strains that support spacer acquisition from RNA compared to spacers acquired from DNA by an Mm RT-Cas1 mutant lacking the RT domain (Cas6-RTΔ-Cas1, denoted RTΔ-Cas1) and Cas1 proteins from *Thermus thermophilus* (Tt), *Staphylococcus epidermis* (Se), and *Streptococcus thermophilus* (St) Type III systems that lack an associated RT.^28,42,43^

The spacers acquired by these proteins have variable percentages (6-71%) of non-coded nucleotides at the spacer-repeat junctions (Figure 1B). However, the number of non-coded nucleotides at these junctions differed markedly for proteins that could or could not acquire spacers from RNA (Figure 1C). The majority of newly acquired spacers for the wild-type Mm, Fs, and Vv RT-Cas1 proteins, which acquire spacers from RNA, had two or more non-coded nucleotides at the spacer-repeat junctions, while deletion of the RT domain in Mm RTΔ-Cas1 sharply decreased the percentage of spacers that had ≥2 non-coded nucleotides, approaching the percentages for the Tt, Se, and St Cas1/Cas2 proteins, which only acquire spacers from DNA (Figure 1C). For WT Mm

RT-Cas1, the first two non-coded nucleotides at the spacer-repeat junctions were predominantly A residues (>50%) followed by G >T > C residues, while spacers acquired by Mm RTΔ-Cas1/Cas2 showed less bias for A residue and higher proportions of C and T residues at all positions (Figure 1D). The length distribution of *in vivo* spacers acquired by Mm WT RT-Cas1/Cas2 and RTΔ-Cas1/Cas2 were similar with peaks at 34-35 nt, but with spacers acquired by WT RT-Cas1/Cas2 requiring addition of 1 to 6 non-coded nucleotides to achieve a length distribution similar to that of spacers acquired from DNA by RTΔ-Cas1 (Figure 1E). These findings suggested that Mm RT-Cas1 has a terminal transferase activity that adds extra non-coded deoxynucleotides, preferentially dA residues, to the 3’ ends of RNA protospacers.

Prompted by these findings, we did terminal transferase assays with purified WT Mm RT-Cas1/Cas2 and 29-nt RNA or DNA oligonucleotide substrates (R29 and D29, respectively), in reaction medium containing 10 mM Mg^2+^ in the absence or presence of 1 mM Mn^2+^, a physiologically relevant divalent cation that modulates the activity of many polymerases.^44,45^ Consistent with *in vivo* findings, the assays showed that WT RT-Cas1/Cas2 has a terminal transferase activity that adds non-coded DNA tails to the 3’ end of RNA and DNA substrates with nucleotide preferences A >> G > C > T for the RNA substrate and A > C > G >T for the DNA substrate and Mn^2+^ strongly increasing both the activity and its preference for adding A residues to the 3’ end of the RNA substrate (Figure 1F). Additional terminal transferase assays with RNA and DNA oligonucleotide substrates having different sequences and 3’ nucleotides showed similar trends with somewhat less bias for A over G residues (Figure S1A). Notably, RT-Cas1/Cas2 terminal transferase had a propensity to stop or pause after addition of short (∼4 nt) dN tails, particularly evident for addition of dA tails to the RNA substrate (Figure 1F and Figure S1A), possibly a feature that keeps overall protospacer length in a range that can be accommodated by Cas1/Cas2. The greater preference of Mm RT-Cas1/Cas2 for adding A residues to the 3’ end of RNA than DNA substrate explains why spacers acquired by Mm WT RT-Cas1/Cas2 *in vivo* have a higher proportions of A residues at RNA proximal positions 1 and 2, but lower proportions of A residues at more distal positions 3 to 5, where nucleotide addition occurs to progressively longer stretches of DNA resulting from prior dNTP additions. Further analysis showed that spacers acquired *in vivo* by WT Mm RT-Cas1/Cas2 were uniformly distributed throughout host RNAs without enrichment for 3’ termini, as expected for spacer acquisition from fragmented RNAs (Figure S1B), and had significantly higher frequencies of non-coded AA, AG, and GG dinucleotides at non-coded nucleotide positions 1 and 2 than those acquired by Mm RTι1-Cas1/Cas (Figure S1C). Collectively, these findings suggested that addition of deoxynucleotides to the 3’ end of RNA fragment protospacers by RT-Cas1/Cas2 terminal transferase activity might play a role in RNA protospacer acquisition.

### Addition of deoxynucleotides to the 3’ end of RNA protospacers is required and sufficient for RNA spacer integration

To investigate if added 3’-dN tails are required for RNA spacer acquisition, we performed spacer acquisition assays with the WT Mm RT-Cas1/Cas2 in the presence or absence of added dNTPs. In an initial assay, we incubated WT Mm RT-Cas1/Cas2 with a double-stranded (ds) internally ^32^P-labeled CRISPR DNA and 29-nt ssDNA (D29) or ssRNA (R29) protospacers having the same nucleotide sequence in the presence or absence of each of the four dNTPs, dideoxy ATP (ddATP), a non-hydrolyzable dATP analog (dApCpp), or ATP. The reactions were done in the absence of Mn^2+^ to limit terminal transferase addition of non-coded nucleotides to the 3’ ends of the labeled CRISPR DNA, and the products were analyzed in a denaturing 6% polyacrylamide gel. Spacer ligation to the 5’ end of the first repeat (R1) on opposite strands was expected to occur via transesterification reactions that yield labeled top-strand ligation products corresponding to the cleaved leader (L, 40 nt) and the 29-nt protospacer (S0) linked to the 5’ end of R1 (S0+R1+S1, 77 nt) and labeled bottom-strand ligation products corresponding to L+R1+S0 (104 nt) and S1 (13 nt, not radiolabeled and run off the gel; schematic Figure 2A).

**Figure 2.**
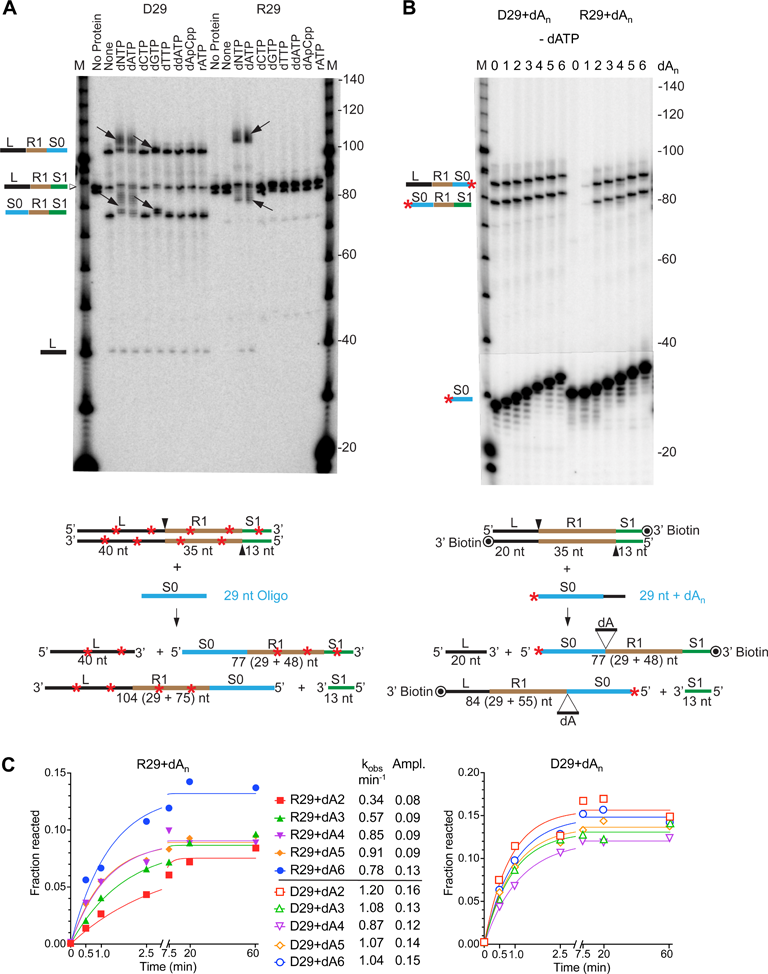
RNA spacer integration into a CRISPR array requires 3’ deoxynucleotides. (A) Spacer integration assays for DNA and RNA protospacers in the presence or absence of dNTPs. Reactions were done as described in Methods by incubating Mm WT RT-Cas1/Cas2 with 29-nt DNA or RNA protospacers having the same sequence (D29 and R29, respectively, denoted S0 in the schematic below) and an 88-bp internally ^32^P-labeled (red stars) CRISPR DNA (5’ L+R1+S1) in the presence of absence (None) of the indicated dNTPs for 1 h at 37°C. The reactions were done in the absence of Mn^2+^ to limit terminal transferase addition of dNTPs to the 3’ ends of the labeled CRISPR DNA. Products were analyzed in a denaturing 6% polyacrylamide gel against 5’-labeled 10-nt DNA ladder size markers run in parallels lanes (M). The arrows point to labeled top- and bottom-strand integration products whose lengths increased in the presence of dNTPs. The “No Protein” control lanes show the labeled CRISPR DNA incubated under identical conditions in the absence of RT-Cas1/Cas2 and dNTPs. ^32^P-labeled CRISPR DNA bands ran as closely spaced doublets due to slightly different electrophoretic mobilities of the two DNA strands. (B) Spacer integration assays done as in panel A by incubating Mm WT RT-Cas1/Cas2 with 5’-^32^P-labeled (red star) D29 and R29 oligonucleotides without or with different numbers of 3’-dA residues and an unlabeled 68-bp CRISPR DNA with a 3’-biotin blocking group. Lane M, 5’-labeled 10-nt DNA ladder size markers. (C) Time courses for integration of DNA and RNA protospacers with different length 3’-dA tails. Reactions were done with 5’-labeled oligonucleotides as in panel B for times up to 1 h. The plots show the fraction of labeled oligonucleotide inserted into the top and bottom strands of the CRISPR DNA as a function of time fit to a single exponential equation to obtain values for k_obs_ and amplitude. R^2^ values for curve fits are listed in the Supplemental data file. The schematics in the middle of the Figure show the substrates and products for the CRISPR integration reactions in panels A and B: incoming protospacer (S0) blue, CRISPR DNA leader (L) black, first repeat (R1) orange, and a 13-bp segment of the 5’-proximal spacer (S1, run off the gels) green. Red stars indicating internal ^32^P-labeling of CRISPR DNA in panel A and 5’-^32^P-labeled DNA or RNA protospacer in panel B.

The results showed that Mm RT-Cas1/Cas2 ligates DNA protospacers to the 5’ end of R1 on both strands in the presence or absence of added dNTPs, as expected, while efficient RNA protospacer integration occurred only in the presence of dNTPs or dATP, which is use most efficiently for adding dA tails in the absence of Mn^2+^ (Figure 2A; see also Figure 1F top). The DNA spacers integrated in the presence of dNTPs, dATP, or dGTP and the RNA spacers integrated in the presence of dNTPs or dATP were slightly longer than the initial protospacers as expected for terminal transferase addition of deoxynucleotide tails (Figure 2A, arrows in gel). The finding that dATP was by itself sufficient for RNA protospacer integration in the absence of other dNTPs indicates that RNA protospacer acquisition required only the addition of a 3’-dA tail to an RNA protospacer and did not require cDNA synthesis to generate a complementary cDNA strand (in agreement with a previous finding^12^).

To investigate how many non-coded 3’ deoxynucleotides are required for RNA spacer acquisition, we performed spacer acquisition assays with the same 29-nt DNA and RNA oligonucleotides without or with 1 to 6 dA residues added to their 3’ ends (Figure 2B). The oligonucleotides were 5’-^32^P-labeled (denoted by *) and used to assay spacer ligation into an unlabeled CRISPR DNA in the absence of added dNTPs or dATP. In this and subsequent experiments, the oligonucleotides comprising the CRISPR array were synthesized with a 3’-blocking group (biotin or dideoxy C (ddC)) to prevent terminal transferase addition of dNTPs to those ends.

The 5’-labeled ssDNA oligonucleotides with or without dA tails were efficiently ligated into the CRISPR DNA and produced the two expected ligation products (L+R1+S0, 84 nt and S0+R1+L, 77 nt), whose length increased progressively with the increasing length of the dA tail (Figure 2B, left side). By contrast, the 29-nt RNA oligonucleotide without a 3’-dA residue was not integrated into the CRISPR DNA and the RNA oligonucleotide with a single 3’-dA residue was integrated inefficiently, whereas RNA oligonucleotides with 2 or more dA residues at their 3’ ends were used more efficiently as protospacers (Figure 2B, right side).

Time courses comparing rates of integration (k_obs_) of the 5’-labeled 29-nt DNA or RNA protospacers with dA tails ranging in length from 2 to 6 nts showed relatively small differences for DNA protospacers but larger dependence on dA-tail length for RNA protospacers, with RNA protospacers having 4- to 6-nt dA tails integrating at rates close to those for DNA protospacers of the same length (Figure 2C and Figure S2A and B). Further experiments showed that adding dATP to integration reactions with 5’-labeled R29 and D29 oligonucleotides that had different length 3’-dA tails increased the length of the major integration products to a minimum of 33 nt (Figure S2C), matching the length distribution of spacers acquired by Mm RT-Cas1/Cas2 or RTΔ-Cas1/Cas2 *in vivo* (Figure 1E). Collectively, these findings show the Mm RT-Cas1/Cas2 terminal transferase activity adds 3’-dN tails to RNAs that match the characteristics of spacers acquired by RT-Cas1/Cas2 proteins *in vivo* and enable relatively efficient integration of RNA protospacers into a CRISPR array *in vitro* with no requirement for synthesis of a complementary cDNA.

### RT-Cas1/Cas2 synthesizes near full-length DNA copies of 50-nt RNA or DNA templates without an added primer in the presence or absence of Mn^2+^

To investigate how RT-Cas1/Cas2 synthesizes DNA copies of single-stranded RNAs or DNAs, we began by using 50-nt RNA or DNA oligonucleotide templates (R50CCC and D50CCC, respectively) of the same sequence that fortuitously included a 3’-proximal CCC sequence that turned out to be a preferred cDNA initiation site for Mm RT-Cas1/Cas2 (Figure 3). The RNA and DNA templates were tested without or with a 3’-ddC blocking group, which prevents both 3’-nucleotide addition by RT-Cas1 terminal transferase activity and “snap-back DNA synthesis”, a reaction in which the 3’ end of a DNA or RNA template folds back to prime DNA synthesis by base pairing to short complementary sequences upstream in the same the template.^46^ For initial experiments, DNA synthesis reactions were done by incubating the R50CCC and D50CCC templates with Mm RT-Cas1/Cas2 and ^32^P-dNTPs (a mixture of [α-^32^P]-dCTP + unlabeled dATP, dGTP and dTTP) in reaction medium containing 10 mM Mg^2+^ in the absence or presence of 1 mM Mn^2+^, and the products were analyzed in a denaturing 20% polyacrylamide gel.

**Figure 3.**
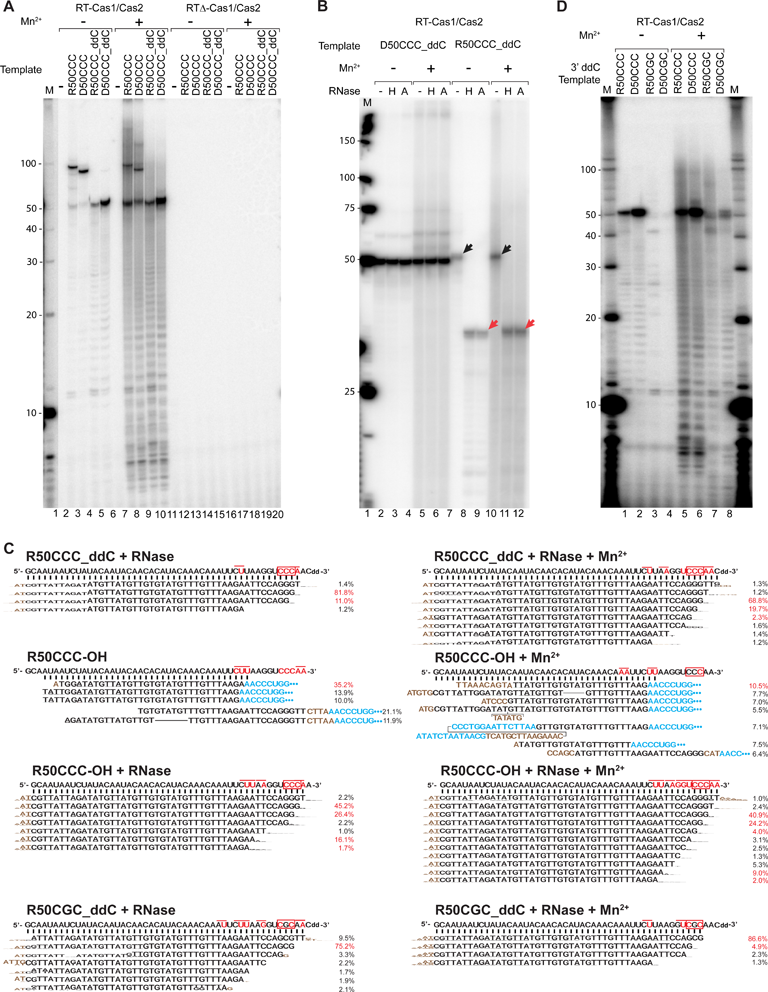
RT-Cas1/Cas2 synthesizes near full-length copies of RNA and DNA templates by initiating at 3’-proximal sites without an added primer. (A) Gel analysis of cDNAs synthesized by WT RT-Cas1/Cas2 (left) or RTΔ-Cas1/Cas2 (right) from 50-nt RNA or DNA oligonucleotide templates (R50CCC and D50CCC, respectively) without or with a 3’-ddC blocking group. DNA synthesis reactions were done in the presence of RNase-treated DNA products were analyzed in a denaturing 20% polyacrylamide gel, which was dried and scanned with a phosphorimager. Lane M, 5’-labeled 10-nt ssDNA ladder size markers run in a parallel lane. (B) Gel analysis showing that reverse transcription by RT-Cas1/Cas2 produces a stable RNA/cDNA duplex. DNA synthesis reactions were done as in panel A. After clean up with a Zymo Oligo Clean and Concentrator kit, the products were incubated for 20 min at 37°C in the presence or absence of RNase H or RNase A at a low salt concentration followed by Proteinase K digestion as described in Methods and analyzed by electrophoresis in a non-denaturing 15% polyacrylamide gel. Black arrows indicate the dsRNA/cDNA duplex, and red arrows indicate the single-stranded cDNA after RNase H or RNase A digestion of the RNA template strand. Lane M, 5’-labeled Low Molecular DNA Ladder size markers (New England Biolabs). (C) TGIRT-seq of cDNAs synthesized from R50CCC_ddC, R50CCC, and R50CGC_ddC (C47G) templates in reaction medium without or with added Mn^2+^. The RNA template sequence is shown above and sequences constituting >1% of the reads for cDNAs synthesized from 3’-blocked RNA templates or >5% of the reads for snap-back DNA synthesis products are shown below in sequence logo format with the height of the letter corresponding to the proportion of the nucleotide found at that position. Non-coded nucleotides are colored brown, and 3’-RNA template sequences used to prime snap-back cDNA synthesis are colored blue with blue dots indicating extended RNA template sequences that are not shown. Nucleotides in the RNA template sequence corresponding to first nucleotide of a cDNA product (N1) are highlighted in red, with a thin red line extending over positions N1 and N2 of the synthesized cDNA. The 3’-proximal CCC or mutant CGC sequence are in red boxes. (D) Gel analysis of DNA products synthesized by WT RT-Cas1/Cas2 from 3’-blocked 50-nt RNA or DNA templates with CCC or CGC at the prominent 3’-proximal CCC initiation site. RNase-treated DNA products were analyzed in a denaturing 20% polyacrylamide gel. Reverse transcription reactions for gel analysis were done as described in Methods by incubating WT RT-Cas1/Cas2 (left) or RTΔ-Cas1/Cas2 with or without (-) the indicated RNA or DNA template and ^32^P-labeled dNTPs ([α-^32^P]-dCTP (17 µM) plus 0.5 mM each of dATP, dGTP, and dTTP) in the presence or absence of 1 mM MnCl_2_ for 1 h at 25°C. TGIRT-seq libraries were prepared as described in Methods from parallel reverse transcription reactions using an equimolar mix of unlabeled dNTPs and sequenced on an Illumina NextSeq550 to obtain ∼1 million reads for each sample.

In the absence of a 3’-blocking, RT-Cas1/Cas2 gave two major products with both the RNA and DNA template, one of ∼100 nt, the size expected for a snap-back DNA synthesis, and the other of ∼50 nt, whose synthesis relative to the snap-back product was stimulated by Mn^2+^ and corresponds to a near full-length DNA copy of the RNA or DNA template beginning near its 3’ end (Figure 3A, left side). As expected, the ∼50-nt DNA product but not the snap-back DNA synthesis product was seen with the 3’-blocked R50CCC_dCC and D50CCC_dCC templates and neither product was seen with Mm RTΔ-Cas1/Cas2, which lacks RT and DNA polymerase activity (Figure 3A, right side). Analysis of the products on a non-denaturing polyacrylamide gel showed that incubation with RNase H or RNase A (lanes labeled H and A, respectively) had no effect on the electrophoretic mobility of the dsDNA resulting from copying of the DNA template, but increased the mobility of the product synthesized from the RNA template, indicating that it had been a stable RNA-cDNA heteroduplex (Figure 3B).

To identify putative cDNA initiation sites, we sequenced the cDNAs synthesized from the 50-nt RNA template with or without a 3’ blocking by using a Thermostable Group II Intron RT (TGIRT)-based DNA sequencing method (see Methods). The cDNA synthesis reactions were done with an equimolar mix of the 4 unlabeled dNTPs in the absence or presence of Mn^2+^. The sequences showed that >90% of the products synthesized from the 3’-blocked R50CCC_ddC template in the absence or presence of Mn^2+^ (left and right columns, respectively) were near full-length cDNAs that began at the CCC sequence near the 3’ end of the RNA (Figure 3C, row 1; cDNA initiation sites highlighted in red letters; 3’ CCC sequence highlighted in a red box). The remaining products began at 3’-proximal A or U residues, with the number of initiations sites increasing in the presence of Mn^2+^ (Figure 3C, top row). Most but not all of the cDNAs extended to 3’ end of the 50-nt RNA template (indicated by proportionately smaller letters in aligned sequences in WebLogo format) followed by non-coded TA residues, reflecting non-templated nucleotide addition by RT-Cas1/Cas1 upon reaching the 3’ end of the template, an activity found for group II intron RTs, as well as other RTs and DNA polymerases.^47,48^

To test if the lack of a 3’ OH in 3’-blocked RNA templates affects the use of cDNA initiation sites, we carried out DNA synthesis reactions with Mm RT-Cas1/Cas2 and the R50CCC template without or with a 3’-blocking group (R50CCC_OH and R50CCC-dCC, respectively) and sequenced the products without or with treatment with RNase A to degrade attached RNAs resulting from snap-back DNA synthesis (Figure 3C, rows 2 and 3, respectively). In the absence of RNase treatment, the major snap-back DNA synthesis initiation site identified as beginning with a sequence corresponding to the 3’ end of the RNA template used as a primer (blue) was a 3’-proximal UU complementary to 3’ AA of the RNA template (Figure 3C, row 2). Surprisingly, snap-back DNA synthesis initiation sites in the absence of Mn^2+^ also included the 3’-terminal AA of the RNA template, with cDNAs initiated at that site preceded by 4 or 5 non-coded nucleotides (brown), likely added by RT-Cas1 terminal transferase activity to give the 3’ end of the RNA template sufficient flexibility to snap-back and anneal to the complementary 3’ nucleotides (Figure 3C, row 2). Most of the snap-back products with attached RNA sequences terminated prior to reaching the 5’ end of the RNA template, possibly reflecting limited processivity of RT-Cas1/Cas2 (Figure 3C, row 2). Sequencing of the products from the same reaction after RNase treatment showed that most (∼70%) of the cDNAs began at the same 3’-proximal CCC sequence found for the 3’-blocked RNA template, with the remainder beginning at 3’-proximal A or U residues, including a substantial proportion beginning at the major UU snap-back DNA synthesis site upstream of the 3’ CCC (see above) and most extending to the 3’ end of the template followed by non-coded TA residues (Figure 3C, row 3). These findings indicated that aside from enabling snap-back DNA synthesis *in vitro*, a 3’ OH instead of a 3’-ddC blocking group did not appreciably affect Mm RT-Cas1/Cas2 cDNA initiation sites. The ability of RT-Cas1/Cas2 to initiate at 3’-proximal sites irrespective of the 3’ moiety was further supported by gel analysis of cDNAs synthesized from the same RNA template with other 3’-blocking groups or a 3’-phosphate (Figure S3A), a desirable characteristic enabling synthesis of near full-length cDNAs from RNA fragments generated by enzymes that leave different 3’ moieties.

Finally, to test the requirement for a dinucleotide CC sequence suggested to be a highly preferred initiation site for a Mn^2+^-dependent primase activity of Mm RT-Cas1/Cas2,^41^ we tested cDNA synthesis with ^32^P-labeled dNTPs from the 3’-blocked RNA and DNA oligonucleotide templates in which 3’-proximal CCC sequence was changed to CGC (R50CGC_ddC and D50CGC_ddC, respectively). Analysis of the products on a denaturing 20% polyacrylamide gel showed that this single nucleotide mutation strongly decreased production of near full-length cDNAs initiated at the trinucleotide site in the absence or presence of Mn^2+^ at the lower dCTP concentration used in this experiment for labeling with [α-^32^P]-dCTP (Figure 3D). However, sequencing of the products synthesized from the same RNA templates with an equimolar mix of unlabeled dNTPs showed that the CGC sequence remained a favored cDNA initiation site (80-90% of sequenced products) in the absence or presence of Mn^2+^, which was reported to be required for primase activity^41^ (Figure 3C). This dependence on dCTP concentration, confirmed separately by gel analysis (Figure S3B), suggested that the stability of base-pairing interactions rather than protein recognition of a specific sequence might be a significant factor in the selection of cDNA initiation sites by Mm RT-Cas1/Cas2. Collectively, these experiments showed that Mm RT-Cas1/Cas2 could initiate cDNA synthesis at different 3’-proximal sites without an added primer in the presence or absence of Mn^2+^, enabling the synthesis of near full-length DNA copies of these templates.

### RT-Cas1/Cas2 initiates cDNA synthesis at multiple 3’-proximal sites by protein-priming with different initiating dNTPs

The findings that Mm RT-Cas1 and a variety of other RTs could synthesize cDNAs that retained a 5’-triphosphate, a key indicator of *de novo* initiation, with a strong preference for initiation at CC sequences were based largely on experiments using short (7-20 nt) RNA substrates with limited sequence diversity.^41^ *De novo* initiation of cDNA synthesis at the CC of a 3’-tRNA-like structure was shown previously for a mitochondrial retroplasmid RT.^49^ However, the more diverse sequences of spacer acquired by Mm RT-Cas1/Cas2 *in vivo* led us to consider an alternate mechanism based on findings for the bacterial AbiK and Abi-P2 RTs. These group II intron-related RTs, which function in abortive phage infection, use non-templated protein-priming to synthesize long “random” sequence ssDNAs that contribute to altruistic cell death in response to phage infection.^18,20^ Protein priming resulting in covalent attachment of labeled nucleotides to an OH group of a tyrosine, threonine or serine is a well-characterized mechanism for the initiation of DNA synthesis by a number of viral and cellular RTs, but generally occurs at a fixed cDNA initiation site for those cased studied in detail.^20,50–52^

To investigate if protein-priming might be used for initiation of cDNA synthesis at the CCC initiation site, we incubated Mm RT-Cas1/Cas2 without or with the 3’-blocked R50CCC_ddC template and [α-^32^P]-dGTP in the presence or absence of Mn^2+^. After the reaction, the protein was analyzed by SDS-PAGE and autoradiography to detect covalently bound [α-^32^P]-dGTP as expected for protein-priming. As shown in Figure 4A, incubation of Mm RT-Cas1/Cas2 with [α-^32^P]-dGTP for 15 min in the presence but not absence of the RNA template resulted in strong Mn^2+^-dependent labeling of RT-Cas1 as well as more weakly labeled low molecular weight bands likely corresponding to ^32^P-dG oligomers (Figure 4A, lanes 1-4 and see below). Time courses showed that labeling of Mm RT-Cas1 by [α-^32^P]-dGTP increased progressively for times up to 60 min (Figure S4A). Adding an equimolar mix of unlabeled dNTPs as a chase after the initial 15-min labeling period with [α-^32^P]-dGTP resulted in the appearance of a higher molecular weight labeled DNA band that migrated above the major Coomassie-blue strain protein band, as well as increased intensity of the lower molecular weight bands (Figure 4A, lane 5). These additional bands were insensitive to digestion with RNase A, but sensitive to digestion with micrococcal nuclease (MNase), indicating that they were extended cDNA products (lanes 6, 7). Digestion with protease K shifted the major ^32^P-labeled Coomassie blue-strand protein band to a lower molecular weight and also resulted in disappearance of the labeled higher molecular weight MNase-sensitive band, suggesting that it was an extended cDNA associated with a fraction of the protein (Figure 4A, lane 8). The labeling of RT-Cas1 was not dependent upon the presence of Cas2, and incubating Cas2 by itself under the same conditions did not result in labeled protein (Figure 4A, lanes 9, 10). The same experiment with a 3’-blocked 50-nt DNA template of the same sequence gave similar results, but with somewhat lower protein labeling intensity compared to that with the 50-nt RNA template assayed in parallel (Figure 4A, lanes 11-21). These findings suggested that Mm RT-Cas/Cas2 could initiate DNA synthesis on RNA or DNA templates via Mn^2+^-stimulated protein priming.

**Figure 4.**
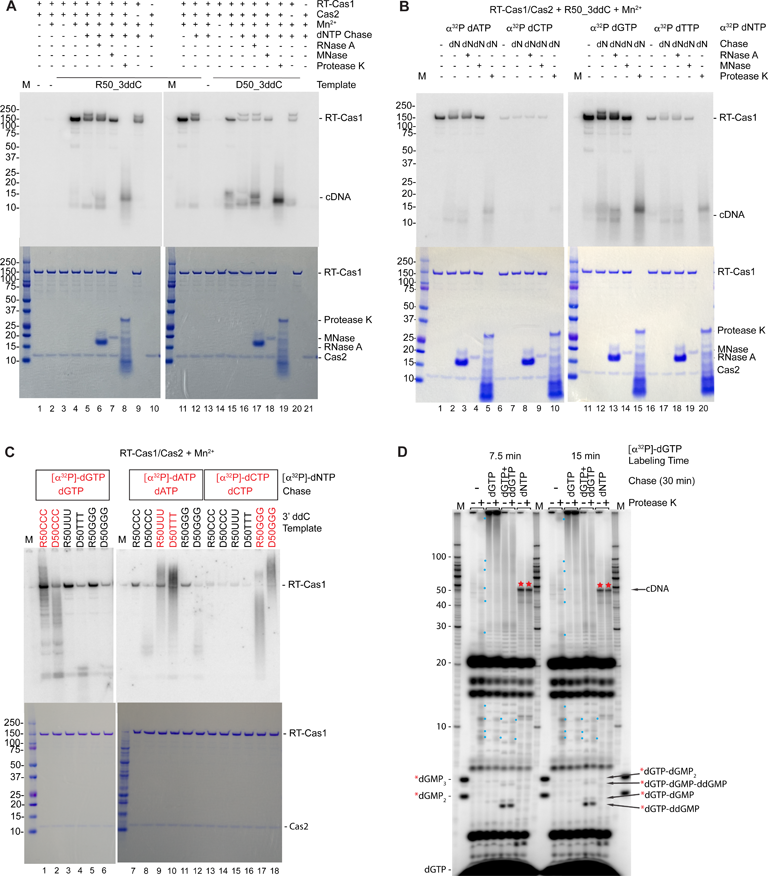
Mm RT-Cas1/Cas2 can initiate DNA synthesis on RNA and DNA templates by protein-priming. (A) Radiolabeling of RT-Cas1 by [α-^32^P]-dGTP in the presence or absence of RNA or DNA templates and other dNTPs. Reactions were done as described in Methods by incubating MM RT-Cas1/Cas2 (500 nM) or Cas2 (500 nM) with 5 µCi [α-^32^P]-dGTP (83 nM) in the presence or absence of 3’-blocked R50CCC and D50CCC templates and 1 mM MnCl_2_. After incubating for 15 min at 25°C, the reactions in the indicated gel lanes were chased with an equimolar mixture of unlabeled dNTPs (500 µM each) for 30 min at 25°C. Products of the reactions in the indicated gel lanes were incubated with RNase A, micrococcal nuclease (MNase) in the presence of Ca^2+^, or protease K for 15 min at 25°C. Two sets of samples, one for RNA (left) and the other for DNA (right) templates, were analyzed in parallel on SDS-polyacrylamide (4-12%) gradient gels (NuPAGE), which were stained with Coomassie blue (bottom panels), dried and scanned with a phosphorimager to detect labeled protein and DNA product bands (top panels). Lane M, Precision Plus (Bio-Rad) size markers. Bands corresponding to added proteins are labeled to right of the Coomassie blue-stained gels. (B) Radiolabeling of RT-Cas1 in the presence of 3’-blocked R50CCC_dCC template with different [α-^32^P]-dNTPs. Reactions were done in the presence of 1 mM Mn^2+^ and analyzed as in panel A. (C) Radiolabeling of RT-Cas1 with different [α-^32^P]-dNTPs in the presence of 3’-blocked RNA or DNA templates having different trinucleotides at the prominent 3’-proximal DNA synthesis initiation site (positions 46 to 48). Reactions were done in the presence of 1 mM Mn^2+^ with the indicated [α-^32^P]-dNTP and chased with higher concentrations of the same dNTP, and the products were analyzed as in panel A. (D) Pulse-chase experiment. RT-Cas1/Cas2 was labeled with [α-^32^P]-dGTP (83 nM) in the presence of R50CCC_ddC and 1 mM MnCl_2_ for 7.5 (left) or 15 min (right), followed by a 30 min chase with reaction medium without a dNTP (-) or with dGTP (500 µM), dGTP + ddGTP (500 and 1,000 µM, respectively), or an equimolar mix of dATP, dCTP, dGTP, and dTTP (500 µM each). Reactions were done in the presence of 1 mM Mn^2+^ and stopped with EDTA, incubated for 15 min without or with protease K (0.32 units), and analyzed in a denaturing 20% polyacrylamide gel, which was dried and scanned with a phosphorimager. Lanes M, 5’-labeled 10-nt ssDNA ladder size marker and a mixture 5’-labeled di- and tri-nucleotides (dGMP_2_ and dGMP_3_). The likely composition of the short oligonucleotide products indicated at the bottom the right of the gel was inferred from their expected differences in mobility relative to the 5’-labeled dGMP_2_ and dGMP_3_ markers. Small blue dots in the gel indicate additional ^32^P-labeled bands that appear after Proteinase K digestion putatively corresponding to peptides with covalently bound labeled DNA products. Red stars in the lanes indicate the labeled near-full length (∼50-nt) cDNA products that appeared after the chase with dNTPs.

To test if protein-priming by RT-Cas1/Cas2 was restricted to using dGTP as the initiating nucleotide, we carried out a similar experiment in which RT-Cas1/Cas2 was incubated with the R50CCC_ddC template in the presence each of the 4 possible ^32^P-dNTPs (Figure 4B). Analysis of the products on an SDS-polyacrylamide gel showed that RT-Cas1 was labeled by all 4 dNTPs with relative efficiency dGTP > dATP > dTTP > dCTP, and that in each case, chasing with an equimolar mix of unlabeled dNTPs (dN) resulted in higher or lower molecular weight bands corresponding to DNA products that were degraded by MNase but not RNase A (Figure 4B). These findings indicate that RT-Cas1/Cas2 could initiate DNA synthesis by protein priming with any dNTP, but with dGTP favored over dATP and purines favored over pyrimidines for initiation of DNA synthesis on the 3’-blocked R50CCC template.

To determine if a 3’-proximal CCC sequence is essential for protein priming, we compared protein labeling using 3’-blocked 50-nt RNA or DNA templates of otherwise identical sequence in which the 3’-proximal CCC was changed to UUU/TTT or GGG (Figure 4C). As in the previous experiment, RT-Cas1 was labeled by each of these dNTPs with efficiency dGTP > dATP > dCTP (Figure 4C). However, chasing with the same unlabeled dNTP resulted in a smear of dissociated ^32^P-labeled DNA products extending up the gel lanes for RNA or DNA templates that had a complementary 3’-trinucleotide sequence but not for those that had a non-complementary trinucleotide, reflecting synthesis of higher molecular products by reiterative copying of the trinucleotide in a sequence-dependent manner (Figure 4C and see below).

To confirm that protein-priming could give rise to free cDNA products, we incubated Mm RT-Cas1/Cas2 with [α-^32^P]-dGTP in reaction medium containing 10 mM Mg^2+^ plus 1 mM Mn^2+^ for 7.5 or 15 min and then chased the reactions with higher concentrations of unlabeled dGTP or an equimolar mix of all 4 dNTPs for 30 min. The products were then analyzed on a denaturing 20% polyacrylamide gel before or after digestions with protease K. Phosphorimager scans of the gels showed that [α-^32^P]-dGTP covalently bound to RT-Cas1 (protease K-sensitive label in well) was chased into larger dissociated DNA products, near full-length cDNAs (marked by red stars in gel lanes) by higher concentrations of all 4 dNTPs, and progressively longer dG oligomers extending up the gel lane by higher concentrations of dGTP in the absence but not presence of ddGTP (Figure 4D). The phosphorimager scan also showed a series of bands (marked by small blue dots) that appeared after protease K digestion in the lanes for the initial labeling periods, but were not visible in the dNTP-chase lanes (Figure 4D), suggesting that they corresponded to peptide fragments with covalently bound ^32^P-dGTP or short ^32^P-dG oligomers that were chased into larger products.

Surprisingly, the autoradiogram also showed a series of intensely ^32^P-labeled protease-insensitive lower molecular weight bands (up to ∼20 nt) that were not appreciably chased into larger products by higher concentrations of dGTP or dNTPs. Further experiments showed the intensely labeled bands were non-protein-associated ^32^P-dG oligomers as short as dinucleotides that accumulated over time in the absence but not appreciably in the presence of higher concentrations of dGTP or dNTPs at the beginning of the labeling period (Figure S4B). Instead, higher concentrations of dGTP at the beginning of the time course led to the synthesis of longer dG oligomers by reiterative copying of the CCC sequence, while higher concentrations of dNTPs at the beginning of the labeling period led to the synthesis of near full-length cDNAs of the RNA template (Figure S4B). These findings suggest that short DNA oligomers synthesized *de novo* or by rapid release after protein priming at early time points were used to prime synthesis of longer DNA products.

Collectively, our findings indicated that Mm RT-Cas1 could use protein-priming to initiate cDNA synthesis at 3’-proximal sites with different trinucleotide sequences, either reiteratively copying those sequences to generate DNA oligomers in the presence of a single complementary dNTP or synthesizing near full-length cDNAs initiated at the trinucleotide in the presence of all 4 dNTPs. Protein priming enabled initiation of cDNA synthesis with different efficiencies by any dNTP, including with dGTP at a 3’-proximal CCC containing a CC dinucleotide that is a highly preferred site for *de novo* initiation.^41^ These experiments also showed that Mm RT-Cas1/Cas2 synthesizes non-protein-associated DNA oligomers, either *de novo* or by rapid release after protein priming, with short DNA oligomers synthesized at early time points potentially used as primers for synthesis of longer cDNA products. Although our results do not preclude use of a primase mechanism for initiation of cDNA synthesis at some sites, time courses comparing rates of cDNA synthesis from the 3’-blocked-R50CCC template with 500 μM dNTPs in the absence or presence of 20 μM dG_2_ primer showed a lag for initiation of DNA synthesis in the absence of the dG_2_ primer, most likely reflecting the time needed to synthesize a dG oligomer primer before beginning processive cDNA synthesis (Figure S4C).

### Mm RT-Cas1/Cas2 initiates cDNA synthesis at complementary 3’-proximal sites by using 2-nt DNA oligonucleotide primers

To test systematically if Mm RT-Cas1/Cas2 could use short DNA oligomer primers to initiate DNA synthesis at complementary sites in an RNA template, we incubated Mm RT-Cas1/Cas2 with 3’-blocked RNA templates R50_ddC templates with each of 4 different 3’ proximal trinucleotide sequences and 5’-labeled dA_2_, dC_2_, dG_2_, or dT_2_ dinucleotide primers. The reactions were done in the presence of high concentrations of the unlabeled dNTP matching that in dinucleotide *(e.g.,* dATP for the dA_2_ primer) or an equimolar mixture of all 4 unlabeled dNTPs in the absence of Mn^2+^ to limit terminal transferase addition to the dN_2_ primer (visible as short DNA ladders in some gel lanes; Figure 5A). Analysis of the products in a denaturing 20% polyacrylamide gel showed that when incubated with high concentrations of the same unlabeled dNTP, RT-Cas1/Cas2 reiteratively copied the complementary trinucleotide in each template, generating a ladder of DNA homopolymers extending up the gel lane for those RNA templates that contained the complementary trinucleotide but not for the other templates (Figure 5A). When the reactions were done with an equimolar mixture of all 4 unlabeled dNTPs, RT-Cas1/Cas2 switched from reiterative copying of the complementary 3’ trinucleotide to synthesis of prominent longer cDNAs extending up to ∼50-nt cDNAs only for those templates that contained the complementary 3’ trinucleotide (including both R50AAA and R50GGG for the dT_2_ primer; Figure 5A). Sequencing of the cDNAs synthesized after the dNTP chase confirmed that the major initiation sites for each dinucleotide were a complementary di- or trinucleotide toward the 3’ end of the RNA template, entirely for the dA_2_, dC_2_, and dT_2_, but with the dG_2_ primer also showing small proportions of initiations at U and non-complementary A residues, the latter possibly reflecting a non-Watson-Crick guanosine-adenosine pairing (Figure 5A, right).^53^ The ability to use very short DNA primers to initiate cDNA synthesis at complementary sites is enabled by the strong high strand-annealing activity of group II intron-like RTs.^22^

**Figure 5.**
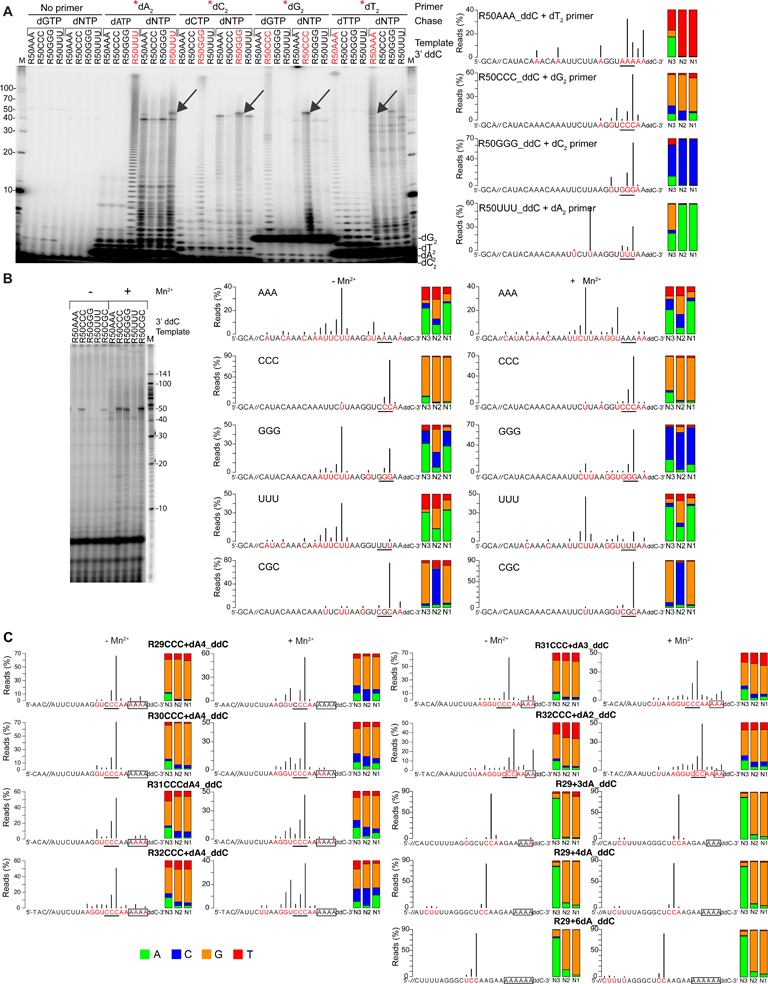
Mm RT-Cas1/Cas2 cDNA start sites on different RNA templates in the presence or absence of exogenous short DNA oligonucleotide primers. (A) Analysis of cDNAs synthesized by RT-Cas1/Cas1 using exogeneous dinucleotide primers. Left, reverse transcription reactions were done by incubating Mm WT RT-Cas1/Cas2 (500 nM) with 250 nM 3’-blocked R50NNN templates with AAA, CCC, GGG, UUU or CGC at nucleotide positions 46-48 and 20 µM 5’-^32^P-labeled (*) dinucleotide primers dA_2_, dC_2_, dG_2_, dT_2_ or no primer in the absence of Mn^2+^ to minimize terminal transferase addition of dNTPs to the 3’ end of the labeled primers. After incubating for 30 min at 25°C, the reactions were chased for 30 min with 500 µM of the dNTP matching the dinucleotide primer or an equimolar mix of all 4 dNTPs (500 µM each). 5’-labeled cDNAs were analyzed in a denaturing 20% polyacrylamide gel against a 5’-labeled 10-nt ssDNA ladder (lanes M). Right, TGIRT-seq analysis of cDNAs from parallel reverse transcription reactions for the same templates with unlabeled dinucleotides and an equimolar mix of all 4 dNTPs (500 µM each). Reactions were incubated for 1 h at 25°C, and cDNAs were analyzed by TGIRT-seq, as described in Methods. Nucleotides in the RNA template that correspond to the first nucleotide of a cDNA product are highlighted in red with black vertical lines indicating the percentage of reads beginning at that nucleotide. The 3’-proximal trinucleotide sequence in each template is underlined. The stacked bar graphs at the right show the proportions of different nucleotides at the N1 to N3 positions of the cDNA color coded as shown at bottom left of panel C. (B) Analysis of cDNAs synthesized from the same templates as in panel A in the absence of an added primer. Reverse transcription reactions were done by incubating Mm RT-Cas1/Cas1 and RNA templates at the same concentrations as in panel A in the absence or presence of 1 mM Mn^2+^ for 1 h at 25°C. Left, reactions for gel analysis were done with [α-^32^P]-dCTP (83 nM) plus an equimolar mix of unlabeled dATP, dCTP, dGTP, and dTTP (500 nM each) and analyzed in a denaturing 20% polyacrylamide gel. Right, TGIRT-seq of cDNAs synthesized in parallel reactions with unlabeled dNTPs. The datasets for the R50CCC_ddC and R50CGC_ddC templates are the same as those in Figure 3C. (C) TGIRT-seq of cDNAs synthesized from RNA templates with different length RNA and dA-tail segments. The RNA segments were shorter versions of the R50CCC template used in Figure 3 (R29-32CCC) or the R29 substrate used for terminal transferase assays in Figure 2 with a 3’-ddC blocking group at the end of the DNA segment. Reverse transcription reactions were done and analyzed by TGIRT-seq as in panel (A). Gel analysis of ^32^P-labeled cDNAs synthesized in parallel reactions with [α-^32^P]-dCTP are shown in Figure S5.

### Mm RT-Cas1/Cas2 initiation sites are dictated in part by the stability of base-pairing interactions over several nucleotides

To investigate the propensity of RT-Cas1/Cas2 to initiate DNA synthesis at different sites in the absence of an added primer, we incubated RT-Cas1/Cas2 with 3’-blocked R50 templates with different 3’ trinucleotide sequences and [α-^32^P]-dCTP plus 500 μM unlabeled dATP, dCTP, dGTP, and dTTP in the presence or absence of Mn^2+^ (Figure 5B). Analysis of the products in a denaturing 20% polyacrylamide gel showed a prominent cDNA band of ∼50 nt cDNA for the 3’-blocked R50CCC, R50GGG, and mutant R50CGC templates, whereas the most prominent bands for the R50UUU, R50AAA, and R50GGG templates were a series of cDNAs ranging in size from 30 to 40 nt (Figure 5B).

To identify putative cDNA initiation, we sequenced the cDNAs synthesized from the different RNA templates in the presence or absence of Mn^2+^ (Figure 5B). The sequencing showed that most (75-92%) of the cDNAs synthesized from the R50CCC_ddC and R50CGC_ddC templates in the presence or absence of Mn^2+^ began at the CCC trinucleotide site, with <15% beginning opposite A or U residues (Figure 5B, rows 2 and 5, data taken from Figure 3C). The first nucleotide of most of the cDNAs initiated on the CCC and CGC templates corresponded to a G residue, while the second nucleotide corresponded to a G residue for the CCC template and a C residue for the CGC template, as expected for *de novo* initiation^41^ or protein priming (Figure 4) at the 3’-proximal C residues of the trinucleotide in both templates (Figure 5B). None of the cDNAs synthesized from the CGC template began at the middle G residues, reflecting that dCTP is used less efficiently for *de novo* or protein-primed initiation of cDNA synthesis than dGTP (Figure 5B).

Switching the 3’-proximal CCC to AAA or UUU, resulted in cDNAs beginning at a nearby upstream AAUUCUU sequence, including but not limited to its 3’ UU sequence used for initiation of snapback DNA synthesis with an RNA template having a 3’ OH (Figure 3B) and surprisingly few initiations within the 3’ AAA or UUU trinucleotide sequences (Figure 5B, rows 1 and 4). Initiation at these upstream sites agrees with the shorter sizes of ^32^P-labeled cDNAs synthesized from these templates in denaturing polyacrylamide gels (Figure 5B and Figure S3B). The favored nucleotide for initiating cDNA synthesis on these templates was dATP (59-70%) followed by dGTP, dTTP, and dCTP (Figure 5B).

Finally, switching the 3’-proximal trinucleotide to GGG also resulted in increased use of initiation sites within the nearby upstream AAUUCUU sequences but with a surprisingly high proportion of cDNAs initiated at the 3’-proximal GGG sequence in the absence of Mn^2+^ and with the 3’-proximal GGG becoming the predominant initiation site in the presence Mn^2+^ (Figure 5C, row 3). In this case, the major, almost exclusively used initiating nucleotide in the presence of Mn^2+^ was dCTP, which can form a stable base pair with G but is the least favored dNTP for protein priming and likely *de novo* initiation by RT-Cas1/Cas2 (Figure 5B). Collectively, these findings indicate that in addition to proximity to 3’ end of the RNA, the stability of base-pairing interactions over several nucleotides can override specific sequences for the choice of cDNA initiation sites on RNA templates, as expected for initiation of cDNA synthesis by annealing of short DNA oligomer primers.

### Mm RT-Cas1/Cas2 initiates at 3’-proximal sites in the RNA segment of 3’-dA tailed RNAs

To investigate how cDNA synthesis is initiated on RNA protospacers with 3’-DNA tails, we tested the effect of varying the lengths of the RNA and 3’-DNA-tail length segments on two different sets of 3’-blocked RNA protospacers, one set (R29-32CCC+dAn) corresponding to the 3’ end of the R50CCC template used above to identify cDNA initiation sites (Figures 3 and 5), and the other (R29+dAn) corresponding to the R29 RNA used to analyze RNA protospacer integration with or without a 3’-dA tail (Figure 2). For both templates, we found that preferred initiation sites in the presence or absence of Mn^2+^ were located toward the 3’ end of the RNA segment, with few initiations occurring within the 3’-dA tail (Figure 5C; gels shown in Figure S5A and S5B). Sequencing showed that the major initiation sites for the R29-32CCC+dAn templates were clustered at or near the CCC trinucleotide, the favored initiation site for the R50CCC template, but with higher proportions outside the CCC sequence, while the major initiation sites for the R29+dAn templates were at a 3’-proximal CC dinucleotide sequence with minor initiation sites elsewhere in the RNA segment (Figure 5C). For both sets of templates, few if any cDNA starts sites were in the 3’-dA tail (Figure 5C). The initiation of cDNA synthesis sites at 3’-proximal sites in the RNA segment upstream of the 3’-dA tail yields RNA/DNA duplexes with a single-stranded 3’-DNA overhang, favored for integration dsDNA protospacers into CRISPR arrays by Cas1/Cas2.^54^

### Kinetic analysis of RNA and DNA protospacer integration

Finally, to investigate the mechanism by which dN-tailed RNA protospacers are integrated into CRISPR arrays, we carried out kinetic assays of spacer integration. For these assays, we used a CRISPR hairpin DNA substrate in which the top and bottom strands were connected by a 5-nt linker, making it possible to identify products resulting from coupled cleavage-ligation of 5’-labeled protospacers into either or both strands by the length of labeled DNA fragments (Figure 6A). The reactions were done with R30CCC or R29 oligonucleotide protospacers with different length 3’-dA overhangs on the top and bottom strands in the absence of Mn^2+^ or dNTPs (Figure 6B-D). In each case, we compared the rates of integration by WT RT-Cas1/Cas2 or RTΔ-Cas1/Cas2 in parallel assays with 4 different 5’-labeled (*) protospacers: a 3’-dA-tailed *RNA (*RNA-dA), a single-strand *DNA, an *RNA-dA/DNA duplex, and an RNA-dA/*DNA duplex, with the RNA-dA and DNA strands having complementary nucleotide sequences except for the dA-tails (plots shown in Figure 6B-D; gels shown in the Supplementary data file). The integration of a stable *RNA-dA/cDNA duplex in a similar spacer integration assay was confirmed by gel electrophoresis of labeled integration products before and after RNase H digestion (Figure S6A).

**Figure 6.**
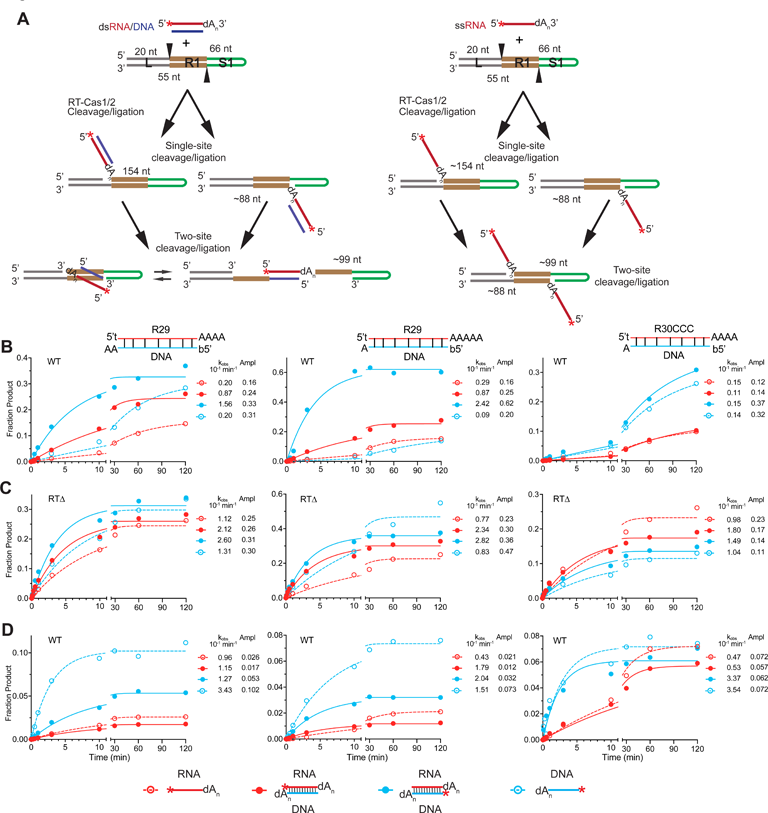
Time courses comparing rates of integration of single-stranded and duplex RNA and DNA protospacers into a CRISPR array. (A) Schematic of spacer integration assays. The assays used a CRISPR hairpin DNA substrate. consisting of a 5’ leader (L, gray), first repeat (R1, orange), and a hairpin corresponding to a segment of the first spacer (S1), green. Spacer integration assays were done by incubating 5’-labeled (red star) 3’-dN-tailed RNAs (RNA-dN; red) with or without a complementary cDNA strand (blue) that leaves different length 3’-dA overhangs on both strands. Products where the unlabeled strand is inserted into the CRISPR DNA are not shown. For the duplex integration pathway only one possible orientation of the two-site cleavage product is shown with or without base-pairing of the repeat. (B) Time courses for spacer integration by WT RT-Cas1/Cas2. Reactions were done as described in Methods by incubating WT RT-Cas1/Cas2 (500 nM) and CRISPR hairpin DNA (100 nM) with a 5’-^32^P-labeled protospacer (5 nM) for times up to 2 h. The schematics above the plots show the configurations for different spacers that were tested for R29-R30CCC-based protospacers. For each configuration, assays were done with 4 different protospacer substrates: a single-stranded 5’ labeled RNA-dN (*RNA-dA), an *RNA-dA/DNA duplex, an RNA/*DNA duplex, and a ssDNA, with t or b indicating the 5’-labeled strand. The plots show production of the labeled 99-nt band resulting from cleavage-ligation reactions at the 5’ end of R1 on both strands. The data were fit to a single exponential equation for calculation of the rate (k_obs_) and amplitude (Ampl). Red, 5’ labeled RNA-dA; blue, 5’ labeled DNA; open symbols, single-stranded RNA or DNA; closed symbols; double-stranded protospacers. (C) Time courses for spacer integration by RT1′-Cas1/Cas2 done as in panel (B). (D) Time courses for spacer integration by WT RT-Cas1/Cas2 done panel B with higher (250 nM) protospacer concentration. Products of the reactions were analyzed in a denaturing 8% polyacrylamide gel, which was dried and scanned with a phosphorimager. Bands were quantitated with ImageQuant software. Phosphorimager scans of the gels are shown in the Supplementary data file. For the reactions shown in Figure 6B, plots showing production of all labeled single-site and two-site cleavage-ligation products in the same graph are shown in Figure S6A-C. Protospacers having the same sequence and overhang configuration with a DNA-dA top strand instead of an RNA-dAf top strand are shown in Figure S7. R^2^ values for the plots are listed in the Supplemental data file.

At an initial protospacer concentration tested (5 nM), the rates and amplitudes for production of the 5’-labeled 99-nt cleavage-ligation product expected for spacer integration in both strands was generally higher for the *RNA-dA/DNA or RNA-dA/*DNA duplexes (closed red and blue circles) than for their *RNA or *ssDNA counterparts (open red and blue circles), as expected for integration of duplexes enabling more rapid sequential ligation into both the top- and bottom- strands (Figure 6B; time courses for all labeled cleavage products shown in Figure S6B-D). For WT RT-Cas1/Cas2, the rates and amplitudes for integration of the *RNA-dA or *RNA-dA/DNA duplex (red open and closed circles, respectively) were generally lower than those for their *ssDNA or RNA-dA/*DNA counterparts (blue open and closed circles, respectively), indicating more efficient integration of the labeled DNA strand, either as a *ssDNA or RNA-dA/*DNA duplex (Figure 6B).

When the same reactions were done with RTι1-Cas1/Cas2, however, the rates of integration increased for all substrates tested, but with increases in rates and amplitudes larger for the *RNA- dA and *RNA-dA/DNA duplexes (red open and closed circles, respectively) than for their *DNA counterparts (blue open and closed circles, respectively), enabling more similar and in one case higher rates and amplitudes for the *RNA-dA and RNA-dA/*DNA protospacers than for their similarly configured *DNA counterparts (Figure 6C). These findings may reflect that the RT domain impedes RNA spacer integration when not directly coupled to cDNA synthesis and suggest there may be relatively little inherent difference in the efficiency of integration of dN-tailed RNA and DNA protospacers.

Notably, the preference for integration of duplex over RNA-dA or ssDNA protospacers was concentration dependent. In reactions done at higher protospacer concentrations (250 nM), both the *ssRNA-dA and *ssDNA protospacers (red and blue open circles, respectively) had higher rates and amplitudes than did their labeled duplex counterparts (red and blue closed circles, respectively), with the preference for ssDNA protospacers being particularly striking for the two R29 protospacers (Figure 6D). These findings suggest that integration of single-strand dN-tailed RNA or ssDNA protospacers might be favored *in vivo* for abundant pathogen RNAs and ssDNA fragments generated by RecBCD or other phage defense nucleases.^55–57^

To directly compare the relative efficiencies of RNA/DNA vs. DNA/DNA protospacers of the same sequence and overhang configuration, we did assays in parallel with those above for protospacers in which the *RNA-dA strand was replaced with a *DNA-dA strand of the same sequence with a complementary DNA strand for duplex protospacers (Figure S7). The integration time courses at 5 nM protospacer concentration generally paralleled those for the RNA-dA protospacers, but with somewhat higher rates and amplitudes for both WT RT-Cas1/Cas2 (0.8- to 1.9-fold for rates, except for the D30CCC-dA/DNA protospacers which were 10- to 18-fold higher, and ≤2 fold for amplitudes) and RTΔ-Cas1/Cas2 (0.7- to 2.9-fold for rates and ≤1.6 fold for amplitudes; Figure S7A and B, compare with Figure 6B and C). In addition to the difference in behavior of the D29-based and D30CCC protospacers, the parallels included the top-strand-labeled DNA-dA protospacers having relatively lower amplitudes than the bottom-strand labeled protospacers for integration by WT RT-Cas1/Cas2 and more similar or one case higher amplitudes for integration by RTΔ-Cas1/Cas2 (Figure S7A and B, compare with Figure 6B and C; time courses for all labeled cleavage products for WT RT-Cas1/Cas2 shown in Figure S7C). The latter findings indicates that differences in protospacer sequences likely contributed to the differences in the relative efficiencies of RNA-dA and DNA protospacers in Figure 6.

## DISCUSSION

Here, we elucidated mechanisms and pathways used by Mm RT-Cas1/Cas2 for cDNA synthesis and site-specific integration of RNA-derived protospacers into CRISPR arrays (Figure 7). All pathways begin with RT-Cas1/Cas2 using its terminal transferase activity to add short DNA tails to the 3’ ends of RNA fragment protospacers generated by RNases *in vivo*. In one set of pathways (Figure 7, left side), RT-Cas1/Cas1 synthesizes near full-length cDNAs of the 3’-dN-tailed RNAs by initiating cDNA synthesis at 3’-proximal sites *de novo*, by protein priming, or by annealing short exogenic or synthesized DNA oligomer primers. The resulting 3’-dN-tailed RNA/DNA duplexes with deoxynucleotides at the 3’ ends of both strands are then ligated into the CRISPR array by a mechanism analogous to that used by conventional Cas1/Cas2 proteins to integrate dsDNA protospacers intro CRIPSR arrays. In a second set of pathways favored a higher protospacer concentrations (Figure 7A, right), 3’-dN-tailed RNAs are integrated directly into the CRISPR array prior to cDNA synthesis. Conversion of the integrated 3’-dN-tailed RNA into a fully integrated dsDNA spacer could then occur by cDNA synthesis after integration or possibly without cDNA synthesis by linked integration/disintegration reactions at CRISPR insertion sites on opposite strands.^58^ Analogous pathways could also be used for integration ssDNA protospacers generated by phage defense nucleases. In all pathways, the resulting integrated protospacer has single-strand gaps and is held together by Cas1/Cas2 until it is dissociated by transcription-coupled DNA repair that fills in single-stranded gaps and seals unconnected DNA segments to fully integrate the newly acquired spacer into the CRISPR array.^39^ Our analysis of these mechanisms explains known features of RNA-spacer acquisition by Mn RT-Cas1/Cas2 *in vivo* and revealed two novel biochemical activities with potential biotechnological applications: the ability of an RT to use multiple mechanisms to synthesize near full-length cDNAs of diverse RNA templates without an added primer or fixed sequence requirements, and the ability of a DNA integrase to site-specifically integrate RNAs into a DNA genome by adding deoxynucleotides at a crucial location.

**Figure 7.**
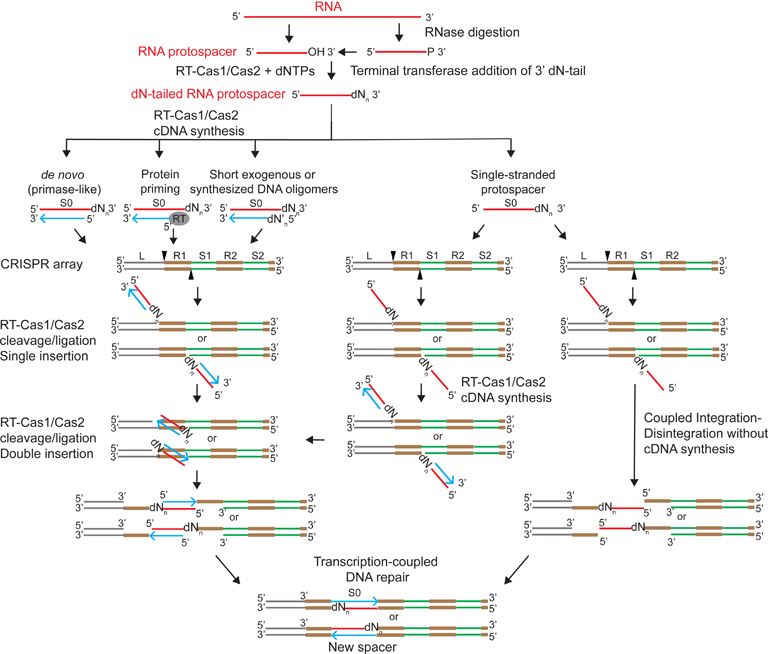
Pathways and mechanism used for RNA protospacer acquisition by Mm RT-Cas1/Cas2. In all pathways a host cellular or pathogen RNA is digested into fragments by RNases that leave a 3’ OH or a 3’ phosphate that can be removed by cellular enzymes that leave a 3’ OH, enabling addition of short DNA tails (dN_n_) by RT-Cas1/Cas2 terminal transferase activity. On the left are pathways in which RT-Cas1/Cas2 uses different mechanism (*de novo* initiation, protein priming, or exogenous or synthesized short DNA oligomer primers) to synthesize cDNAs that remain annealed to the RNA-dN template in an RNA-dN/cDNA duplex. The duplex protospacers are then integrated into the CRISPR array by RT-Cas1/Cas2 via a mechanism analogous to that used by conventional Cas1/Cas2 proteins to integrate duplex DNA protospacers. Only one orientation of the fully integrated spacer is shown in the Figure. On the right are alternative pathways in which 3’-dN-tailed RNA protospacers are directly integrated into the CRISPR array prior to cDNA synthesis. The integrated RNA-dN protospacers could then be used as a template for cDNA synthesis by RT-Cas1/Cas2, resulting in an RNA-dN/cDNA duplex that becomes fully integrated into the CRISPR array by a mechanism analogous to the second step for DNA duplex protospacers (horizontal arrow pointing left). Alternatively, a single-stranded RNA-dN protospacer integrated at one site could in principle be integrated into both sites by coupled integration/disintegration reactions in the absence of cDNA synthesis.^56^ In all pathways, protospacer integration leaves DNA segments with single-strand gaps that are held together by the bound RT-Cas1/Cas2 protein until it is displaced by transcription-coupled DNA repair, enabling complete integration of the spacers without introducing deleterious double-stranded breaks.^56^

The requirement for addition of short 3’-dN tails to enable direct integration of RNA protospacers by RT-Cas1/Cas2 proteins was suggested by the finding that spacers acquired by RT-Cas1/Cas2 proteins *in vivo* have larger numbers of non-coded nucleotides at spacer-repeat junctions than do those acquired by Mm RT-ΔCas1/Cas2 or Cas1/Cas2 proteins from CRISPR systems that acquire spacers only from DNA (Figure 1B and C). Biochemical assays confirmed this requirement by showing that the non-coded nucleotides at spacer-repeat junctions of spacers acquired by Mm RT-Cas1/Cas2 *in vivo* matched the nucleotide preferences of Mm RT-Cas1/Cas2 terminal transferase *in vitro* (Figure 1D) and that Mm RT-Cas1/Cas2 could directly integrate single-stranded 3’-dN-tailed RNA protospacers into CRISPR arrays both in coupled terminal transferase/spacer-ligation reactions in the presence of a single nucleotide (dATP) and in uncoupled spacer integration reactions using synthetic RNA oligonucleotides with different length 3’-dA tails (Figure 2). The latter assays showed that addition of a single 3’-dA residue enabled detectable RNA protospacer integration and addition of as few as 2 to 6 3’-dA residues enabled increased rates of RNA protospacer integration to levels similar to those of ssDNA protospacers with the same nucleotide sequence (Figure 2). RNA protospacers with a 3’ OH needed for terminal transferase addition of DNA tails could be generated by a cellular RNase, such as RNase III, which cleaves double-strand regions of structured RNAs to generate RNA fragments with a 3’ OH;^59^ by phosphatase removal of 3’ phosphates left by other RNases; or by an inherent RT-Cas1/Cas1 endonuclease activity that cleaves upstream of a 3’-blocked RNA, an activity reported for telomerase RT.^60^

As desired for an enzyme whose biological function is to acquire RNA-derived spacers that defend against diverse pathogens, we found that Mm RT-Cas1/Cas2 could use multiple mechanisms to initiate cDNA synthesis on RNA protospacers at different 3’-proximal sites with no fixed sequence requirements. In addition to recently reported *de novo* initiation with a strong preference for initiating at CC sequences,^41^ we found that Mm RT-Cas1/Cas2 can use protein-priming to initiate at multiple 3’-proximal sites with any dNTP, greatly expanding the number of suitable cDNA initiation sites (Figures 4 and 5). We also found that Mm RT-Cas1/Cas2, like other group II intron-related RTs,^22^ has a strong strand-annealing activity that enables it initiate site-specific cDNA synthesis by using primers as short as dinucleotides (Figure 5A). Short DNA oligomer primers could be generated *in vivo* by a variety of cellular enzymes. Particularly cogent, however, are phage defense enzymes, such as RecBCD that has both DNA exonuclease and DNA unwinding activities, enabling it to generate short (3 to 6 nt) ssDNAs that could be used to prime DNA synthesis on phage DNA protospacers and possibly suitably sized degradation intermediates that could be directly integrated into a CRISPR array by RT-Cas1/Cas2 proteins with or without a complementary DNA strand (Figure 6).^55–57,61,62^

Our findings also provide evidence that Mm RT-Cas1/Cas2 can use *de novo* initiation or protein priming to synthesize short DNA oligomer primers by reiteratively copying short repeated sequences and use these short oligomers to initiate DNA synthesis at complementary sequences on RNA or DNA templates (Figures 4 and 5). Supporting this mechanism, we found that the stability of base-pairing interactions over several nucleotides plays a substantial role in the selection of cDNA initiation sites (Figure 5). Most compelling was the finding that both CCC and GGG at the same 3’-proximal location in otherwise identical RNA templates were both preferred initiation sites for cDNA synthesis (Figure 5B), even though dCTP is the least favored dNTP for protein priming (Figure 4B) and likely also for *de novo* initiation. Initiation at G residues with dCTP at the 3’ proximal GGG sequence trumped initiation at sites with C, U, or A residues that were otherwise favored for cDNA initiation in otherwise identical templates that lacked a 3’ GGG (Figure 5B). Although our results do not exclude a primase-like mechanism, time courses for initiation of cDNA synthesis at a 3’-proximal CCC sequence with 500 μM dNTPs showed a pronounce lag for initiation of cDNA synthesis compared to a parallel assay in the presence of 20 μM dinucleotide dG2 primer, most likely reflecting the time needed for RT-Cas1/Cas2 to synthesize a short DNA primer (Figure S5C).

Unlike the AbiK and Abi-P1 RTs, which synthesize random sequence DNA oligomers in the absence of an RNA or DNA template,^20^ the synthesis of short DNA oligomers by Mm RT-Cas1/Cas2 required the presence of an RNA template (Figure 3D, 4D, S3B). This difference may reflect that the AbiK and Abi-P1 RTs have structural differences that prevent binding of a template but leave the RT active site accessible to dNTPs,^20^ while group II intron RT apoenzymes have a tendency to fold into an inactive structure in which the RT active site is blocked until activated by template binding.^63^ If so, the synthesis of short DNA oligomer primers by Mm RT-Cas1/Cas2 could occur by a *de novo* or protein-primed, non-templated mechanism analogous to that used by the Abi RTs.^20^ If not, the synthesis of short DNA oligomer primers by Mm RT-Cas1/Cas2 could involve transient association with an RNA template until the primer has sufficient length and base-pairing stability to initiate processive DNA synthesis. The finding that group II intron, LINE-1, telomerase, and retroviral RTs are capable Mn^2+^-dependent *de novo* initiation of cDNA synthesis,^41^ raises the possibility that *de novo* synthesis of short DNA oligomer primers may be an inherent activity of many if not all RTs.

The ability of RT-Cas1/Cas2 to integrate RNA-dN/cDNA duplexes with overhanging single-stranded DNA tails into a CRISPR array is consistent with and in retrospect might have been predicted from the structures of dsDNA integration complexes of Cas1/Cas2 from CRISPR systems that lack an associated RT.^25,34,54^ These structures showed the Cas1/Cas2 proteins form a hexameric complex with two Cas1 homodimers on either side separated by a Cas2 homodimer. The latter forms a platform for binding a DNA duplex across the length of the complex via non-sequence specific base and phosphate-backbone interactions with splayed or overhanging single-stranded 3’-DNA ends of the two DNA strands inserted into Cas1 active sites on opposite sides of the complex. The findings that Mm RT-Cas1/Cas2 initiates cDNA synthesis on dA-tailed RNAs at 3’-proximal sites within the RNA segment (Figure 5C) and can also add a small numbers (1 or 2) non-coded nucleotides to the 3’ end of completed cDNAs (Figure 3) has the potentially desirable effect of generating RNA/cDNA duplexes with single-strand DNA overhangs on both strands that may be favorable substrates for integration by Cas1/Cas2. The inability of Mm RT-Cas1/Cas2 proteins to integrate RNA spacers without short DNA tails may reflects steric hindrance by 2’ OH groups that impede binding of 3’-terminal nucleotides at the Cas1 active sites, with this steric hindrance possibly less for the *Thiomicrospira* spp. RT-Cas1/Cas2, which was reported to inefficiently but detectably integrate RNAs lacking a 3’-dN-tail.^14,25^

Previous findings showed that Mm and *Thiomicrospira* RT-Cas1/Cas2 differ from conventional Cas1/Cas2 proteins in being able to integrate ssDNA as well as dsDNA protospacers into CRISPR arrays and that *Thiomicrospira* RT-Cas1 could by itself integrate DNA protospacers in the absence of Cas2,^12,25^ possibly reflecting structural differences in RT-Cas1 proteins that relax strict requirements for protospacer binding. Here we extended these findings by showing that Mm RT-Cas1 could efficiently integrate 3’-dN RNAs into CRISPR arrays, with integration of duplex protospacers favored at lower protospacer concentrations and single-stranded protospacers favored at higher protospacer concentrations. The favored integration of duplex protospacers at lower protospacer concentrations likely reflects that they are more stably bound than single-stranded protospacers by phosphate-backbone contacts to both DNA strands and/or rapid sequential binding of the two 3’-DNA ends to Cas1 active sites on opposite sides of the complex. The steeper concentrations dependence for integration of a single-stranded protospacer may reflect that its less constrained binding to one active site is rate-limiting at lower protospacers concentrations, but more readily overcome at higher protospacers concentrations. This concentration dependence may be a built-in mechanism that favors protospacer acquisition from abundant RNAs produced by invading pathogens as well as abundant ssDNA fragments generated by degradation of pathogen DNAs by RecBCD or other phage defense nucleases.^55–57^ Also noteworthy was that deletion of the RT domain of Mm RT1′-Cas1/Cas2 increased the rate and amplitudes of protospacer integration for RNA-dA and RNA-dA/DNA duplexes to a greater extent than similarly configured DNA protospacers (Figure 6). This finding may reflect that the RT domain impedes RNA spacer integration when not directly coupled to cDNA synthesis and suggests there may be little inherent difference in the integration efficiencies of dN-tailed RNA and DNA protospacers by the Cas1/Cas2 integrase (Figure 6C).

Possible additional mechanism for RNA protospacer acquisition include: (i) snap-back cDNA synthesis followed by RNase digestion of the single-strand RNA template loop, resulting in an RNA/cDNA heteroduplex, and (ii) RNase H-digestion of the RNA strand of the initial RNA-dA/cDNA heteroduplex followed by second-strand DNA synthesis by RT-Cas1 or a host DNA polymerase to generate a dsDNA protospacer. While neither mechanism can be completely excluded, snap-back DNA synthesis would not generate an RNA-DNA duplex with a DNA-tail at the 3’ end of the RNA strand, while RNase H digestion of the RNA strand of an initial RNA/cDNA or RNA-dN/cDNA duplex followed by second-strand DNA synthesis would result in dsDNA protospacers lacking longer (≥2 nt) non-coded nucleotide tails linked to the 3’ end of the RNA sense-strand, a feature that distinguishes the majority of spacers acquired *in vivo* by Mm WT RT-Cas1/Cas2 and other RT-Cas1/Cas2 proteins from those acquired by Mm RT1′-Cas1/Cas2 and Cas1/Cas2 proteins that acquire spacers only from DNA (Figure 1C).

The spacers acquired by WT Mm RT-Cas1/Cas2 in its native host correspond to coding regions of highly expressed protein-coding and sncRNA genes integrated in either orientation into the CRISPR array with no specific sequence requirements. enrichment for specific regions of the RNA (Figure 1C), or discrimination of host over pathogen RNAs.^12,14^ The integration of protospacers in either orientation by Cas1/Cas1 reflects that Type III CRISPR systems have no requirement for a protospacer adjacent motif (PAM), enabling double-stranded protospacers to bind the two Cas1 active sites in either orientation and for single-stranded protospacers to bind to either Cas1 active site for integration into the top or bottom strand. Ma *et al.* suggested that dsDNA protospacers with one 3’ end inserted into either strand of the CRISPR array could become fully integrated simply by coupled integration/disintegration reactions followed by transcription-coupled DNA repair.^56^ Our finding that single-stranded 3’-dN RNAs can be site-specifically integrated into a CRISPR array raises the intriguing possibility that only the terminal transferase activity RT-Cas1/Cas2 proteins is required for RNA protospacer integration (Figure 7, right side), thereby increasing the number and diversity of RNA protospacers that could be integrated in proportion to the cellular abundance. The acquisition of RNA-derived spacers in *M. mediterranea* required over-expression of RT-Cas1/Cas2 and was not seen with endogenously expressed proteins.^12^ These findings may reflect that RT-Cas1/Cas2 over-expression is activated as a last-resort when pathogen RNAs reach high copy number by evading other cellular defense mechanisms. Under those circumstances, the over-expressed, RT-Cas1/Cas2 has no compunction about integrating host-derived RNA protospacers, potentially leading to autoimmunity and altruistic cell death.

Phylogenetic analyses indicate that CRISPR-associated RTs evolved on at least three different occasions, with the largest and most ancient clade having an RT domain closely related to those of group II introns RTs, with the RTs of two smaller clades related to abortive phage infection Abi-P2 RTs or retron RTs,^64^ both of which likely evolved earlier from a group II intron RT. Phylogenetic analyses also found that RT-Cas1 fusion proteins comprise a single evolutionary-branch of CRISPR-associated RTs, but with CRISPR-associated RTs in other branches juxtaposed and transcribed in the same direction as genes encoding Cas6 and Cas1/Cas2 proteins, suggesting that a functional link between these proteins preceded the fusions.^13,14^ The phylogenetic branching patterns also suggested that fusion of RT to Cas1 occurred first, followed by more recent fusion of Cas6 and that both fusions have been horizontally transferred to different Type III and one Type I CRISPR systems, suggesting an inherent ability to function together with other Cas1/Cas2 proteins.^14^ Frequent horizontal transfer and rapid sequence divergence make it difficult to distinguish whether RT-Cas1 fusions occurred independently on multiple occasion,^64^ but if so, it would further support a selective advantage for linked functions. Analogous fusion of a primase (Pri) domain to a CRISPR-associated RT in a different lineage may likewise confer a selective advantage by facilitating functions related to spacer acquisition or processing.^12,14,65^

Mobile group II intron-encoded RTs, the likely direct or once removed ancestors of CRISPR-associated RTs, have throughout evolution exhibited remarkable flexibility to modulate the biochemical activities of their RT active site and acquire additional domain to perform cellular functions. Bacterial examples include diversity generating retroelement RTs, which preferentially mis-incorporate specific dNTPs into coding regions to enable host-phage tropism switching;^66,67^ retron RTs and abortive phage infection (abi) RTs, which using different mechanisms to synthesize ssDNAs thought to trigger phage defense mechanisms;^18,20^ and group II intron-like 4 (G2L4) RTs, which evolved to function in double-strand break repair via microhomology-mediated end joining by optimizing by optimizing the strong strand annealing activity of group II intron RTs.^22^ Other striking examples are the evolution of mobile group II introns or their close relatives into the eukaryotic RNA splicing apparatus, including the core spliceosomal protein PrP8, which evolved from a group II intron-like RT and promotes RNA splicing by interacting with snRNAs, which evolved from group II intron RNA domains, as well as closely related RTs encoded by LINE and other eukaryotic non-LTR retrotransposons, which use variations of the reverse transcription and DNA integration mechanisms adapted to eukaryotic genomes and nuclear-cytoplasmic compartmention.^68^

The flexibility and advantageous biochemical activities of group II intron-related RTs may also be useful for biotechnological applications. In addition to transcriptional recording devices,^23^ group II intron and other non-LTR-retroelement RTs that use multiple cDNA initiation mechanisms analogous to those used by RT-Cas1 may be advantageous for genome engineering methods, such as prime editing^69–72^, that could benefit from synthesizing full-length cDNAs without an added primer as well as from the high fidelity, processivity, and strand-displacement activity enabled or engineered into the distinctive conserved structural features of these RTs.^40^ Further, the finding that the addition of short DNA segments enables Mm RT-Cas1/Cas2 to site-specifically integrate RNA into a DNA genome suggests a general method for integration of RNA into genomes by other DNA integrases and recombinases.

### Limitations of study

This study reveals novel mechanisms used by an RT-Cas1/Cas2 fusion for reverse transcription and integration of RNA-derived spacers into a CRISPR array but further studies are needed to elucidate the structural basis for these mechanisms. The reverse transcription and DNA integration mechanisms elucidated here for Mm RT-Cas1/Cas2 are consistent with features of spacers acquired by RT-Cas1/Cas2 proteins in different bacteria, but further studies are needed to assess the contribution of different mechanism to spacer acquisition *in vivo*.

## ACKNOWLEGMENTS

This work was supported by NIH grant R35 GM136216 and Welch Foundation grant F-1607. We thank Rick Russell and Debamita Paul (UT Austin) for comments on the manuscript, and Andrew Fire (Stanford) for suggesting the use of the CRISPR DNA hairpin substrate. We acknowledge the Biomedical Research Computing Facility (BRCF) at The University of Texas at Austin for providing data storage and computing resources that have contributed to the research results reported within this paper.

## SUPPLEMENTAL INFORMATION

## AUTHOR CONTRIBUTIONS

G.M. and A.M.L. designed the experiments and wrote the manuscript with help from J.Y., S.K.P, and L.M. G.M. performed most of the experiments, L.M. purified and did quality control of all of proteins. S.K.P. developed and used a new method for constructing TGIRT-based DNA-seq libraries. J.Y. analyzed TGIRT-seq data and did bioinformatic analysis of spacer acquisition *in vivo* and *in vitro*. All authors contributed to the interpretation of experimental results, and all authors read and commented on the manuscript.

## DECLARATION OF INTERESTS

A.M.L. is an inventor on patents owned by the University of Texas at Austin (UTA) for non-retroviral reverse transcriptase template switching. A.M.L and G.M are inventors on patents owned by UTA for TGIRT enzymes and other stabilized group II intron reverse transcriptase fusion proteins. A.M.L., S.K.P. and G.M. are inventors on a patent owned by and a continuation patent application filed by UTA for non-LTR-retroelement reverse transcriptase and uses thereof. A.M.L. and G.M. are inventors on a patent application filed by UTA for methods and compositions for synthesis of cDNAs by RTs using multiple reverse transcription initiation mechanisms and site-specific integration of RNA into DNA genomes.

**Figure S1.** Further analysis of RT-Cas1/Cas2 terminal transferase activity and characteristics of protospacers integrated by Mm WT RT-Cas1/Cas2 or RT1′-Cas1/Cas2 *in vivo*

(A) Terminal transferase assays were done as in Figure 1, but with unlabeled DNA or RNA oligonucleotide substrates with each of the 4 possible [α-^32^P]-dNTPs. D29 and R29 are the same 29-nt DNA and RNA oligonucleotides used for terminal transferase assays in Figure 1F, and D34 and R34 are 34-nt RNA and DNA oligonucleotides corresponding to a randomly chosen spacer acquired by Mm RT-Cas1/Cas2 *in vivo*. The sequences of the R29 and R34 oligonucleotides and a schematic of the terminal transferase (TT) assays with [α-^32^P]-dNTPs are shown beneath the gel.

(B) Density plots showing the relative location of spacers acquired by Mm WT RT-Cas1/Cas2 and RTΔ-Cas1/Cas2 within the genes from which they originated. The analysis was done using the same spacer dataset as in Figure 1B. The relative positions of the spacers across the genes to which they mapped were calculated as the percentile of the mid-point of the acquired sequence in the mapped gene. Spacers derived from both coding and non-coding RNAs were used for this analysis.

(C) Dinucleotide frequencies of the first 2 non-coded nucleotides (N_1_ and N_2_) at spacer-repeat junctions. Spacers with non-coded nucleotide on only one end, assumed to be the 3’ end, were aligned to have non-coded nucleotides on the right. Frequencies of dinucleotides at positions N1 and N2 (boxed) that were significantly higher by chi-square test (p <0.001) for spacers acquired by Mm WT RT-Cas1/Cas2 than those acquired by RTΔ-Cas1/Cas2 spacers are highlighted in red and marked with an asterisk (*).

**Figure S2. Additional data for spacer acquisition assays with RNA and DNA oligonucleotides having different length 3’-dA tails**

(A) Phosphorimager scans of gels for which time courses plots are shown in Figure 2B.

(B) Phosphorimager scans of gels and plots for a repeat of the experiment of Figure 2B. The gels were analyzed with ImageQant software, and the data were fit to a single-exponential equation using Prism for calculation of k_obs_ and Amplitude (Ampl.). R^2^ values for the plots are listed in the Supplementary data file.

(C) Spacer acquisition assays for the same DNA and RNA oligonucleotides analyzed in Figure 2B but in the presence of 1 mM dATP. Reactions were done in parallel and analyzed on the same gel as those in Figure 2B. Optimal integration of shorter RNA spacers into the CRISPR array required extension to 33 nt (brackets in gel) by Mm RT-Cas1/Cas2 terminal addition of 3’-dA tails. A schematic of the assay is shown to the right.

**Figure S3. Gel analysis of labeled cDNAs synthesized by Mm RT-Cas1/Cas2 from different templates without an added primer**

(A) cDNA synthesis from D50CCC and R50CCC templates with different 3’-blocking groups. Reactions were done at described in Methods by incubating Mm WT RT-Cas1/Cas2 (500 nM) and 250 nM D50CCC or R50CCC templates having 3’ dideoxy (ddC), phosphate (P) or Inverted dT (INVdT) 3’-blocking groups in reaction medium containing and 83 nM [α-^32^P]-dNTP and an equimolar mix of all 4 unlabeled dNTPs (500 µM each) for 1 h at 25°C. The products were analyzed on a 20% denaturing acrylamide gel. The uniform size of the products synthesized from RNA and DNA templates with a 3’-phosphate or 3’-blocking groups indicates that initiation of cDNA synthesis at 3’ proximal sites is not dependent upon a 3’ OH.

(B) Gel analysis of cDNAs synthesized from RNA and DNA templates with lower or higher dCTP concentrations. Reactions were done as described in Methods by incubating RT-Cas1/Cas2 (500 nM) with 250 nM 3’-dCC or 3’-P blocked D50NNN or R50NNN templates with AAA, CCC, GGG, UUU or CGC substituted at the location of 3’-proximal CCC initiation site (nucleotides 46 to 48) in the absence or presence of 1 mM MnCl_2_. In the left panel, reactions were done with [α-^32^P]-dCTP (20 µM) and 500 µM each other 3 dNTPs, and in the right panel, reactions were done with 83 nM [α-^32^P]-dNTP and an equimolar mix of all 4 unlabeled dNTPs (500 µM each). Black arrows indicate the cDNAs initiated at the GGG and CGC sequences in the R50GGG and R50CGC templates, respectively, visible at the high dCTP concentration (right side of right panel) but not or barely at the low dCTP concentration (left panel). Markers were 5’ labeled 10-nt ladder (Invitrogen, left) or low molecular weight DNA ladder (New England Biolabs, right).

**Figure S4.** Time courses for protein priming and synthesis of labeled cDNAs and short DNA oligomers synthesized by Mm RT-Cas1/Cas2 from R50CCC_ddC template in the absence of an added primer and comparison of rates of cDNA synthesis from the same template in the absence or presence of a dinucleotide dG2 primer

(A) The gel shows a time course for labeling of RT-Cas1 with [α-^32^P]-dGTP. Reactions were done as in Figure 4 with Mm WT RT-Cas1/Cas2 (500 nM), R50CCC_ddC template (250 nM), [α-^32^P]-dGTP (73 nM) in the presence of 1 mM MnCl_2_ at 25°C. Samples were taken from 0 to 90 min and the reactions stopped with EDTA (25 mM). An aliquot of each sample was mixed with SDS-PAGE loading dye and analyzed on an SDS 4-20% polyacrylamide gradient gel (BioRad) with Precision Plus Marker (BioRad) in a parallel lane (M) of the same gel. A phosphorimager scan is shown above, and the Coomassie blue-stained gel is shown below. Cas2 is present in too low concentrations to be visible in the Coomassie blue stained gel under these conditions.

(B) Time courses for labeling of RT-Cas1 and synthesis of DNA products. Four different reactions were done as in Figure 4 with Mm WT RT-Cas1/Cas2 (500 nM), R50CCC_ddC template (250 nM) and 1 mM MnCl_2_ at 25°C. The 4 reactions contained [α-^32^P]-dGTP (73 nM), [α-^32^P]-dGTP (73 nM) plus dGTP (500 µM), [α-^32^P]-dGTP (73 nM) plus and equimolar mix of all 4 dNTPs (500 µM each), or [α-^32^P]-dGTP (73 nM) and dGTP (500 µM) plus ddGTP (1 mM). Aliquots of the samples were incubated with or without protease K and analyzed on a denaturing 20% polyacrylamide/8 M urea gel, which was dried and scanned with a phosphorimager. Size markers were a homemade mixture of ^32^P-labeled oligonucleotides and dGMP_2_ and dGMP_3_ run in adjacent lanes (M) of the same gel. Blue arrows indicate bands that appear after proteinase K treatment.

(C) Comparison of rates of cDNA synthesis from the R50CCC_dCC template with and without added dinucleotide (dG_2_) primer. RT-Cas1/Cas2 (500 nM) was incubated with 3’-blocked R50CCC template in the presence of [α-^32^P]-dCTP (83 nM) and 500 µM dATP, dGTP, and dTTP for the indicated times at 25°C in the presence or absence of 20 µM dG_2_ primer and 1 mM MnCl_2_. The products were analyzed in 20% polyacrylamide/8 M urea gels (left), which were dried and scanned with a phosphorimager. The plots to the right show the rate of synthesis of the ∼50-nt cDNA product quantitated with ImageQuant and fit to a single exponential rate equation using Prism 10. Rates (k_obs_) and Amplitudes (Ampl.) calculated from the curve fits are shown above the plots, and R^2^ values for the plots are listed in the Supplementary data file.

**Figure S5.** Gel analysis of ^32^P-labeled cDNA synthesized from 3’-blocked RNA templates with different length RNA and 3’-dA tail segments Templates were derivatives of the 50 nt R50CCC oligonucleotide with the indicated number of dA-residues added to their 3’ end. For the group on the left, the 3’-dA tail was kept at a constant 4 nt while varying the length of the RNA segment from 29 to 32 nt. For the group in the middle, the length of the RNA segment was kept constant at 30 nt while the length of the 3’-dA tail varied from 1 to 5 nt. For the group on the right, the overall length of the template was kept constant at 34 nt while the length of the RNA segment and 3’-dA tail varied as indicated by the name of the oligonucleotide.

(B Templates were derivatives the R29 oligonucleotide with different length dA-tails. Reverse transcription reactions were done with 500 nM RT-Cas1/Cas2, 250 nM template oligonucleotide, [α-^32^P]-dCTP (19 µM) and 500 µM dATP, dGTP, and dTTP, and ± 1 mM MnCl_2_, and the products were analyzed on a denaturing 20% polyacrylamide/8 M urea gel, which was dried and scanned with a phosphorimager. cDNA start sites from parallel reactions with the same templates with unlabeled dNTPs are shown in Figure 5C.

**Figure S6.** RNase H control experiment confirming integration of a stable RNA-dN/cDNA duplex into a CRISPR array and time courses for all ^32^P-labeled products resulting from cleavage-ligation reactions at single or both sites on the top and bottom strands of the CRISPR DNA in the RNA protospacer integration assays of Figure 6B.

(A) RNase H control. cDNAs were synthesized by incubating 5’-labeled (*) R29+dA2 or R29+dA2_ddC protospacers (250 nM) with RT-Cas1/Cas2 (500 nM) and dNTPs (100 µM) in the presence or absence of MnCl_2_ (30 µM) and a dinucleotide dG_2_ or dT_2_ primer (20 µM) for 1 h at 25°C. After clean up with a Zymo Oligo Clean and Concentrator kit, the products of the reverse transcription reactions were used for protospacer integration assays by incubating them with Mm RT-Cas1/Cas2 (500 nM) and CRISPR DNA hairpin DNA substrate (100 nM) for 1 h at 25°C. After addition of EDTA to a final concentration of 25 mM, the reaction products were treated with Protease K (0.32 unit, 15 min, 37°C), cleaned up with a Zymo Oligo Clean and Concentration kit, incubated with or without RNase H (12.5 unit; New England Biolabs) for 15 min at 37°C, and analyzed on an 8% polyacrylamide/8 M urea gel against a 5’-labeled New England Biolabs low molecular weight DNA ladder run in a parallel lane (M). The Figure shows phosphorimager scans of the dried gel at two different exposures (top and bottom panels) in order to better visualize the residual 5’-labeled RNA substrate that was not digested by RNase H (bottom panel). Arrows point to reaction products that showed significantly lower labeling after RNase H treatment, indicating that they were in RNA-dN/cDNA duplexes that stably integrated into the CRISPR array as shown in the schematic above of the gel. The lighter product bands for the 3’-blocked R29+dA2_ddC RNA template/protospacer were not seen in the absence of dNTPs, suggesting that they reflect addition of 3’-dN tails to a fraction of RNA protospacers that lacked or lost the 3’-ddC blocking group.

(B-D) Time courses for all ^32^P-labeled products the RNA protospacer integration assays of Figure 6B. The three different ^32^P-labeled protospacer configurations are shown schematically at the top of each set of time courses. For each protospacer configuration, assays were done with single-stranded *R29+dAn (top left), single-stranded *D29+dAn (bottom left), double-stranded *R29+dAn/D29+dAn (top right), and double-stranded R29+dAn/*D29+dAn (bottom right), with t indicating the top strand, b indicating the reverse complement of the top strand, and * indicating the 5’-labeled strand. Symbols and color codes for the 3 labeled products are shown below panel D. The plots for the single-strand protospacer reactions (left) show that the product for cleavage-ligation at the bottom-strand site (red open triangles) occurs rapidly and plateaus, while the product for cleavage-ligation at the top-strand site (red open squares) accumulates more slowly and then declines in parallel with accumulation the two-site cleavage ligation product (black open circles), as expected for an intermediate rate-limiting step. By contrast, the plots for the duplex protospacers reactions (right) show that the two-site cleavage-ligation product (black closed circles) accumulates faster and to a higher amplitude than the intermediate top-strand cleavage-ligation product for the two R29-based protospacers, as expected for more rapid sequential ligation of both strands of the duplex. For the R30CCC protospacer, the two-site cleavage-ligation product accumulates more slowly but eventually to higher amplitude than both intermediate single site cleavage-ligation reaction products, possibly reflecting slower but ultimately more efficient sequential integrations.

**Figure S7.** Time courses showing rates of integration of single-stranded and duplex DNA protospacers having the same configurations and sequences as those for the RNA-dN and RNA-dN/cDNA duplexes in Figure 6B and C.

(A, B) Time courses for spacer integration reactions done in parallel with those in Figure 6B and 6C. The plots show time courses for production the labeled 99-nt band resulting from cleavage-ligation reactions at the 5’ end of R1 on both strands. The data were fit to a single exponential equation for calculation of rate (k_obs_) and amplitudes (Ampl). t or b indicate the top or bottom strands of the duplex DNA spacers, respectively, with * indicating the 5’-labeled strand. Orange, 5’-labeled DNA-dA strand; blue, 5’-labeled complementary DNA strand; open symbols, single-stranded RNA or DNA; closed symbols; double-stranded protospacers. Symbols and schematics for the labeled protospacers are shown below the plots in panel B.

(C) Time courses for all ^32^P-labeled products resulting from cleavage-ligation reactions at single sites or both sites on the top and bottom strands of the CRISPR DNA for the DNA protospacers iin panel A. Symbols and color codes for the 3 labeled products are shown at the bottom.

## METHODS

### Bioinformatic analysis of spacer acquired *in vivo*

Spacer sequences were obtained from the following sources: published results (Mm (MMB1) WT and RT1′-Cas1/Cas2) or SRA depository (Fs: SRR8102160, 163, 166-8, 170, 172-7, 194, 207; Vv: SRR8962131-134,137,138; Tt: SRR11818505-8, SRR12227011-4; Se: SRR16249482-3; St: SRR1595171-3, 176,187,198,209).

Raw reads were downloaded from the SRA depository and merged if the dataset was comprised of pair-end reads by using fastq-join (https://github.com/brwnj/fastq-join). Spacers were identified as sequences between 2 repeats. Candidate spacers were mapped to the genome of the bacterial strain from which they were obtained (Mm: MMB-1 NC_015276.1; Fs: *E. coli* BL21 (DE3) NC_012947.1; Vv: *E. coli* HMS174 (DE3) NZ_LM993812.1; Tt: Tt phage phiFa/KO MH673671.2 and MH673672.2; Se: *S. aureus* RN4220 NZ_CP076105.1; St: Sth JIM8232 FR875178.1) by using

Bowtie2 (https://github.com/BenLangmead/bowtie2 with the following parameters: --very-sensitive-local. After mapping, duplicate spacer sequences were removed by samtools using the parameter markdup -r. Spacers with unique sequences were then analyzed by using customized bash and R scripts to identify non-encoded nucleotides and nucleotide frequencies at different position. Density plots for spacer length distribution were plotted in R.

The frequencies of the first 5 non-coded nucleotides (Figure 1D) and di-nucleotides frequency of the first two non-encoded nucleotides (Figure S1C) were calculated for acquired spacers with non-encoded nucleotide at one end only, assume to the 3’ end. Spacer sequences were rearranged to have the same orientation with nucleotides derived from encoded region on the left (5’ end) and non-encoded region on the right (3’ end) as shown in the Figures. The distribution of spacer sequences across the gene from which they were originated were calculated as the percentile of the midpoint of the mapped feature from which they were acquired (host, viral) protein-coding and non-coding RNAs.

### Strains and culture conditions

*E. coli* DH5α (F^−^ *endA1 glnV44 thi-1 recA1 relA1 gyrA96 deoR nupG purB20* φ80d*lacZ*ΔM15 Δ(*lacZYA-argF*) U169 hsdR17(*r_K_*_–_*m_K_*_+_) λ^−^) was used for cloning and *E. coli* Rosetta2 (DE3) (F^-^ *ompT hsdS*_B_(r_B-_ m_B-_) *gal dcm* (DE3) pRARE2 (Cam^R^)), Novagen) was used for protein expression.

### DNA and RNA constructs

Mm WT RT-Cas1 and RTΔ-Cas1 were expressed from plasmids pMalRF-RT-Cas1-cHis and pMalRF-RTΔ-Cas1-cHis, repspectively.^12,14^ The expressed proteins have a maltose-binding protein (MBP) tag fused to their N-terminus via a non-cleavable rigid linker to increase their stability in the absence of bound nucleic acids,^73^ and a C-terminal 8x His tag. The amino acid at position 2 of the Cas6-RT-Cas1 domain was changed from leucine to valine to accommodate the MBP fusion.

Plasmid CassetteAv2_pBAD contains the CRISPR03 array cloned into the pBAD/Myc–His B backbone (Life Technologies).^12^ It was used to prepare CRISPR DNA substrates by PCR with primers MMB1Lead40-5 and MMB1crisp3-r1 using Phusion High-Fidelity DNA polymerase according to the manufacturer’s protocol (New England Biolabs or ThermoFisher). The resulting 88-bp PCR product has a 40-bp leader, 35-bp repeat 1, and 13 bp of spacer 1. The substrate was labeled by PCR with 20 µCi [α-^32^P]-dCTP (Revvity), 120 µM dCTP, and 600 µM of the other three dNTPs. The labeled DNA was purified by electrophoresis in a non-denaturing 6% polyacrylamide gel, cutting out the labeled band, and electro-eluting the DNA using midi D-Tube dialyzer cartridges (Novagen). The eluted DNA was concentrated by butanol extraction, purified by using an Zymo Oligo Clean and Concentrator kit, and quantitated by using a [α-^32^P]-dCTP standard curve in a Beckman LS6500 liquid scintillation counter.

pET9Cas2, used to express Cas2 protein without an added tag, was constructed by PCR amplifying the Cas2 coding region from pCassetteAv2_pBAD (see above), using Cas2_pet_5 and Cas2_pet_3, and cloning the PCR product between the NdeI and BamHI sites of pET9a (EMD Millipore). PCR-amplified DNA sequences were confirmed by Sanger sequencing.

A minimal 68-bp CRISPR DNA was prepared by annealing complementary oligonucleotides L20R1top and L20R1bot, with biotin moieties attached to their 3’ ends. 10 µM of each oligonucleotide was mixed, heated to 82°C for 2 min and slowly cooled to room temperature. The 68-bp CRISPR DNA contains a 20-bp leader, 35-bp repeat, and 13 bp of spacer 1. A hairpin variant (HP-3) of the 68-nt CRISPR DNA was made by connecting the 3’ end of the top strand to the 5’ end of the bottom strand by a 5-nt linker (5’-TACAT).

### Radiolabeling of protospacers

Oligonucleotides were 5’ end-labeled in 50-µL reactions containing 1 nmol oligonucleotide, 600 µCi [γ-^32^P]-ATP (6,000 Ci/mmol, Revvity), 1 nmol ATP, 40 units T4 polynucleotide kinase (New England Biolabs) for 45 min at 37°C in the manufacturer’s buffer. Labeled oligonucleotides were purified in a denaturing 10% polyacrylamide gel. The gel area containing the oligonucleotide was cut out, crushed, and incubated with 1 mL of TE (10 mM Tris-HCl, pH 7.5, 1 mM EDTA) per ∼1.5 cm of gel slice at 4°C overnight on a rotator. The supernatant was removed, and the gel pieces were washed with an equal volume of TE. The combined supernatants were extracted repeatedly with butanol until the volume was ≤200 µL. After clean up with a Zymo Oligo Clean and Concentrator kit, the labeled oligonucleotide was eluted in 40 to 60 µL H_2_O and quantitated against a [γ-^32^P]-ATP standard curve using a LS 6500 scintillation counter (Beckman-Coulter).

### Protein expression and purification

Protein expression plasmids were transformed into *E. coli* Rosetta2 (EMD Millipore) and plated on LB plates containing Ampicillin (Amp, 100 µg/mL) and Chloramphenicol (Cap, 25 µg/mL). Single transformed colonies were inoculated in 100-mL LB medium supplemented with Amp and Cap in a 250-mL Erlenmeyer flask and incubated overnight at 37°C with shaking. For preparation of Mm RT-Cas1 proteins, six 4-liter flasks with 1 L LB broth were each inoculated with 10 mL of the overnight culture and grown at 37°C with shaking to log phase (O.D._600_ 0.6 to 0.8). IPTG was then added to a final concentration of 1 mM, and the cultures were incubated at 19°C for 20 to 24 h. Cells were harvested by centrifugation (JLA-8.1 rotor, 20 min, 5,000 x g), and the pelleted cells were resuspended in 10 mL/g cells of A1 buffer (25 mM Tris-HCl pH 7.5, 500 mM NaCl, 10% glycerol, 10 mM ß-mercaptoethanol (BME)) on ice. The cells were then incubated with lysozyme (1 mg/mL, 30 min, 4°C) and sonicated (Branson Sonifier 450; 3 bursts of 15 sec each with 15 sec between each burst). The lysate was cleared by centrifugation (Beckman-Coulter JA-14 rotor; 29,400 × g, 25 min, 4°C), and polyethyleneimine (PEI) was added to the supernatant with stirring on ice in six steps to a final concentration of 0.4%. After 10 min, precipitated nucleic acids were removed by centrifugation (29,400 × g, 25 min, 4°C), and proteins were precipitated from the supernatant by adding ammonium sulfate to 60% saturation and incubating on ice for 30 min.

Proteins were collected by centrifugation (Beckman-Coulter JA-14 rotor; 29,400 × g, 25 min, 4°C), dissolved in 20 mL A1 buffer, and filtered through a 0.45-µm polyethersulfone membrane (Whatman Puradisc). Protein purifications were done by using an ÄKTA start system (Cytiva). RT-Cas1 proteins were purified by loading the filtered crude protein onto an amylose column (5 ml; MBPTrap HP; Cytiva), washing with 50 mL of A1 buffer, followed sequentially by 30 mL of A1 plus 1.5 M NaCl and 30 mL of A1 buffer. Bound proteins were eluted with 50 mL of 10 mM maltose in A1 buffer. Fractions containing RT-Cas1 were identified by SDS-PAGE and pooled. The protein was then loaded onto a Nickel column (5-mL, HisTrap FF column; Cytiva) and eluted with a 100 mL 25 to 100 mM imidazole gradient. Peak fractions were identified by SDS-PAGE, pooled, and dialyzed into A1 buffer (10 MWCO; SnakeSkin™ Dialysis Tubing; Thermo Fisher Scientific). The dialyzed protein was concentrated to >10 µM by using an 30K Pall concentrator and stored at −80°C.

The initial steps in the preparation of Cas2 expressed from pET9-Cas2 were similar, except that the cell paste was resuspended in H1 buffer (25 mM Tris-HCl pH 7.5, 100 mM KCl, 10% glycerol, 10 mM dithiothreitol (DTT)), and ammonium sulfate precipitation was done at 40% saturation. The Cas2 containing supernatant after ammonium sulfate precipitation was loaded directly onto a heparin-Sepharose column (5ml; HiTrap Heparin HP column; Cytiva). The protein was eluted with a linear 100 mM to 1 M KCl gradient, and Cas2 peak fractions (∼800 mM KCl) identified by SDS-PAGE were frozen in elution buffer for storage overnight. The peak fractions were thawed, diluted to about 125 mM KCl, and loaded onto a second 5-mL heparin-Sepharose column. Cas 2 protein was eluted with a 100 to 400 mM KCl gradient, followed sequentially by extended washes with 400 mM KCl and a shallow elution gradient from 400 mM to 1 M KCl to separate Cas2 from contaminating nucleases. Peak fractions were identified by SDS-PAGE, dialyzed against H1 buffer, and loaded onto a SP column (5-mL, HiTrap SP HP column; Cytiva). Proteins were eluted with a 100 mM to 1 M KCl linear gradient. Peak fractions were identified by SDS-PAGE and stored frozen at −80°C.

Protein concentrations were measured by using a Qubit Protein assay kit (Life Technologies) according to the manufacturer’s protocol. Nucleic acid contamination was determined by using Qubit ssDNA, dsDNA and ssRNA assay kits. ssDNA was below 1 ng/mL purified protein preparation and RNAs and dsDNA were undetectable. All Proteins were >90% pure as assayed by SDS-PAGE.

### Terminal transferase assays

Purified RT-Cas1-8xHis protein (2 µM) was mixed with purified Cas2 (2 µM) in reaction medium containing 100 mM KCl, 100 mM NaCl, and 25 mM Tris-HCl pH 7.5, 10 mM MgCl_2_, 1 mM DTT, 1 mM BME, and 2% glycerol and incubated at room temperature for 5 min before placing on ice prior to use. Assays were done in 20 µL of 25 mM Tris-HCl, pH 7.5, 10 mM free MgCl_2_, 5’-labeled DNA or RNA oligonucleotides (100 nM), and 1 mM of a single dNTP with 1 mM MnCl_2_ added where indicated. Reactions were initiated by adding 5 µL of protein complex. Terminal transferase assays with unlabeled oligonucleotides substrates were done in reaction medium containing 83 nM [α-^32^P]-dNTP (Revvity) and 250 µM of unlabeled dNTP added with an equimolar amount of MgCl_2_.

Reactions were incubated at 37°C for 1 h and stopped by adding phenol-CIA-Isoamyl alcohol (25:24:1). The supernatant was mixed at a 2:1 ratio with loading dye (90% formamide, 20 mM EDTA, and 0.25 mg/ml bromophenol blue and xylene cyanol), and nucleic acids were analyzed on a denaturing 8% polyacrylamide/8 M urea gel. The gels were dried and scanned with a phosphorimager (Typhoon, Cytiva). Molecular weight markers were a 5’-labeled 10-nt ssDNA ladder (Invitrogen) run in a parallel lane.

### Spacer integration assays

Duplex protospacers, consisting of a top strand 29- or 30-nt RNA or DNA oligo having 3’ dA_n_ tails and a complementary DNA oligonucleotide also with 3’ dA_n_ tails, were used for spacer integration assays. The oligonucleotides were annealed at 2.5 µM concentration each in 20 µL of 25 mM Tris-HCl pH 7.5, 20 mM NaCl by heating to 94°C for 30 s and slowly cooling (0.1°C/s) to 25°C followed by further incubation for 5 min at 25°C. The annealed oligonucleotides were then placed on ice prior to use. Analysis of the annealed protospacers on a non-denaturing 15% polyacrylamide gel showed that >95% were double-stranded. Purified MBP-Cas6-RT-Cas1-8xHis protein was mixed with purified Cas2 in reaction medium containing 100 mM KCl, 100 mM NaCl, and 25 mM Tris-HCl pH 7.5, 10 mM MgCl_2_, 1 mM DTT, 1 mM BME, and 2% glycerol and incubated at room temperature for 5 min and also placed on ice prior to use. Spacer integration reactions contained RT-Cas1/Cas2 complex (500 nM final), MMB-1 CRISPR DNA (5 to 100 nM), 20 mM Tris-HCl, pH 7.5, and 10 mM free MgCl_2_, single-strand or duplex DNA or RNA oligonucleotides (250 nM final concentration) plus an equimolar solution of dNTPs with MnCl_2_ (1 mM final concentration) added where indicated. Total volume for each time course is either 250 µL when taking 20 µL samples/time point or 120 µL when taking 10 µL samples. Up to 8 time course reactions are run in parallel in a PCR machine. Reactions were initiated by adding the Cas1/Cas2 protein complex, incubated at 37°C for 1 h, and stopped by adding phenol-CIA (Figure 2A and B) or EDTA (25 mM final concentration) and 1.8-units Proteinase K, and incubating at 25°C for 15 min (all other reactions). The supernatant was mixed at a 2:1 ratio with loading dye (90% formamide, 20 mM EDTA, and 0.25 mg/mL bromophenol blue and xylene cyanol), and nucleic acids were analyzed on a denaturing 6% polyacrylamide/8 M urea gel. Gels were dried and scanned with a phosphorimager. Molecular weight markers were a 5*’*-labeled 10-nt ssDNA ladder (Invitrogen) run in a parallel lane.

Time course gels using the hair-pin CRISPR DNA were quantitated by exposing the autoradiogram so that none of the bands were saturated upon scanning. Gels were analyzed in ImageQuant (Cytiva) by boxing labeled bands along with a background control above or below the labeled band. The fraction of product was determined from the counts in the labeled band minus the background relative to the total counts of all labeled products in the lane. The data for the largest band (154 nt) which is an intermediate that has a protospacer inserted into the top strand was fit to the following equation: Y=[(a*(k1))/((k2-k1))]*[exp((-k1*t))-exp((-k2*t))] where t=time and k1 and k2 indicate the rate constants for production and destruction of the intermediate. The other two bands were fit to a single exponential rate equation. R^2^ values for all curve fits are shown in the Supplementary data file.

### cDNA synthesis assays

Reactions were done with Mm WT RT-Cas1/Cas2 protein (500 nM final concentration), 25 mM Tris-HCl pH 7.5, and 10 mM free MgCl_2_, and DNA or RNA template oligonucleotides (250 nM). dNTPs were added at 500 nM or 1 mM together with an equimolar amount of MgCl_2_. For the assays in Figure 5A, dinucleotide primers were 5’-labeled using T4 polynucleotide kinase (New England Biolabs) and added at 20 µM without pre-annealing to the template. cDNA synthesis assays with unlabeled primers or without primers were done with a dNTP mix containing 83 nM [α-^32^P]-dNTP (3000 Ci/mmol, Revvity) and 20 µM of the same unlabeled nucleotide and 500 µM of the other 3 dNTPs or 83 nM [α-^32^P]-dNTP (3,000 Ci/mmol) plus 500 µM of each dNTP, as specified for individual experiments. MnCl_2_ was added where indicated at 1 mM final concentration. Reactions were incubated at 25°C for 1 h and stopped by adding phenol-CIA (Figures 1, 2, S1, and S2) or EDTA (25 mM final concentration) plus 1.8-units Protease K (all other Figures, New England Biolabs), and incubating at 25°C for 15 min. The extracted nucleic acids were mixed at a 2:1 ratio with loading dye (90% formamide, 20 mM EDTA, and 0.25 mg/ml bromophenol blue and xylene cyanol) and analyzed in a denaturing 20% polyacrylamide/8 M urea gel, which was dried and scanned with a phosphorimager. Molecular weight markers were a 5’-labeled 10-nt ssDNA ladder (Invitrogen) run in a parallel lane.

In order to determine if RT-Cas1/Cas2 produces a stable RNA/cDNA, DNA synthesis reactions in the absence of primer were done as described above using the D/R50CCC-ddC template oligos. The reactions were scaled up to 40 µL and stopped by adding EDTA (25 mM final) and Protease K (1.8 units). The cDNA products were purified by using a Zymo Oligo Clean and Concentrator kit and incubated for 20 min at 37°C in the presence or absence of RNase H or RNase A (New England Biolabs) at a low salt concentration followed by Protease K digestion (as above). Samples were analyzed in a non-denaturing 15% polyacrylamide gel which was dried and analyzed with a phosphorimager.

### Protein-priming assays

Purified RT-Cas1 protein was mixed with purified Cas2 (2 µM each) in reaction medium containing 100 mM KCl, 100 mM NaCl, and 25 mM Tris-HCl pH 7.5, 10 mM MgCl_2_, 1 mM DTT, 1 mM BME, and 2% glycerol and incubated at room temperature for 5 min before placing on ice until ready for use. Reactions were done with 500 nM RT-Cas1/Cas2 in the absence or presence of 250 nM DNA or RNA oligonucleotide templates in reaction medium containing 25 mM Tris-HCl, pH 7.5 and 10 mM free MgCl_2_, with 1 mM MnCl_2_ added where indicated. Reactions were initiated by adding 83 nM [α-^32^P]-dNTP (3000 Ci/mmol, Revvity) in 25 mM Tris-HCl pH 7.5, 10 mM MgCl_2_ and incubated at 25°C for 15 min. Some reactions were chased by adding 500 µM of a single dNTP, all 4 dNTPs or 25 mM Tris-HCl pH 7.5, 10 mM MgCl_2_ buffer as a control followed by further incubation at 25°C for 30 min. Some samples were treated with protease K (1.8 units; New England Biolabs), micrococcal nuclease (MNase; 4×10^5^ units; New England Biolabs) or RNase A (2 µg, Monarch RNase A; New England Biolabs) and incubating for another 15 min at 25°C. The reactions were stopped by adding 4x NuPAGE Sample Buffer (Invitrogen) or 4x Laemmli Buffer (BioRad) to a final 1x concentration and heating to 90°C for 3 min. Samples were analyzed on a 4-15% polyacrylamide NuPage Bis-Tris gel (invitrogen) in 1x morpholineethanesulfonic acid (MES) SDS running buffer (invitrogen) or on a 4-20% polyacrylamide Criterion gels (BioRad) in 1x Tris/glycine/SDS buffer (BioRad) with Precision Plus Protein standards (BioRad) as size markers. Protein gels were stained with Coomassie blue as described^74^, dried, and scanned with a phosphorimager.

### Thermostable Group II Intron Reverse Transcriptase sequencing (TGIRT-seq) of cDNAs synthesized by Mm RT-Cas1/Cas2

Mm RT-Cas1/Cas2 reverse transcription reactions used to synthesize cDNAs for sequencing were done as above with 500 nM RT-Cas1/Cas2, 250 nM RNA template and 1 mM dNTPs without or with 20 µM dinucleotide DNA primers in 160 µL of reaction medium (see above) for 1 h at 25°C. The reaction was stopped by adding 20-µL stop solution (20 mM Tris-HCl pH 7.5, 200 mM EDTA) with or without 0.8 µg/µL RNase A as indicated and incubated for 15 min at 37°C followed by digestion with Protease K (20 µL, 0.4 unit/µL) for 15 min at 37°C. The products were cleaned-up with a Zymo Oligo Clean & Concentrator Kit. Nucleic acid concentrations were measured with a Qubit ssDNA Assay kit (Invitrogen) according to the manufacturer’s protocols, and 10-20 ng of cDNA product from each reaction was used for library preparation.

For construction of TGIRT-seq libraries, first-strand DNA synthesis was done by TGIRT template switching using a 34-nt RNA Illumina R2 adapter annealed to a complementary 35-nt R2 (R2R) DNA leaving a single nucleotide 3’ DNA overhang (an equimolar mixture of A, T, G and C) that can base pair to the 3’ nucleotide of the cDNA products as described.^75,76^ The reaction was stopped by adding proteinase K (1 unit; New England Biolabs) and 25 mM EDTA followed by clean-up with a Monarch PCR & DNA Cleanup Kit (New England Biolabs). Second-strand DNA synthesis was done by ligating 10 µM of a blunt-end duplex comprised of a 32-bp Illumina R1-3’SpC3 and 5’-phosphorylated R1R-3’SpC3 DNA (pre-annealed by incubating at 95°C for 3 min and slowly cooling to 25°C) using a Quick ligase kit (New England Biolabs) according to manufacturer’s protocol followed by clean-up as described above. The ligated dsDNA products were amplified by PCR using Phusion High-Fidelity PCR Master Mix with HF Buffer (New England Biolabs) with 200 nM of Illumina multiplex and index barcode primers (98°C for 10 sec pre-denaturation followed by 15 cycles of 98°C for 5 sec, 60°C 10 sec, 72°C 15 sec). The resulting cDNA libraries were cleaned up by using 1.4X AMPure XP beads (Beckman Coulter) and eluted in 25 µL double-distilled H_2_O. One µL of the library was analyzed on an Agilent 2100 Bioanalyzer using a High Sensitivity DNA chip (Agilent) to assess product profiles and concentrations, and the remainder was sequenced on an Illumina Nextseq 550 to obtain ∼1 million 2 x 75 nt paired-end reads per sample at the Genome Sequencing and Analysis Facility (GSAF) at the University of Texas at Austin.

Illumina TruSeq adapters and PCR primer sequences were trimmed from the reads with Cutadapt v3.5 ((https://github.com/marcelm/cutadapt) sequencing quality score cut-off at 20; p-value <0.01) and reads <15-nt after trimming were discarded. Pair-ended reads were merged by using fastq-join. The merged reads were then mapped to the reverse complement of the corresponding template sequences using Bowtie2 with the parameters --very-sensitive-local -L 5. Mapped reads with 5 or fewer soft-clipped nucleotides were retrieved and re-aligned to the reverse complement of the template sequence using MAFFT (Multiple alignment using fast Fourier transform, https://github.com/GSLBiotech/mafft) with the following settings: --auto --addfragments -reorder --keeplength –preservecase.^77^ Mapping of cDNA start sites onto the template sequence, stacked-bar graphs of nucleotide frequencies, and sequence logos of reads comprising ≥1% of all reads were made using R.

For analysis of snap-back DNA synthesis products, reads containing the full-length template sequence linked to the 5’ end of a cDNA were collected and RNA template sequences were trimmed by using Cutadapt v3.5 with default settings. Trimmed reads were then mapped to the reverse complement of the corresponding template sequences using Bowtie2 with the parameters--very-sensitive-local −L 5. Mapped reads were retrieved and re-aligned to the reverse complement of the template sequence by using MAFFT with the following settings: --auto --addfragments -- reorder --keeplength –preservecase. Products comprising ≥5% of all snap-back products were plotted against the corresponding template sequences using Illustrator (Adobe).

### Quantification and Statistical Analysis

Products of terminal transferase, spacer acquisition, and cDNA synthesis assays were quantitated using ImageQuant TL ver. 8.1 (General Electric). Excel ver. 16 (Microsoft) was used to determine mean, median, and standard deviation values. Prism 9.0 (Graphpad Software) was used for curve fitting of quantitated assays to determine k_obs_ and Amplitude. R (v4.0.3) package ggplot2 was used to generate sequence logos from high-throughput sequencing data.

### Data and software availability

TGIRT-seq dataset used in this study were deposited in the SRA with the accession number PRJNA1003443. Scripts used in analysis of spacers, *de novo* cDNAs, and snap-back products have been deposited in Github (https://github.com/reykeryao/RTCas1).

## REFERENCES

1. Koonin, E.V., and Makarova, K.S. (2022). Evolutionary plasticity and functional versatility of CRISPR systems. Plos Biol. 20, e3001481. 10.1371/journal.pbio.3001481.

2. Brouns, S.J.J., Jore, M.M., Lundgren, M., Westra, E.R., Slijkhuis, R.J.H., Snijders, A.P.L., Dickman, M.J., Makarova, K.S., Koonin, E.V., and Oost, J. van der (2008). Small CRISPR RNAs Guide Antiviral Defense in Prokaryotes. Science 321, 960–964. 10.1126/science.1159689.

3. Terns, M.P., and Terns, R.M. (2011). CRISPR-based adaptive immune systems. Curr. Opin. Microbiol. 14, 321–327. 10.1016/j.mib.2011.03.005.

4. Jiang, F., and Doudna, J.A. (2015). The structural biology of CRISPR-Cas systems. Curr. Opin. Struct. Biol. 30C, 100–111. 10.1016/j.sbi.2015.02.002.

5. Barrangou, R., and Horvath, P. (2017). A decade of discovery: CRISPR functions and applications. Nature Microbiol. 2, 17092. 10.1038/nmicrobiol.2017.92.

6. Liu, T.Y., and Doudna, J.A. (2020). Chemistry of Class 1 CRISPR-Cas effectors: Binding, editing, and regulation. J. Biol. Chem. 295, 14473–14487. 10.1074/jbc.rev120.007034.

7. Nussenzweig, P.M., and Marraffini, L.A. (2020). Molecular Mechanisms of CRISPR-Cas Immunity in Bacteria. Annu. Rev. Genet. 54, 1–28. 10.1146/annurev-genet-022120-112523.

8. Makarova, K.S., Wolf, Y.I., and Koonin, E.V. (2022). Evolutionary Classification of CRISPR-Cas Systems. In CRISPR: Biology and Applications, R. Barrangou, E. J. Sontheimer, and L. A. Marraffini, eds. (American Society for Microbiology), pp. 13–38. 10.1002/9781683673798.ch2.

9. Goldberg, G.W., Jiang, W., Bikard, D., and Marraffini, L.A. (2014). Conditional tolerance of temperate phages via transcription-dependent CRISPR-Cas targeting. Nature 514, 633–637. 10.1038/nature13637.

10. Samai, P., Pyenson, N., Jiang, W., Goldberg, G.W., Hatoum-Aslan, A., and Marraffini, L.A. (2015). Co-transcriptional DNA and RNA Cleavage during Type III CRISPR-Cas Immunity. Cell 161, 1164–1174. 10.1016/j.cell.2015.04.027.

11. Marraffini, L.A. (2022). Mechanism of Type III CRISPR-Cas Immunity. In CRISPR: Biology and Applications, S. L. A. Marraffini, ed., pp. 71–84. 10.1002/9781683673798.ch5.

12. Silas, S., Mohr, G., Sidote, D.J., Markham, L.M., Sanchez-Amat, A., Bhaya, D., Lambowitz, A.M., and Fire, A.Z. (2016). Direct CRISPR spacer acquisition from RNA by a natural reverse transcriptase-Cas1 fusion protein. Science 351, aad4234. 10.1126/science.aad4234.

13. Silas, S., Makarova, K.S., Shmakov, S., Páez-Espino, D., Mohr, G., Liu, Y., Davison, M., Roux, S., Krishnamurthy, S.R., Fu, B.X.H., et al. (2017). On the Origin of Reverse Transcriptase-Using CRISPR-Cas Systems and Their Hyperdiverse, Enigmatic Spacer Repertoires. mBio 8, e00897–17. 10.1128/mbio.00897-17.

14. Mohr, G., Silas, S., Stamos, J.L., Makarova, K.S., Markham, L.M., Yao, J., Lucas-Elío, P., Sanchez-Amat, A., Fire, A.Z., Koonin, E.V., et al. (2018). A Reverse Transcriptase-Cas1 Fusion Protein Contains a Cas6 Domain Required for Both CRISPR RNA Biogenesis and RNA Spacer Acquisition. Mol. Cell 72, 700–714.e8. 10.1016/j.molcel.2018.09.013.

15. Toro, N., Martínez-Abarca, F., and González-Delgado, A. (2017). The Reverse Transcriptases Associated with CRISPR-Cas Systems. Sci. Rep. 7, 7089. 10.1038/s41598-017-07828-y.

16. Toro, N., Martinez-Abarca, F., González-Delgado, A., and Mestre, M.R. (2018). On the Origin and Evolutionary Relationships of the Reverse Transcriptases Associated With Type III CRISPR-Cas Systems. Front. Microbiol. 9, 1792. 10.3389/fmicb.2018.01317.

17. Doulatov, S., Hodes, A., Dai, L., Mandhana, N., Liu, M., Deora, R., Simons, R.W., Zimmerly, S., and Miller, J.F. (2004). Tropism switching in Bordetella bacteriophage defines a family of diversity-generating retroelements. Nature 431, 476–481. 10.1038/nature02833.

18. Wang, C., Villion, M., Semper, C., Coros, C., Moineau, S., and Zimmerly, S. (2011). A reverse transcriptase-related protein mediates phage resistance and polymerizes untemplated DNA in vitro. Nucleic Acids Res. 39, 7620–7629. 10.1093/nar/gkr397.

19. Handa, S., Reyna, A., Wiryaman, T., and Ghosh, P. (2021). Determinants of adenine-mutagenesis in diversity-generating retroelements. Nucleic Acids Res. 49, 1033–1045. 10.1093/nar/gkaa1240.

20. Figiel, M., Gapińska, M., Czarnocki-Cieciura, M., Zajko, W., Sroka, M., Skowronek, K., and Nowotny, M. (2022). Mechanism of protein-primed template-independent DNA synthesis by Abi polymerases. Nucleic Acids Res. 50, 10026–10040. 10.1093/nar/gkac772.

21. Wang, Y., Guan, Z., Wang, C., Nie, Y., Chen, Y., Qian, Z., Cui, Y., Xu, H., Wang, Q., Zhao, F., et al. (2022). Cryo-EM structures of Escherichia coli Ec86 retron complexes reveal architecture and defence mechanism. Nat. Microbiol. 7, 1480–1489. 10.1038/s41564-022-01197-7.

22. Park, S.K., Mohr, G., Yao, J., Russell, R., and Lambowitz, A.M. (2022). Group II intron-like reverse transcriptases function in double-strand break repair. Cell 185, 3671–3688.e23. 10.1016/j.cell.2022.08.014.

23. Schmidt, F., Cherepkova, M.Y., and Platt, R.J. (2018). Transcriptional recording by CRISPR spacer acquisition from RNA. Nature 562, 380–385. 10.1038/s41586-018-0569-1.

24. González-Delgado, A., Mestre, M.R., Martínez-Abarca, F., and Toro, N. (2019). Spacer acquisition from RNA mediated by a natural reverse transcriptase-Cas1 fusion protein associated with a type III-D CRISPR–Cas system in Vibrio vulnificus. Nucleic Acids Res. 47, 10202–10211. 10.1093/nar/gkz746.

25. Wang, J.Y., Hoel, C.M., Al-Shayeb, B., Banfield, J.F., Brohawn, S.G., and Doudna, J.A. (2021). Structural coordination between active sites of a CRISPR reverse transcriptase-integrase complex. Nat Commun. 12, 2571. 10.1038/s41467-021-22900-y.

26. Yosef, I., Goren, M.G., and Qimron, U. (2012). Proteins and DNA elements essential for the CRISPR adaptation process in Escherichia coli. Nucleic Acids Res. 40, 5569–5576. 10.1093/nar/gks216.

27. Nuñez, J.K., Lee, A.S.Y., Engelman, A., and Doudna, J.A. (2015). Integrase-mediated spacer acquisition during CRISPR-Cas adaptive immunity. Nature 519, 193–198. 10.1038/nature14237.

28. Artamonova, D., Karneyeva, K., Medvedeva, S., Klimuk, E., Kolesnik, M., Yasinskaya, A., Samolygo, A., and Severinov, K. (2020). Spacer acquisition by Type III CRISPR–Cas system during bacteriophage infection of Thermus thermophilus. Nucleic Acids Res. 48, gkaa685-. 10.1093/nar/gkaa685.

29. Makarova, K.S., Wolf, Y.I., Iranzo, J., Shmakov, S.A., Alkhnbashi, O.S., Brouns, S.J.J., Charpentier, E., Cheng, D., Haft, D.H., Horvath, P., et al. (2020). Evolutionary classification of CRISPR–Cas systems: a burst of class 2 and derived variants. Nat. Rev. Microbiol. 18, 67–83. 10.1038/s41579-019-0299-x.

30. Flusche, T., and Rajan, R. (2022). Molecular Details of DNA Integration by CRISPR-Associated Proteins During Adaptation in Bacteria and Archaea. In Protein Reviews, Vol. 23 Advances in Experimental Medicine and Biology, Vol. 1414., M. Z. Atassi, ed., pp. 27–43. 10.1007/5584_2022_730.

31. Nuñez, J.K., Kranzusch, P.J., Noeske, J., Wright, A.V., Davies, C.W., and Doudna, J.A. (2014). Cas1-Cas2 complex formation mediates spacer acquisition during CRISPR-Cas adaptive immunity. Nat. Struct. Mol. Biol. 21, 528–534. 10.1038/nsmb.2820.

32. Rollins, M.F., Chowdhury, S., Carter, J., Golden, S.M., Wilkinson, R.A., Bondy-Denomy, J., Lander, G.C., and Wiedenheft, B. (2017). Cas1 and the Csy complex are opposing regulators of Cas2/3 nuclease activity. Proc. Natl. Acad. Sci. USA 23, 201616395. 10.1073/pnas.1616395114.

33. Wright, A.V., Liu, J.-J., Knott, G.J., Doxzen, K.W., Nogales, E., and Doudna, J.A. (2017). Structures of the CRISPR genome integration complex. Science 357, 1113–1118. 10.1126/science.aao0679.

34. Xiao, Y., Ng, S., Nam, K.H., and Ke, A. (2017). How type II CRISPR-Cas establish immunity through Cas1-Cas2-mediated spacer integration. Nature 550, 137–141. 10.1038/nature24020.

35. Lee, H., Dhingra, Y., and Sashital, D.G. (2019). The Cas4-Cas1-Cas2 complex mediates precise prespacer processing during CRISPR adaptation. eLife 8, 2460. 10.7554/elife.44248.

36. Tang, D., Li, H., Wu, C., Jia, T., He, H., Yao, S., Yu, Y., and Chen, Q. (2021). A distinct structure of Cas1–Cas2 complex provides insights into the mechanism for the longer spacer acquisition in Pyrococcus furiosus. Int. J. Biol. Macromol. 183, 379–386. 10.1016/j.ijbiomac.2021.04.074.

37. Wang, J.Y., Tuck, O.T., Skopintsev, P., Soczek, K.M., Li, G., Al-Shayeb, B., Zhou, J., and Doudna, J.A. (2023). Genome expansion by a CRISPR trimmer-integrase. Nature 618, 855–861. 10.1038/s41586-023-06178-2.

38. Sasnauskas, G., and Siksnys, V. (2020). CRISPR adaptation from a structural perspective. Curr. Opin. Struct. Biol. 65, 17–25. 10.1016/j.sbi.2020.05.015.

39. Budhathoki, J.B., Xiao, Y., Schuler, G., Hu, C., Cheng, A., Ding, F., and Ke, A. (2020). Real-time observation of CRISPR spacer acquisition by Cas1-Cas2 integrase. Nat. Struct. Mol. Biol. 27, 489–499. 10.1038/s41594-020-0415-7.

40. Stamos, J.L., Lentzsch, A.M., and Lambowitz, A.M. (2017). Structure of a Thermostable Group II Intron Reverse Transcriptase with Template-Primer and Its Functional and Evolutionary Implications. Mol. Cell 68, 926–939.e4. 10.1016/j.molcel.2017.10.024.

41. Zabrady, M., Zabrady, K., Li, A.W.H., and Doherty, A.J. (2023). Reverse transcriptases prime DNA synthesis. Nucleic Acids Res. 10.1093/nar/gkad478.

42. Aviram, N., Thornal, A.N., Zeevi, D., and Marraffini, L.A. (2022). Different modes of spacer acquisition by the Staphylococcus epidermidis type III-A CRISPR-Cas system. Nucleic Acids Res. 50, 1661–1672. 10.1093/nar/gkab1299.

43. Zhang, X., Garrett, S., Graveley, B.R., and Terns, M.P. (2021). Unique properties of spacer acquisition by the type III-A CRISPR-Cas system. Nucleic Acids Res. 50, 1562–1582. 10.1093/nar/gkab1193.

44. Frank, E.G., and Woodgate, R. (2007). Increased Catalytic Activity and Altered Fidelity of Human DNA Polymerase ι in the Presence of Manganese. J. Biol. Chem. 282, 24689–24696. 10.1074/jbc.m702159200.

45. Tabor, S., and Richardson, C.C. (1989). Effect of manganese ions on the incorporation of dideoxynucleotides by bacteriophage T7 DNA polymerase and Escherichia coli DNA polymerase I. Proc. Natl. Acad. Sci. USA 86, 4076–4080. 10.1073/pnas.86.11.4076.

46. Kent, T., Mateos-Gomez, P.A., Sfeir, A., and Pomerantz, R.T. (2016). Polymerase θ is a robust terminal transferase that oscillates between three different mechanisms during end-joining. elife *5*, e13740. 10.7554/elife.13740.

47. Motea, E.A., and Berdis, A.J. (2010). Terminal deoxynucleotidyl transferase: The story of a misguided DNA polymerase. Biochim. Biophys. Acta 1804, 1151–1166. 10.1016/j.bbapap.2009.06.030.

48. Lentzsch, A.M., Yao, J., Russell, R., and Lambowitz, A.M. (2019). Template-switching mechanism of a group II intron-encoded reverse transcriptase and its implications for biological function and RNA-Seq. J. Biol. Chem. 294, 19764–19784. 10.1074/jbc.ra119.011337.

49. Wang, H., and Lambowitz, A.M. (1993). The Mauriceville plasmid reverse transcriptase can initiate cDNA synthesis de novo and may be related to reverse transcriptase and DNA polymerase progenitor. Cell 75, 1071–1081.

50. Salas, M., Martín, G., Bernad, A., Garmendia, C., Lázaro, J.M., Zaballos, A., Serrano, M., Otero, M.J., Gutiérrez, J., and Parés, E. (1988). Protein-primed replication of bacteriophage phi 29 DNA. Biochim. Biophys. Acta. 951, 419–424. 10.1016/0167-4781(88)90115-7.

51. Wang, G.-H., and Seeger, C. (1992). The reverse transcriptase of hepatitis B virus acts as a protein primer for viral DNA synthesis. Cell 71, 663–670. 10.1016/0092-8674(92)90599-8.

52. Yushenova, I.A., and Arkhipova, I.R. (2018). Biochemical properties of bacterial reverse transcriptase-related (rvt) gene products: multimerization, protein priming, and nucleotide preference. Curr. Genetics 64, 1287–1301. 10.1007/s00294-018-0844-6.

53. Pan, B., Mitra, S.N., and Sundaralingam, M. (1999). Crystal Structure of an RNA 16-mer Duplex R(GCAGAGUUAAAUCUGC)2 with Nonadjacent G(Syn)·A+(Anti) Mispairs †. Biochemistry 38, 2826–2831. 10.1021/bi982122y.

54. Nuñez, J.K., Harrington, L.B., Kranzusch, P.J., Engelman, A.N., and Doudna, J.A. (2015). Foreign DNA capture during CRISPR-Cas adaptive immunity. Nature 527, 535–538. 10.1038/nature15760.

55. Simmon, V.F., and Lederberg, S. (1972). Degradation of Bacteriophage Lambda Deoxyribonucleic Acid After Restriction by Escherichia coli K-12. J. Bacteriol. 112, 161–169. 10.1128/jb.112.1.161-169.1972.

56. Dharmalingam, K., and Goldberg, E.B. (1976). Mechanism localisation and control of restriction cleavage of phage T4 and lambda chromosomes in vivo. Nature 260, 406–410. 10.1038/260406a0.

57. Cheng, K., Wilkinson, M., Chaban, Y., and Wigley, D.B. (2020). A conformational switch in response to Chi converts RecBCD from phage destruction to DNA repair. Nat. Struct. Mol. Biol. 27, 71–77. 10.1038/s41594-019-0355-2.

58. Ma, C.-H., Javanmardi, K., Finkelstein, I.J., and Jayaram, M. (2021). Disintegration promotes protospacer integration by the Cas1-Cas2 complex. elife *10*, e65763. 10.7554/elife.65763.

59. Blaszczyk, J., Tropea, J.E., Bubunenko, M., Routzahn, K.M., Waugh, D.S., Court, D.L., and Ji, X. (2001). Crystallographic and Modeling Studies of RNase III Suggest a Mechanism for Double-Stranded RNA Cleavage. Structure 9, 1225–1236. 10.1016/s0969-2126(01)00685-2.

60. Melek, M., Greene, E.C., and Shippen, D.E. (1996). Processing of Nontelomeric 3′ Ends by Telomerase: Default Template Alignment and Endonucleolytic Cleavage. Mol. Cell. Biol. 16, 3437– 3445. 10.1128/mcb.16.7.3437.

61. Dillingham, M.S., and Kowalczykowski, S.C. (2008). RecBCD Enzyme and the Repair of Double-Stranded DNA Breaks. Microbiol. Mol. Biol. Rev. 72, 642–671. 10.1128/mmbr.00020-08.

62. Goldmark, P.J., and Linn, S. (1972). Purification and Properties of the recBC DNase of Escherichia coli K-12. J. Biol. Chem. 247, 1849–1860. 10.1016/s0021-9258(19)45550-6.

63. Zhao, C., and Pyle, A.M. (2016). Crystal structures of a group II intron maturase reveal a missing link in spliceosome evolution. Nat. Struct. Mol. Biol. 23, 558–565. 10.1038/nsmb.3224.

64. Toro, N., Martinez-Abarca, F., Mestre, M.R., and González-Delgado, A. (2019). Multiple origins of reverse transcriptases linked to CRISPR-Cas systems. RNA Biol. 7, 1–8. 10.1080/15476286.2019.1639310.

65. Zabrady, K., Zabrady, M., Kolesar, P., Li, A.W.H., and Doherty, A.J. (2021). CRISPR-Associated Primase-Polymerases are implicated in prokaryotic CRISPR-Cas adaptation. Nat. Commun. 12, 3690. 10.1038/s41467-021-23535-9.

66. Guo, H., Arambula, D., Ghosh, P., and Miller, J.F. (2014). Diversity-generating Retroelements in Phage and Bacterial Genomes. Microbiology Spectrum 2, 1237–1252. 10.1128/microbiolspec.mdna3-0029-2014.

67. Handa, S., Jiang, Y., Tao, S., Foreman, R., Schinazi, R.F., Miller, J.F., and Ghosh, P. (2018). Template-assisted synthesis of adenine-mutagenized cDNA by a retroelement protein complex. Nucleic Acids Res. 46, 9711–9725. 10.1093/nar/gky620.

68. Lambowitz, A.M., and Belfort, M. (2015). Mobile Bacterial Group II Introns at the Crux of Eukaryotic Evolution. Microbiology Spectrum 3, MDNA3-0050–2014. 10.1128/microbiolspec.mdna3-0050-2014.

69. Anzalone, A.V., Randolph, P.B., Davis, J.R., Sousa, A.A., Koblan, L.W., Levy, J.M., Chen, P.J., Wilson, C., Newby, G.A., Raguram, A., et al. (2019). Search-and-replace genome editing without double-strand breaks or donor DNA. Nature 576, 149–157. 10.1038/s41586-019-1711-4.

70. Liu, B., Dong, X., Cheng, H., Zheng, C., Chen, Z., Rodríguez, T.C., Liang, S.-Q., Xue, W., and Sontheimer, E.J. (2022). A split prime editor with untethered reverse transcriptase and circular RNA template. Nat. Biotechnol. 40, 1388–1393. 10.1038/s41587-022-01255-9.

71. Chen, P.J., and Liu, D.R. (2023). Prime editing for precise and highly versatile genome manipulation. Nat. Rev. Genet. 24, 161–177. 10.1038/s41576-022-00541-1.

72. Grünewald, J., Miller, B.R., Szalay, R.N., Cabeceiras, P.K., Woodilla, C.J., Holtz, E.J.B., Petri, K., and Joung, J.K. (2023). Engineered CRISPR prime editors with compact, untethered reverse transcriptases. Nat. Biotechnol. 41, 337–343. 10.1038/s41587-022-01473-1.

73. Mohr, S., Ghanem, E., Smith, W., Sheeter, D., Qin, Y., King, O., Polioudakis, D., Iyer, V.R., Hunicke-Smith, S., Swamy, S., et al. (2013). Thermostable group II intron reverse transcriptase fusion proteins and their use in cDNA synthesis and next-generation RNA sequencing. RNA 19, 958–970. 10.1261/rna.039743.113.

74. Studier, F.W. (2005). Protein production by auto-induction in high density shaking cultures. Protein Expr. Purif. 41, 207–234. 10.1016/j.pep.2005.01.016.

75. Xu, H., Nottingham, R., and Lambowitz, A.M. (2021). TGIRT-seq Protocol for the Comprehensive Profiling of Coding and Non-coding RNA Biotypes in Cellular, Extracellular Vesicle, and Plasma RNAs. Bio-protocol 11, e4239. 10.21769/bioprotoc.4239.

76. Xu, H., Yao, J., Wu, D.C., and Lambowitz, A.M. (2019). Improved TGIRT-seq methods for comprehensive transcriptome profiling with decreased adapter dimer formation and bias correction. Sci. Rep. 9, 7953. 10.1038/s41598-019-44457-z.

77. Katoh, K., Misawa, K., Kuma, K., and Miyata, T. (2002). MAFFT: a novel method for rapid multiple sequence alignment based on fast Fourier transform. Nucleic Acids Res. 30, 3059–3066. 10.1093/nar/gkf436.

