## Supplementary figures and images for "Mechanisms used for cDNA synthesis and site-specific integration of RNA into DNA genomes by a reverse transcriptase-Cas1 fusion protein"

### Supplemental Figures S1 to S7

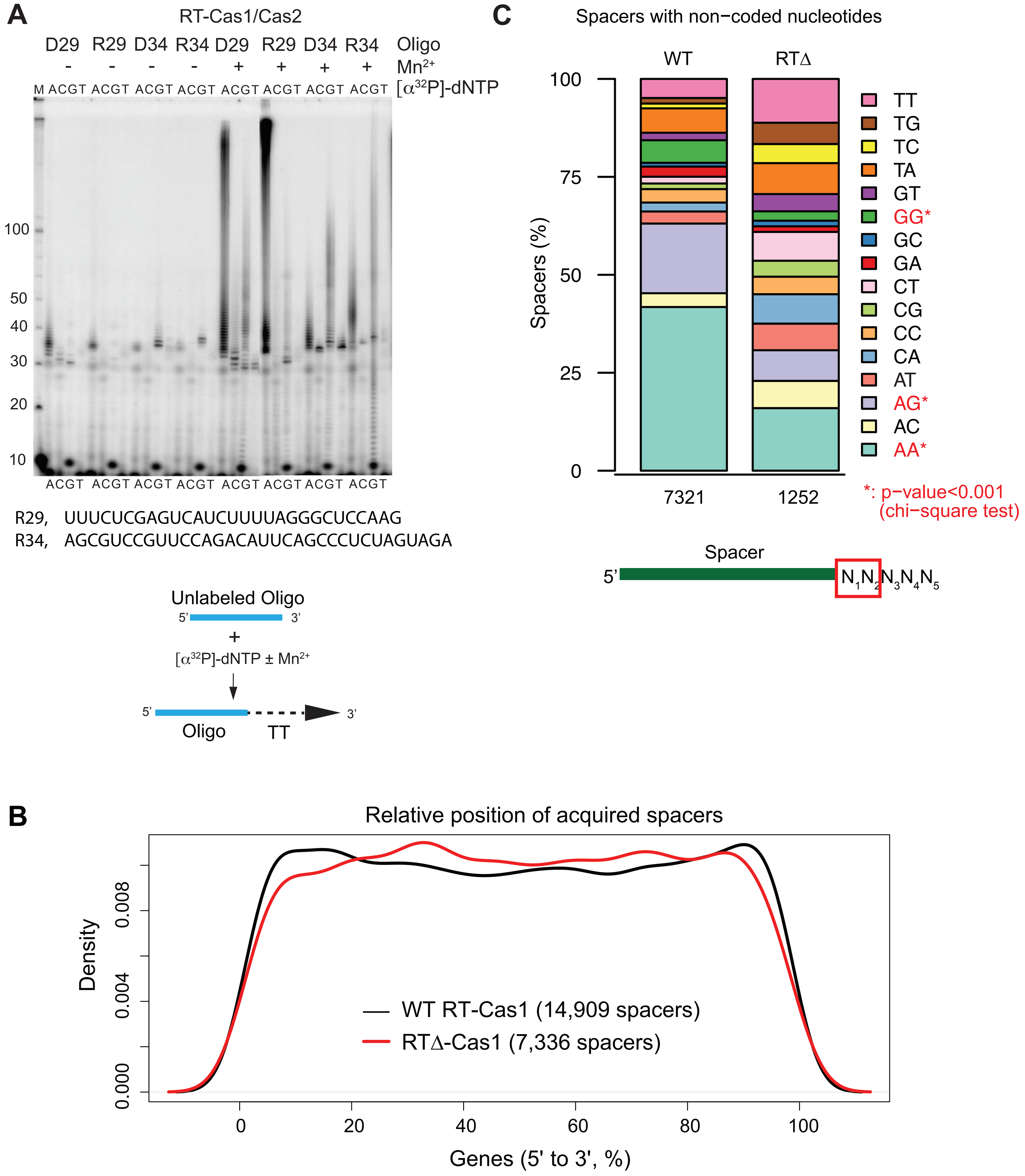

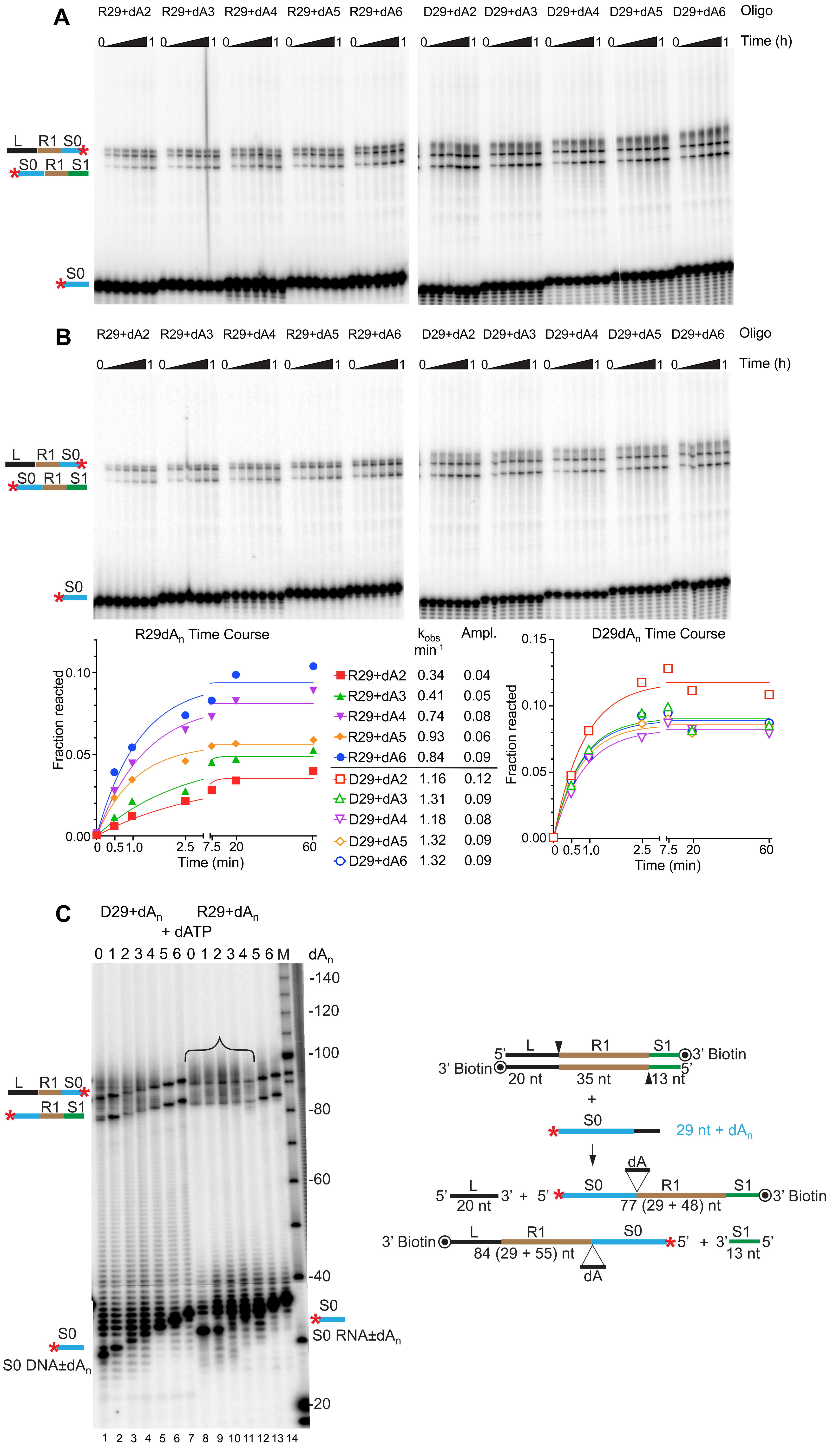

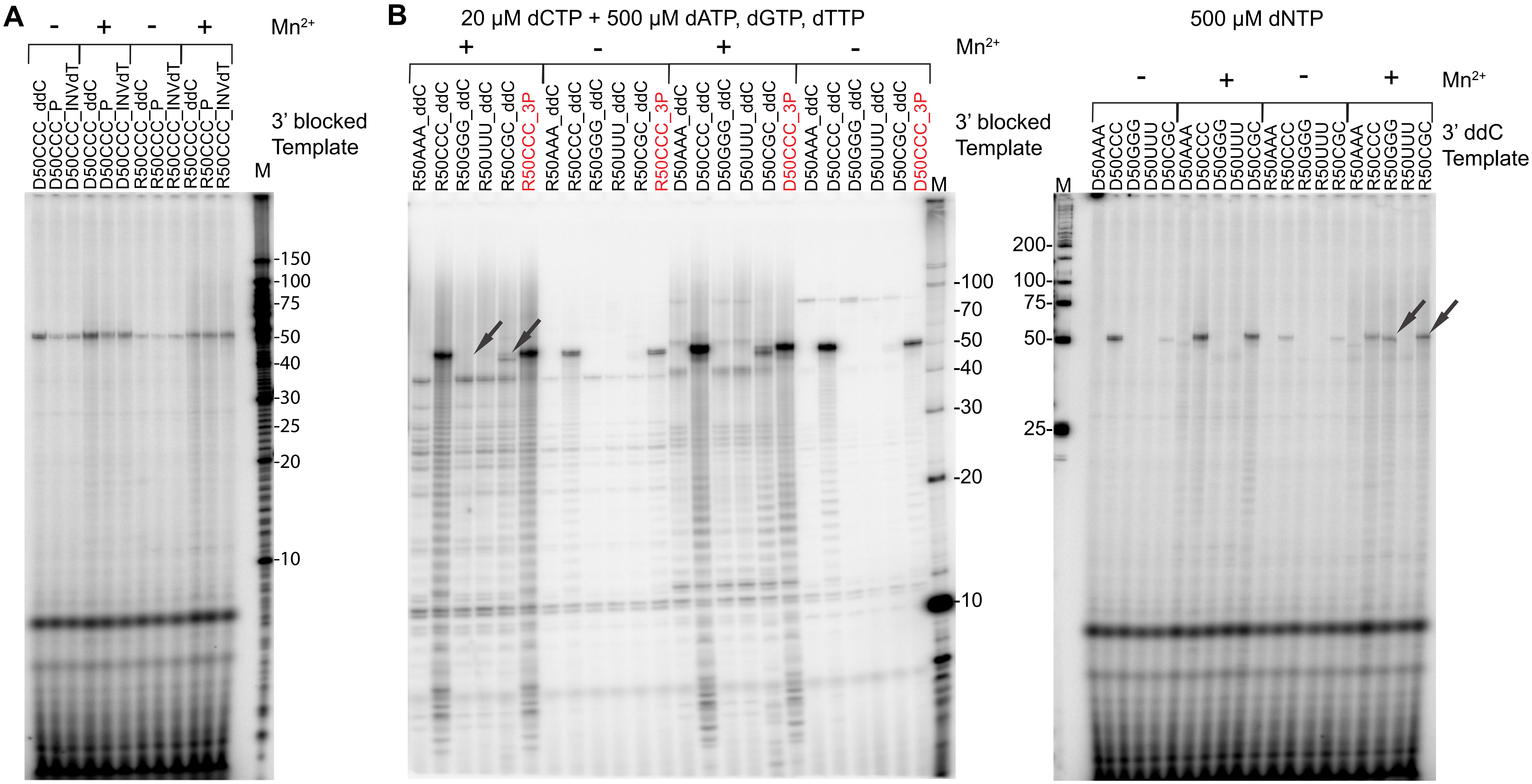

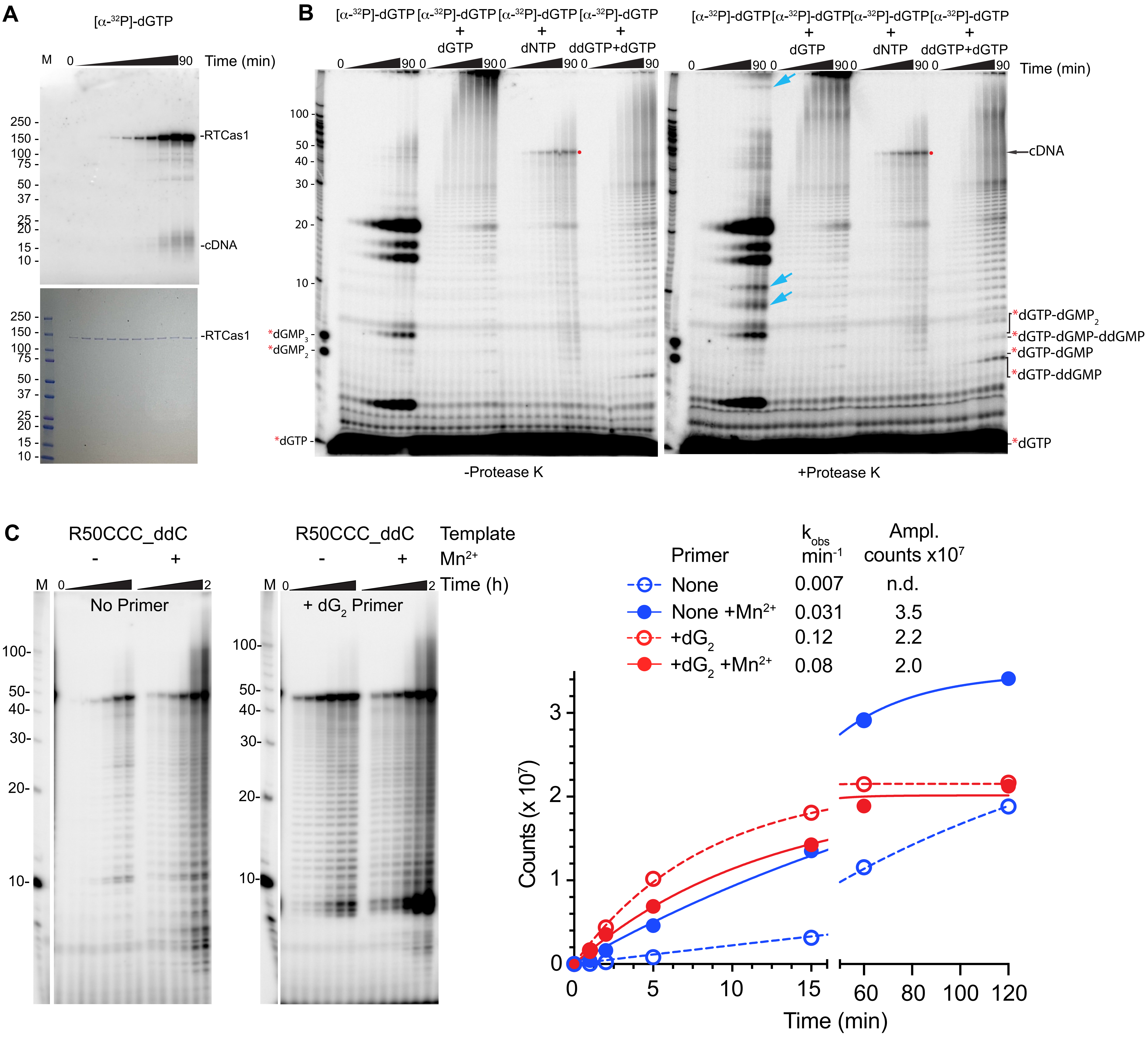

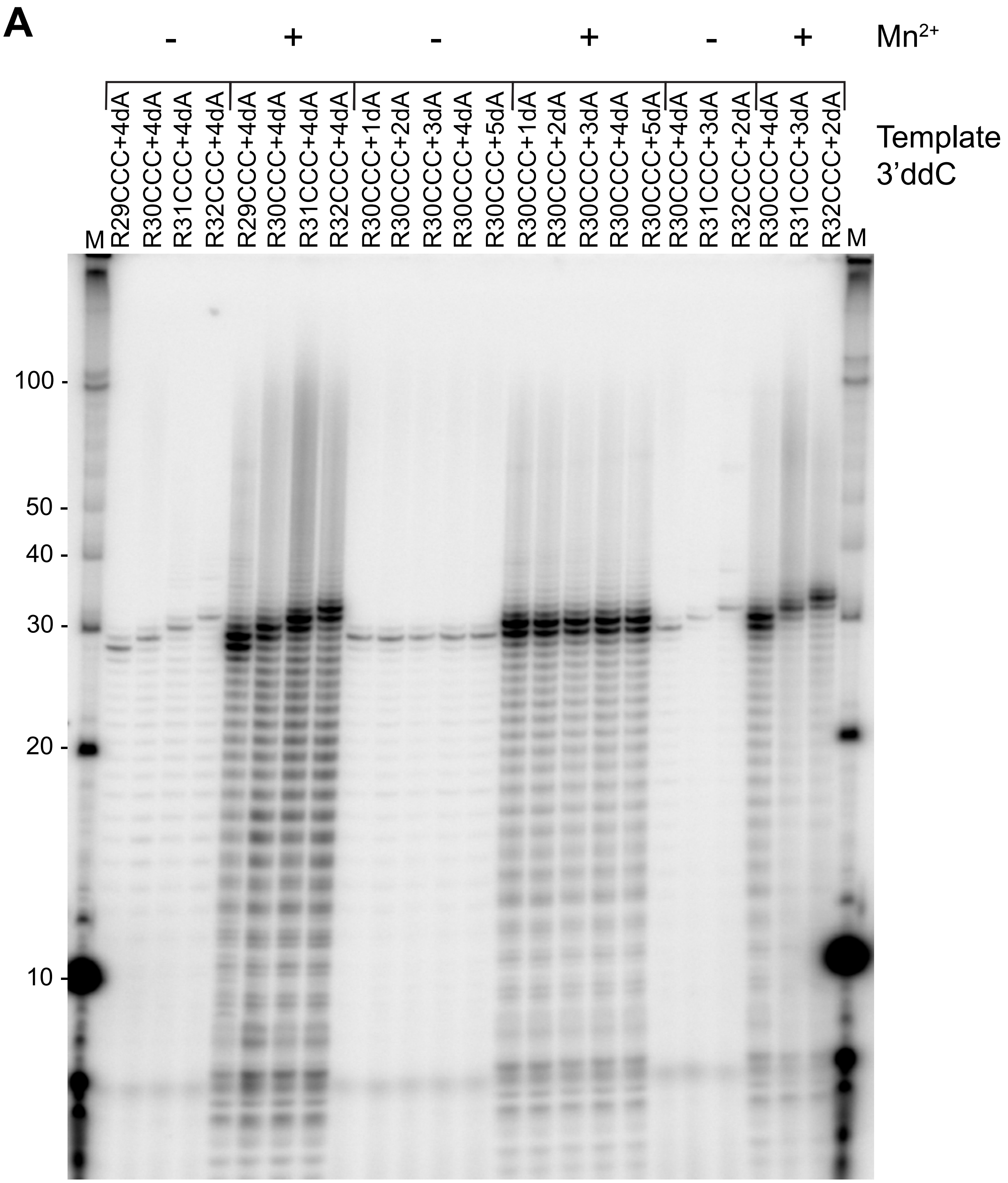

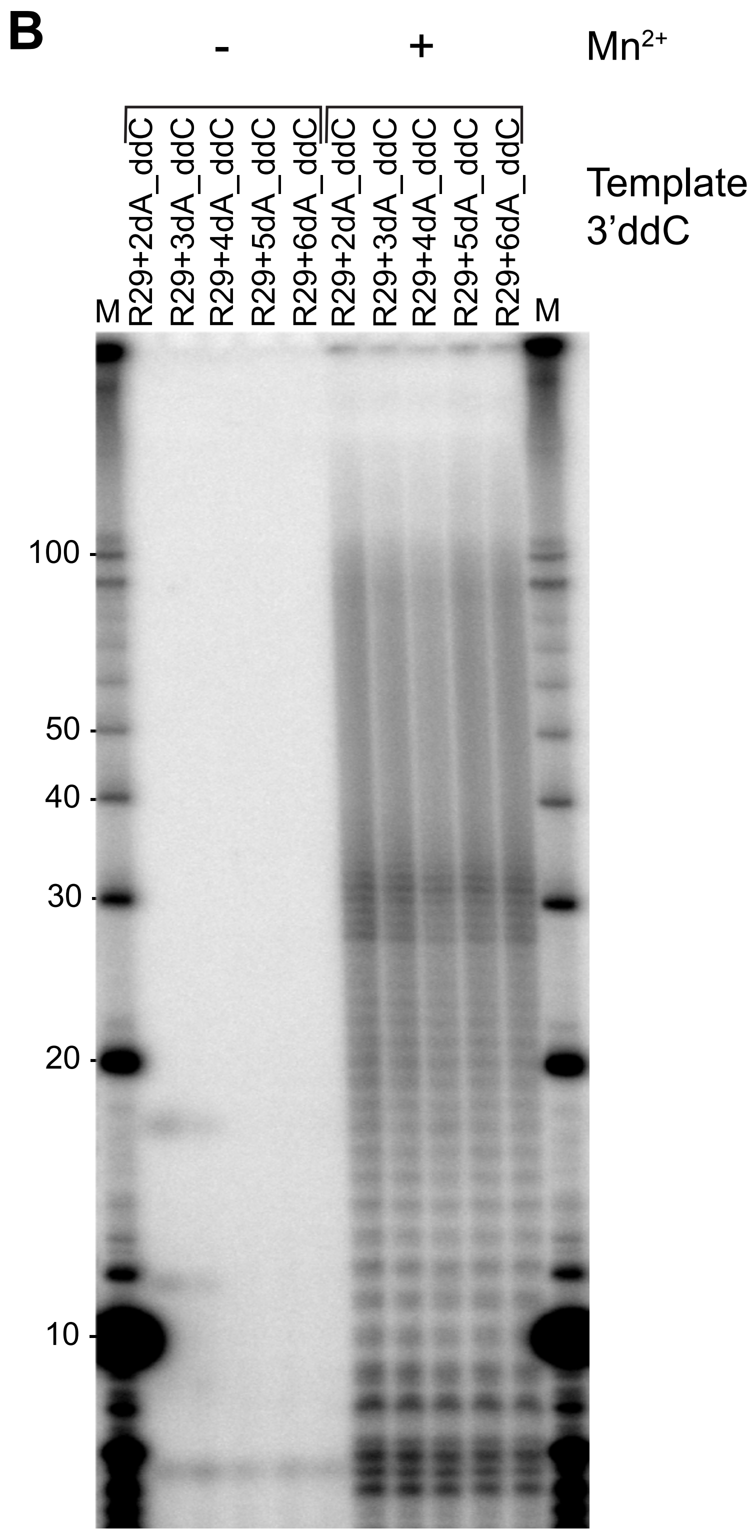

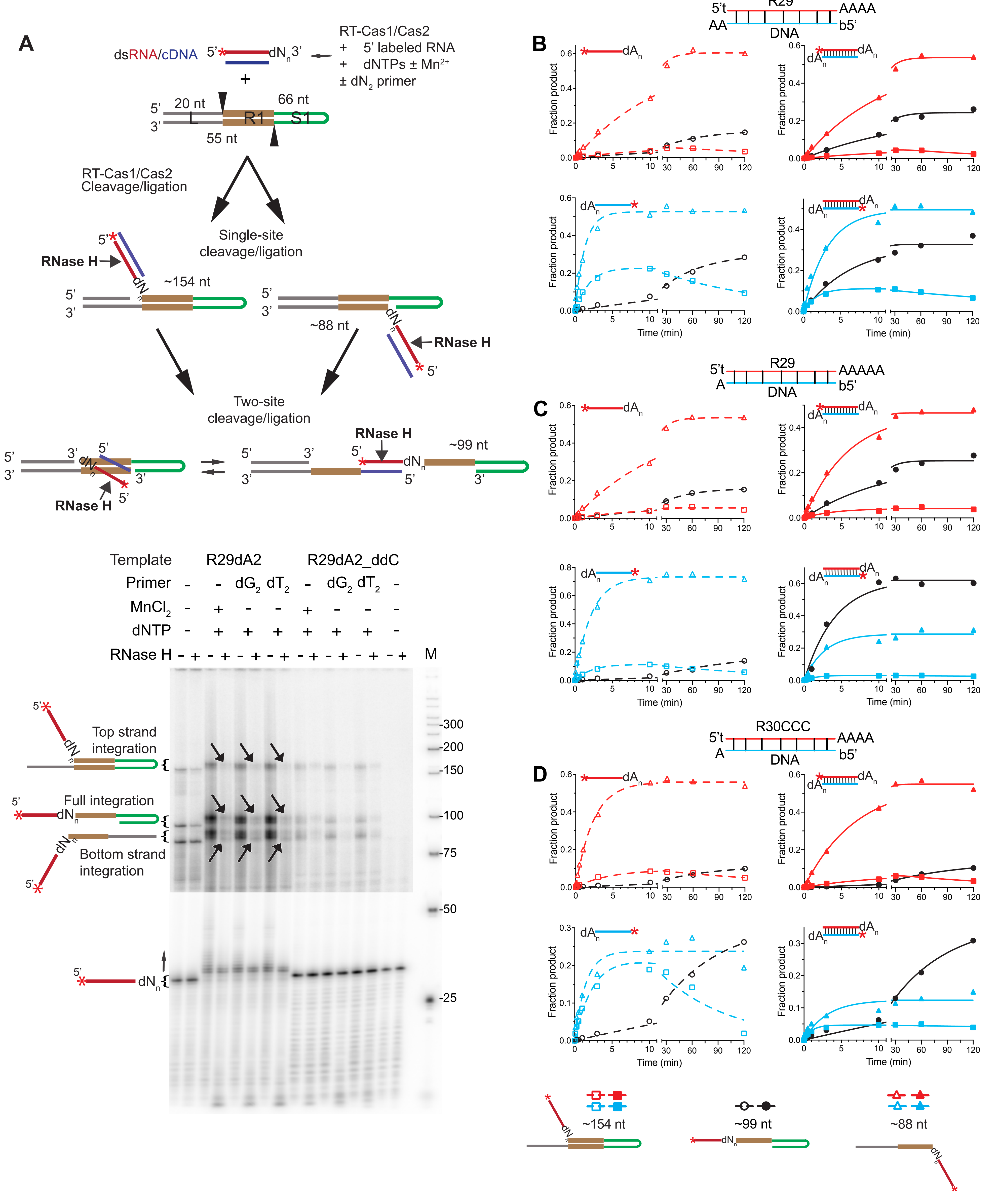

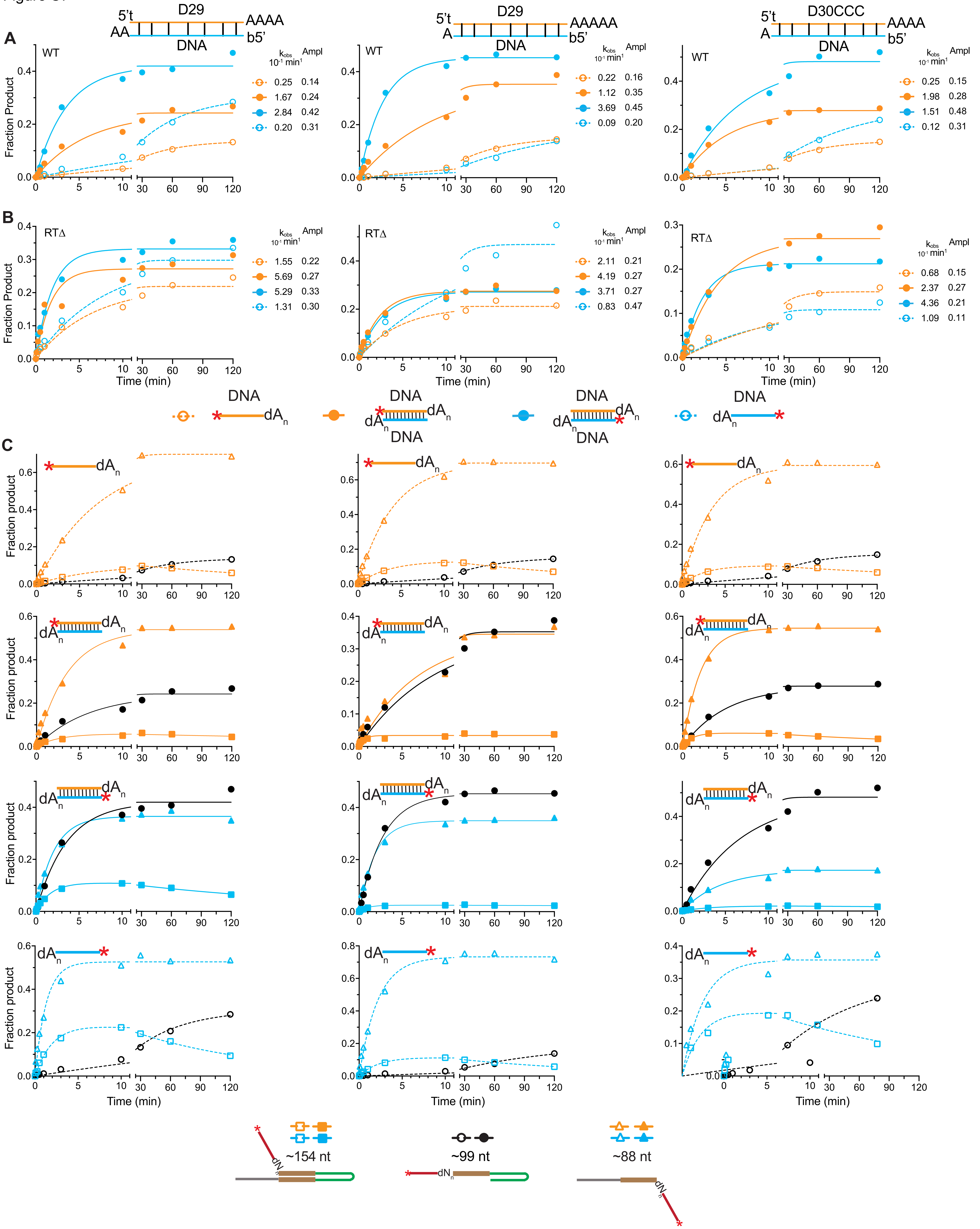
