## Supplemental data file for "Mechanisms used for cDNA synthesis and site-specific integration of RNA into DNA genomes by a reverse transcriptase-Cas1 fusion protein"

This file contains:

1. List of oligonucleotides used. RNA sequences are shown in bold and DNA sequences in regular font.
2. A list of $R^2$ values for curve fits for experiments shown in Figures 1F, 2C, S2B, S4C, 6B-D, S6A-C and S7A-C.
3. Gels for spacer acquisition time courses. (A) Gels for Figure 6B and S6B. (B) Gels for Figure 6C and S6C. (C) Gels for Figure 6D and S6D. (D) Gels for Figure S7A and S7C. (E) Gels for Figure S7B and S7C.
4. Table S1: Datasets used.
5. Data for Figures 1B and 1C.

Oligonucleotides

| Project | Name | Sequence |
|---|---|---|
| **cDNA templates** | R50CCC | **GCAAUAAUCUAUACAAUACAACACAUACAAACAAAUUCUUAAGGUCCCAA** |
| | D50CCC | GCAATAATCTATACAATACAACACATACAAACAAATTCTTAAGGTCCCAA |
| | R50CCC_ddC | **GCAAUAAUCUAUACAAUACAACACAUACAAACAAAUUCUUAAGGUCCCAA**/3ddC |
| | D50CCC_ddC | GCAATAATCTATACAATACAACACATACAAACAAATTCTTAAGGTCCCAA/3ddC |
| | R50CCC_3P | **GCAAUAAUCUAUACAAUACAACACAUACAAACAAAUUCUUAAGGUCCCAA**/3Phos |
| | D50CCC_3P | GCAATAATCTATACAATACAACACATACAAACAAATTCTTAAGGTCCCAA/3Phos |
| | R50CCC_INVdT | **GCAAUAAUCUAUACAAUACAACACAUACAAACAAAUUCUUAAGGUCCCAA**/3INVdT |
| | D50CCC_INVdT | GCAATAATCTATACAATACAACACATACAAACAAATTCTTAAGGTCCCAA/3INVdT |
| | R50CGC_ddC | **GCAAUAAUCUAUACAAUACAACACAUACAAACAAAUUCUUAAGGUCGCAA**/3ddC |
| | D50CGC_ddC | GCAATAATCTATACAATACAACACATACAAACAAATTCTTAAGGTCGCAA/3ddC |
| | R50AAA_ddC | **GCAAUAAUCUAUACAAUACAACACAUACAAACAAAUUCUUAAGGUAAAAA**/3ddC |
| | D50AAA_ddC | GCAATAATCTATACAATACAACACATACAAACAAATTCTTAAGGTAAAAA/3ddC |
| | R50GGG_ddC | **GCAAUAAUCUAUACAAUACAACACAUACAAACAAAUUCUUAAGGUGGGAA**/3ddC |
| | D50GGG_ddC | GCAATAATCTATACAATACAACACATACAAACAAATTCTTAAGGTGGGAA/3ddC |
| | R50UUU_ddC | **GCAAUAAUCUAUACAAUACAACACAUACAAACAAAUUCUUAAGGUUUUAA**/3ddC |
| | D50TTT_ddC | GCAATAATCTATACAATACAACACATACAAACAAATTCTTAAGGTTTTAA/3ddC |
| | R29CCC+dA4_ddC | **CACAUACAAACAAAUUCUUAAGGUCCCAA**AAAA/3ddC |
| | R30CCC+dA4_ddC | **ACACAUACAAACAAAUUCUUAAGGUCCCAA**AAAA/3ddC |
| | R31CCC+dA4_ddC | **AACACAUACAAACAAAUUCUUAAGGUCCCAA**AAAA/3ddC |
| | R32CCC+dA4_ddC | **CAACACAUACAAACAAAUUCUUAAGGUCCCAA**AAAA/3ddC |
| | R31CCC+dA3_ddC | **AACACAUACAAACAAAUUCUUAAGGUCCCAA**AAA/3ddC |
| | R32CCC+dA4_ddC | **CAACACAUACAAACAAAUUCUUAAGGUCCCAA**AA/3ddC |
| | R30CCC+dA1ddC | **ACACAUACAAACAAAUUCUUAAGGUCCCAA**A/3ddC |
| | R30CCC+dA2ddC | **ACACAUACAAACAAAUUCUUAAGGUCCCAA**AA3ddC |
| | R30CCC+dA3ddC | **ACACAUACAAACAAAUUCUUAAGGUCCCAA**AAA/3ddC |
| | R30CCC+dA4ddC | **ACACAUACAAACAAAUUCUUAAGGUCCCAA**AAAA/3ddC |
| | R30CCC+dA5ddC | **ACACAUACAAACAAAUUCUUAAGGUCCCAA**AAAAA/3ddC |
| | R29+dA2_ddC | **UUUCUCGAGUCAUCUUUUAGGGCUCCAAG**AA/3ddC |
| | R29+dA3_ddC | **UUUCUCGAGUCAUCUUUUAGGGCUCCAAG**AAA/3ddC |
| | R29+dA4_ddC | **UUUCUCGAGUCAUCUUUUAGGGCUCCAAG**AAAA/3ddC |
| | R29+dA5_ddC | **UUUCUCGAGUCAUCUUUUAGGGCUCCAAG**AAAAA/3ddC |
| | R29+dA6_ddC | **UUUCUCGAGUCAUCUUUUAGGGCUCCAAG**AAAAAA/3ddC |
| **Cloning** | Cas2_pet_5 | GATCCATATGATGAGAATATACCTTGCCTGTTT |
| | Cas2_pet_3 | TTTGGATCCTTATCATAACAGGACGGCGG |
| **Primers** | $dA_2$ | AA |
| | $dC_2$ | CC |
| | $dG_2$ | GG |
| | $dT_2$ | TT |
| **CRISPR DNA** | MMB1Lead40-5 | AGGTTAACCTGCTGAAATGATTGG |
| | MMB1crisp3_r1 | GGAGATCTTTAAAGTCTCAACG |
| | L20R1top_3BIO | TTGGAAAAAATAAGGGTACTGTTTCAGACCCGCTGGCCGCTTAGGCCGTTGAGACTTTAAAGATCTCC/3BIOTIN |
| | L20R1bot_3Bio | GGAGATCTTTAAAGTCTCAACGGCCTAAGCGGCCAGCGGGTCTGAAACAGTACCCTTATTTTTTCCAA/3BIOTIN |
| | CRISPR-HP3blunt | TTGGAAAAAATAAGGGTACTGTTTCAGACCCGCTGGCCGCTTAGGCCGTTGAGACTTTAAAGATCTCCTACATGGAGATCTTTAAAGTCTCAACGGCCTAAGCGGCCAGCGGGTCTGAAACAGTACCCTTATTTTTTCCAA |
| **CRISPR assay** | R29 | **UUUCUCGAGUCAUCUUUUAGGGCUCCAAG** |
| | R29+dA1 | **UUUCUCGAGUCAUCUUUUAGGGCUCCAAG**A |
| | R29+dA2 | **UUUCUCGAGUCAUCUUUUAGGGCUCCAAG**AA |
| | R29+dA3 | **UUUCUCGAGUCAUCUUUUAGGGCUCCAAG**AAA |
| | R29+dA4 | **UUUCUCGAGUCAUCUUUUAGGGCUCCAAG**AAAA |
| | R29+dA5 | **UUUCUCGAGUCAUCUUUUAGGGCUCCAAG**AAAAA |
| | R29+dA6 | **UUUCUCGAGUCAUCUUUUAGGGCUCCAAG**AAAAAA |
| | D29 | TTTCTCGAGTCATCTTTTAGGGCTCCAAG |
| | D29+dA1 | TTTCTCGAGTCATCTTTTAGGGCTCCAAGA |
| | D29+dA2 | TTTCTCGAGTCATCTTTTAGGGCTCCAAGAA |
| | D29+dA3 | TTTCTCGAGTCATCTTTTAGGGCTCCAAGAAA |
| | D29+dA4 | TTTCTCGAGTCATCTTTTAGGGCTCCAAGAAAA |
| | D29+dA5 | TTTCTCGAGTCATCTTTTAGGGCTCCAAGAAAAA |
| | D29+dA6 | TTTCTCGAGTCATCTTTTAGGGCTCCAAGAAAAAA |
| | D29r+dA1 | CTTGGAGCCCTAAAAGATGACTCGAGAAAA |
| | D29r+dA2 | CTTGGAGCCCTAAAAGATGACTCGAGAAAAA |
| | R30CCC+dA4 | **ACACAUACAAACAAAUUCUUAAGGUCCCAA**AAAA |
| | D30CCCr+dA1 | TTGGGACCTTAAGAATTTGTTTGTATGTGTA |
| **Terminal transferase** | R29 | **UUUCUCGAGUCAUCUUUUAGGGCUCCAAG** |
| | D29 | TTTCTCGAGTCATCTTTTAGGGCTCCAAG |
| | R34 | **AGCGUCCGUUCCAGACAUUCAGCCCUCUAGUAGA** |
| | D34 | AGCGTCCGTTCCAGACATTCAGCCCTCTAGTAGA |
| **Oligonucleotide markers** | dG2 | GG |
| | dG3 | GGG |

R squared values for curve fits

| Protospacer | Band | R squared values | | |
|---|---|---|---|---|
| | | WT 5 nM protospacer | RTΔ 5 nM protospacer | WT 250 nM protospacer |
| *R29+dA4 ss | **154 nt** | 0.9932 | 0.9865 | 0.8303 |
| *R29+dA4 ss | **99 nt** | 0.9983 | 0.9901 | 0.9993 |
| *R29+dA4 ss | **88 nt** | 0.9968 | 0.97 | 0.9676 |
| *R29+dA4/D29r+dA2 ds | **154 nt** | 0.9882 | 0.9255 | 0.8719 |
| *R29+dA4/D29r+dA2 ds | **99 nt** | 0.9929 | 0.9827 | 0.9972 |
| *R29+dA4/D29r+dA2 ds | **88 nt** | 0.9974 | 0.9357 | 0.9693 |
| R29+dA4/*D29r+dA2 ds | **154 nt** | 0.9727 | 0.9534 | 0.4491 |
| R29+dA4/*D29r+dA2 ds | **99 nt** | 0.9775 | 0.9595 | 0.9943 |
| R29+dA4/*D29r+dA2 ds | **88 nt** | 0.926 | 0.8398 | 0.8862 |
| *D29r+dA2 ss | **154 nt** | 0.9952 | 0.9679 | 0.8245 |
| *D29r+dA2 ss | **99 nt** | 0.9915 | 0.9721 | 0.9908 |
| *D29r+dA2 ss | **88 nt** | 0.9863 | 0.9393 | 0.9476 |
| *D29+dA4 ss | **154 nt** | 0.9984 | 0.9759 | |
| *D29+dA4 ss | **99 nt** | 0.9993 | 0.9712 | |
| *D29+dA4 ss | **88 nt** | 0.9983 | 0.9408 | |
| *D29+dA4/D29r+dA2 ds | **154 nt** | 0.9485 | 0.9013 | |
| *D29+dA4/D29r+dA2 ds | **99 nt** | 0.9726 | 0.9213 | |
| *D29+dA4/D29r+dA2 ds | **88 nt** | 0.989 | 0.7121 | |
| D29+dA4/*D29r+dA2 ds | **154 nt** | 0.9924 | 0.9633 | |
| D29+dA4/*D29r+dA2 ds | **99 nt** | 0.9853 | 0.9754 | |
| D29+dA4/*D29r+dA2 ds | **88 nt** | 0.9896 | 0.7568 | |
| *R29+dA5 ss | **154 nt** | 0.9681 | 0.9884 | 0.9306 |
| *R29+dA5 ss | **99 nt** | 0.9994 | 0.9661 | 0.9972 |
| *R29+dA5 ss | **88 nt** | 0.9981 | 0.974 | 0.9323 |
| *R29+dA5/D29r+dA1 ds | **154 nt** | 0.9439 | 0.9705 | 0.567 |
| *R29+dA5/D29r+dA1 ds | **99 nt** | 0.9893 | 0.9807 | 0.9939 |
| *R29+dA5/D29r+dA1 ds | **88 nt** | 0.9954 | 0.9695 | 0.9628 |
| R29+dA5/*D29r+dA1 ds | **154 nt** | 0.9908 | 0.9879 | 0.9125 |
| R29+dA5/*D29r+dA1 ds | **99 nt** | 0.9862 | 0.9946 | 0.9981 |
| R29+dA5/*D29r+dA1 ds | **88 nt** | 0.9766 | 0.9635 | 0.9778 |
| *D29r+dA1 ss | **154 nt** | 0.9959 | 0.9641 | 0.7696 |
| *D29r+dA1 ss | **99 nt** | 0.9794 | 0.9549 | 0.9964 |
| *D29r+dA1 ss | **88 nt** | 0.9934 | 0.9329 | 0.9346 |
| *D29+dA5 ss | **154 nt** | 0.9978 | 0.9413 | |
| *D29+dA5 ss | **99 nt** | 0.9961 | 0.972 | |
| *D29+dA5 ss | **88 nt** | 0.9981 | 0.9169 | |
| *D29+dA5/D29r+dA1 ds | **154 nt** | 0.8823 | 0.937 | |
| *D29+dA5/D29r+dA1 ds | **99 nt** | 0.9801 | 0.9854 | |
| *D29+dA5/D29r+dA1 ds | **88 nt** | 0.957 | 0.9118 | |
| D29+dA5/*D29r+dA1 ds | **154 nt** | 0.9781 | 0.9513 | |
| D29+dA5/*D29r+dA1 ds | **99 nt** | 0.9968 | 0.99 | |
| D29+dA5/*D29r+dA1 ds | **88 nt** | 0.995 | 0.9569 | |
| *R30CCC+dA4 ss | **154 nt** | 0.9507 | 0.9585 | 0.7574 |
| *R30CCC+dA4 ss | **99 nt** | 0.9885 | 0.9649 | 0.9938 |
| *R30CCC+dA4 ss | **88 nt** | 0.9974 | 0.9406 | 0.8893 |
| *R30CCC+dA4/D30CCCr+dA1 ds | **154 nt** | 0.9974 | 0.9385 | 0.5661 |
| *R30CCC+dA4/D30CCCr+dA1 ds | **99 nt** | 0.999 | 0.9826 | 0.9856 |
| *R30CCC+dA4/D30CCCr+dA1 ds | **88 nt** | 0.9976 | 0.9686 | 0.928 |
| R30CCC+dA4/*D30CCCr+dA1 ds | **154 nt** | 0.9614 | 0.8651 | 0.8928 |
| R30CCC+dA4/*D30CCCr+dA1 ds | **99 nt** | 0.9949 | 0.9763 | 0.9407 |
| R30CCC+dA4/*D30CCCr+dA1 ds | **88 nt** | 0.8734 | 0.8389 | 0.8487 |
| *D30CCCr+dA1 ss | **154 nt** | 0.928 | 0.8795 | 0.5414 |
| *D30CCCr+dA1 ss | **99 nt** | 0.9976 | 0.941 | 0.9415 |
| *D30CCCr+dA1 ss | **88 nt** | 0.9328 | 0.8814 | 0.8747 |
| *D30CCC+dA4 ss | **154 nt** | 0.9948 | 0.9655 | |
| *D30CCC+dA4 ss | **99 nt** | 0.9944 | 0.9742 | |
| *D30CCC+dA4 ss | **88 nt** | 0.9917 | 0.9289 | |
| *D30CCC+dA4/D30CCCr+dA1 ds | **154 nt** | 0.9946 | 0.9598 | |
| *D30CCC+dA4/D30CCCr+dA1 ds | **99 nt** | 0.9966 | 0.982 | |
| *D30CCC+dA4/D30CCCr+dA1 ds | **88 nt** | 0.9991 | 0.9555 | |
| D30CCC+dA4/*D30CCCr+dA1 ds | **154 nt** | 0.9891 | 0.8706 | |
| D30CCC+dA4/*D30CCCr+dA1 ds | **99 nt** | 0.9828 | 0.9903 | |
| D30CCC+dA4/*D30CCCr+dA1 ds | **88 nt** | 0.9802 | 0.8806 | |
| *D30CCCr+dA1 ss | **154 nt** | 0.9732 | 0.9246 | |
| *D30CCCr+dA1 ss | **99 nt** | 0.9979 | 0.9557 | |
| *D30CCCr+dA1 ss | **88 nt** | 0.9688 | 0.9009 | |

| Protospacer | | R squared values | |
|---|---|---|---|
| | | Fig 2 | Fig. S2 |
| *R29+dA2 | | 0.9663 | 0.9652 |
| *R29+dA3 | | 0.9614 | 0.9720 |
| *R29+dA4 | | 0.9742 | 0.9743 |
| *R29+dA5 | | 0.9807 | 0.9851 |
| *R29+dA6 | | 0.960 | 0.9537 |
| *D29r+dA2 | | 0.9595 | 0.9769 |
| *D29r+dA3 | | 0.9886 | 0.9658 |
| *D29r+dA4 | | 0.9981 | 0.9904 |
| *D29r+dA5 | | 0.9004 | 0.9876 |
| *D29r+dA6 | | 0.9885 | 0.9761 |
| | **dNTP** | **Fig. 1F** | |
| *R29 | dA | 0.9501 | |
| *R29 | dA + $Mn^{2+}$ | 0.9990 | |
| *R29 | dC + $Mn^{2+}$ | 0.9033 | |
| *R29 | dG + $Mn^{2+}$ | 0.9858 | |
| *D29 | dA | 0.9953 | |
| *D29 | dA + $Mn^{2+}$ | 0.9954 | |
| *D29 | $dC^{+}$ | 0.9913 | |
| *D29 | dC + $Mn^{2+}$ | 0.9821 | |
| *D29 | dG | 0.9131 | |
| *D29 | dG + $Mn^{2+}$ | 0.9704 | |
| *D29 | dT | 0.8459 | |
| *D29 | dT + $Mn^{2+}$ | 0.9050 | |
| | **Primer** | **Fig. S4C** | |
| R50CCC_ddC | None | 0.9992 | |
| R50CCC_ddC | None + $Mn^{2+}$ | 0.9990 | |
| R50CCC_ddC | $dG_2$ | 0.9983 | |
| R50CCC_ddC | $dG_2$ + $Mn^{2+}$ | 0.9938 | |

Supplemental data file

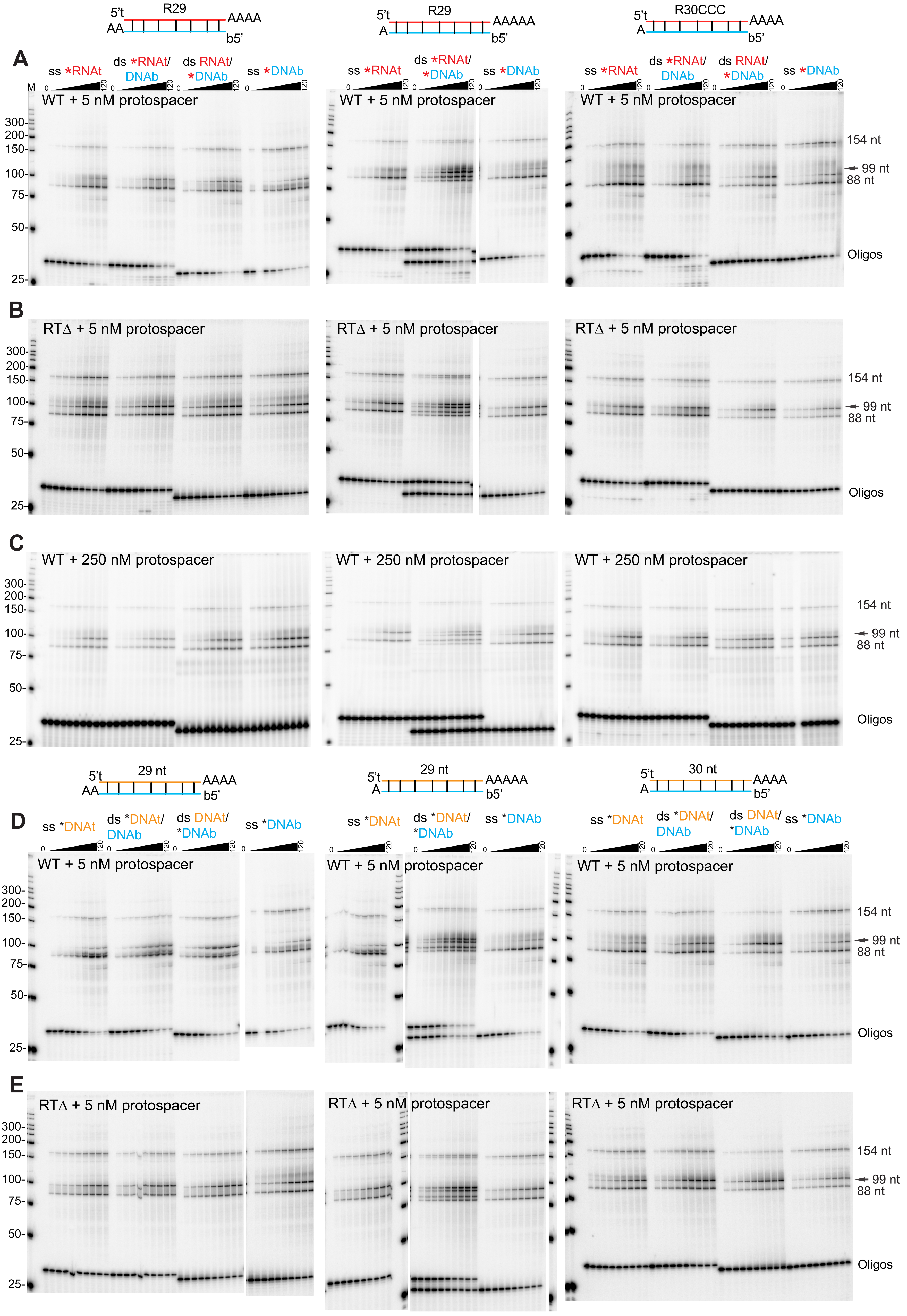

Table S1. Datasets used.

| Dataset | Template | Primer | $Mn^{2+}$ | RNaseA | Template oligo sequences | Accession |
|---|---|---|---|---|---|---|
| Dataset1 | R29dA3_ddC | - | - | - | UUUCUCGAGUCAUCUUUUAGGGCUCCAAGAAAAA/ddC | SRR25570276 |
| Dataset2 | R29dA4_ddC | - | - | - | UUUCUCGAGUCAUCUUUUAGGGCUCCAAGAAAAAA/ddC | SRR25570275 |
| Dataset3 | R29dA6_ddC | - | - | - | UUUCUCGAGUCAUCUUUUAGGGCUCCAAGAAAAAAAA/ddC | SRR25570264 |
| Dataset4 | R29dA3_ddC | - | + | - | UUUCUCGAGUCAUCUUUUAGGGCUCCAAGAAAAA/ddC | SRR25570253 |
| Dataset5 | R29dA4_ddC | - | + | - | UUUCUCGAGUCAUCUUUUAGGGCUCCAAGAAAAAA/ddC | SRR25570245 |
| Dataset6 | R29dA6_ddC | - | + | - | UUUCUCGAGUCAUCUUUUAGGGCUCCAAGAAAAAAAA/ddC | SRR25570244 |
| Dataset7 | R50AAA_ddC | - | - | + | GCAAUAAUCUAUACAAUACAACACAUACAAACAAAUUCUUAAGGUAAAAA/ddC | SRR25570243 |
| Dataset8 | R50CCC_ddC | - | - | + | GCAAUAAUCUAUACAAUACAACACAUACAAACAAAUUCUUAAGGUCCCAA/ddC | SRR25570242 |
| Dataset9 | R50GGG_ddC | - | - | + | GCAAUAAUCUAUACAAUACAACACAUACAAACAAAUUCUUAAGGUGGGAA/ddC | SRR25570241 |
| Dataset10 | R50UUU_ddC | - | - | + | GCAAUAAUCUAUACAAUACAACACAUACAAACAAAUUCUUAAGGUUUUAA/ddC | SRR25570240 |
| Dataset11 | R50CGC_ddC | - | - | + | GCAAUAAUCUAUACAAUACAACACAUACAAACAAAUUCUUAAGGUCGCAA/ddC | SRR25570274 |
| Dataset12 | R50AAA_ddC | - | + | + | GCAAUAAUCUAUACAAUACAACACAUACAAACAAAUUCUUAAGGUAAAAA/ddC | SRR25570273 |
| Dataset13 | R50CCC_ddC | - | + | + | GCAAUAAUCUAUACAAUACAACACAUACAAACAAAUUCUUAAGGUCCCAA/ddC | SRR25570272 |
| Dataset14 | R50GGG_ddC | - | + | + | GCAAUAAUCUAUACAAUACAACACAUACAAACAAAUUCUUAAGGUGGGAA/ddC | SRR25570271 |
| Dataset15 | R50UUU_ddC | - | + | + | GCAAUAAUCUAUACAAUACAACACAUACAAACAAAUUCUUAAGGUUUUAA/ddC | SRR25570270 |
| Dataset16 | R50CGC_ddC | - | + | + | GCAAUAAUCUAUACAAUACAACACAUACAAACAAAUUCUUAAGGUCGCAA/ddC | SRR25570269 |
| Dataset17 | R50AAA_ddC | $dT_2$ | - | + | GCAAUAAUCUAUACAAUACAACACAUACAAACAAAUUCUUAAGGUAAAAA/ddC | SRR25570268 |
| Dataset18 | R50CCC_ddC | $dG_2$ | - | + | GCAAUAAUCUAUACAAUACAACACAUACAAACAAAUUCUUAAGGUCCCAA/ddC | SRR25570267 |
| Dataset19 | R50GGG_ddC | $dC_2$ | - | + | GCAAUAAUCUAUACAAUACAACACAUACAAACAAAUUCUUAAGGUGGGAA/ddC | SRR25570266 |
| Dataset20 | R50UUU_ddC | $dA_2$ | - | + | GCAAUAAUCUAUACAAUACAACACAUACAAACAAAUUCUUAAGGUUUUAA/ddC | SRR25570265 |
| Dataset21 | R50CGC_ddC | $dT_2$ | - | + | GCAAUAAUCUAUACAAUACAACACAUACAAACAAAUUCUUAAGGUCGCAA/ddC | SRR25570263 |
| Dataset22 | R50CCC | - | - | - | GCAAUAAUCUAUACAAUACAACACAUACAAACAAAUUCUUAAGGUCCCAA | SRR25570262 |
| Dataset23 | R50CCC | - | - | + | GCAAUAAUCUAUACAAUACAACACAUACAAACAAAUUCUUAAGGUCCCAA | SRR25570261 |
| Dataset24 | R50CCC | - | + | - | GCAAUAAUCUAUACAAUACAACACAUACAAACAAAUUCUUAAGGUCCCAA | SRR25570260 |
| Dataset25 | R50CCC | - | + | + | GCAAUAAUCUAUACAAUACAACACAUACAAACAAAUUCUUAAGGUCCCAA | SRR25570259 |
| Dataset26 | R29CCCdA4_ddC | - | - | + | AACACAUACAAACAAAUUCUUAAGGUCCCAAAAAA/ddC | SRR25570258 |
| Dataset27 | R30CCCdA4_ddC | - | - | + | CAACACAUACAAACAAAUUCUUAAGGUCCCAAAAAA/ddC | SRR25570257 |
| Dataset28 | R31CCCdA4_ddC | - | - | + | ACAACACAUACAAACAAAUUCUUAAGGUCCCAAAAAA/ddC | SRR25570256 |
| Dataset29 | R32CCCdA4_ddC | - | - | + | UACAACACAUACAAACAAAUUCUUAAGGUCCCAAAAAA/ddC | SRR25570255 |
| Dataset30 | R31CCCdA3_ddC | - | - | + | ACAACACAUACAAACAAAUUCUUAAGGUCCCAAAAA/ddC | SRR25570254 |
| Dataset31 | R32CCCdA2_ddC | - | - | + | UACAACACAUACAAACAAAUUCUUAAGGUCCCAAAA/ddC | SRR25570252 |
| Dataset32 | R29CCCdA4_ddC | - | + | + | AACACAUACAAACAAAUUCUUAAGGUCCCAAAAAA/ddC | SRR25570251 |
| Dataset33 | R30CCCdA4_ddC | - | + | + | CAACACAUACAAACAAAUUCUUAAGGUCCCAAAAAA/ddC | SRR25570250 |
| Dataset34 | R31CCCdA4d_dC | - | + | + | ACAACACAUACAAACAAAUUCUUAAGGUCCCAAAAAA/ddC | SRR25570249 |
| Dataset35 | R32CCCdA4_ddC | - | + | + | UACAACACAUACAAACAAAUUCUUAAGGUCCCAAAAAA/ddC | SRR25570248 |
| Dataset36 | R31CCCdA3_ddC | - | + | + | ACAACACAUACAAACAAAUUCUUAAGGUCCCAAAAA/ddC | SRR25570247 |
| Dataset37 | R32CCCdA2_ddC | - | + | + | UACAACACAUACAAACAAAUUCUUAAGGUCCCAAAA/ddC | SRR25570246 |

| | MmRT-Cas1 | | FsRT-Cas1 | VvRT-Cas1 | Tt-Cas1 | Se-Cas1 | St-Cas1 |
|---|---|---|---|---|---|---|---|
| Cas system | WT | RTD | WT | WT | WT | WT | WT |
| Origin organism | Marinomonas mediterranea | | Fusicatenibacter saccharivorans | Vibrio vulnificus | Thermus thermophilus | Staphylococcus epidermidis | Streptococcus thermophilus |
| Assay organism | Marinomonas mediterranea | | Escherichia coli | Escherichia coli | Thermus thermophilus | Staphylococcus aureus | Streptococcus thermophilus |
| Assay organism strain | MMB-1 | | BL21-Gold(DE3) pLysS AG | HMS174(DE3) | HB27c | RN4220 | JIM 8232 |
| Target genome | MMB-1 | | BL21-Gold(DE3) pLysS AG | HMS174(DE3) | Phage phiFa/Ko | RN4220 | JIM 8232 |
| Target genome ID | NC_015276.1 | | NC_012947.1 | NZ_LM993812.1 | MH673671.2,MH673672.2 | NZ_CP076105.1 | FR875178.1 |
| BioProject | PRJNA301768 | | PRJNA484149 | PRJNA539885 | PRJNA631468 | PRJNA769698 | PRJNA762861 |
| BioSample | SRR2913703-810 | SRR2913815-818 | SRR8102160, 163, 166-8, 170, 172-7, 194, 207 | SRR8962131-134,137,138 | SRR11818505-8,SRR12227011-4 | SRR16249482-3 | SRR1595171-3, 176,187,198,209 |
| Spacers (unique) | 15,498 | 7,678 | 22,245 | 356 | 6,116 | 67,607 | 178,486 |
| None | 4,809 | 5,192 | 6,464 | 295 | 1,208 | 63,824 | 131,867 |
| 5'and3' ends | 279 | 117 | 2,693 | 4 | 813 | 130 | 2,967 |
| 5' end only | 4,933 | 1,255 | 5,509 | 15 | 2,771 | 1,861 | 21,912 |
| 3' end only | 5,477 | 1,114 | 7,579 | 42 | 1,324 | 1,792 | 21,740 |
| Spacers with soft clip (only in one end, 5' or 3', not both) | 10,410 | 2,369 | 13,088 | 57 | 4,095 | 3,653 | 43,652 |
| 1nt | 3,089 | 1,075 | 1,787 | 18 | 3,388 | 2,798 | 20,221 |
| ≥2nt | 7,321 | 1,294 | 11,301 | 39 | 707 | 855 | 23,431 |
